# Multimodal Hypersampling of the adult human heart resolves tissue-specific cell states and disease-associated variants

**DOI:** 10.64898/2026.06.16.732617

**Authors:** James Cranley, Lukas Mach, Takahiro Jimba, Kazumasa Kanemaru, Jing Zhao, Shani Perera, Yasemin-Xiomara Zurke, Simon Koplev, Sam Barnett, Lu Wang, Kenny Roberts, Rakeshlal Kapuge, Laura Richardson, Krzysztof Polanski, Claudia Semprich, Siew Yen Ho, Jan Lukas Robertus, Monika Madrova, Bulgantamir Tegshjargal, Antonio M.A. Miranda, Ioannis Sarropoulos, Ana-Maria Cujba, Woochan Lee, Anna Wilbrey-Clark, Jack Palmer, Joseph Brunet, Rasheda Chowdhury, Sanjay Prasad, John Dark, Helle Jørgensen, Michela Noseda, Sarah A. Teichmann

**Affiliations:** Cambridge Stem Cell Institute, University of Cambridge, Cambridge, UK; National Heart and Lung Institute, Imperial College London, London, UK; Wellcome Sanger Institute, Wellcome Genome Campus, Hinxton, UK; Victor Phillip Dahdaleh Heart & Lung Research Institute, University of Cambridge, Cambridge, UK; Department of Mechanical Engineering, University College London, London, UK; ESRF – The European Synchrotron, Grenoble, France; CIFAR Macmillan Multi-scale Human Programme, CIFAR, Toronto, Canada; Department of Medicine, University of Cambridge, Cambridge, UK; Department of Gene Technology, KTH Royal Institute of Technology, Science for Life Laboratory, Stockholm, Sweden; Translational and Clinical Research Institute, Newcastle University, Newcastle upon Tyne, UK; Royal Papworth Hospital, Cambridge, UK; British Heart Foundation Centre of Research Excellence, Imperial College London, London, UK; Royal Brompton Hospital, London, UK; UCL Cancer Institute, University College London, London, UK; Institute of Clinical Sciences, Faculty of Medicine, Imperial College London, London, UK; MRC Laboratory of Medical Sciences, London, UK

## Abstract

Cardiovascular diseases often arise in anatomically specialised regions of the heart that remain poorly represented in existing single-cell and spatial atlas references. Here we profiled 28 regions of the adult human heart from 36 donors, combining paired single-nucleus RNA and chromatin accessibility assays with spatial transcriptomics. The resulting atlas integrates 1,059,175 expression profiles and 452,614 chromatin accessibility profiles, and defines spatial niches across the heart. In cardiac valves, we identify a valve fibroblast–macrophage niche that is polarised to the non-fibrosa surface and expands with age. Aortic stenosis genetic risk was enriched in inflammatory regulatory programmes of the Valve Fibroblast Immune cell state. In pulmonary veins, we define *PITX2*^+^ Myocardial Sleeve Cells, a region-restricted cardiomyocyte population in which a fine-mapped 4q25 atrial fibrillation variant intersects a cell type-specific open chromatin region. In coronary arteries, we resolve a *RUNX1*-marked synthetic smooth muscle layer in the subintima that provides a molecular identity for diffuse intimal thickening, a constitutive feature of the healthy vessel wall and a substrate for atherosclerosis, and show by perturbation in human smooth muscle cells that RUNX1 promotes cell-cycle entry while restraining inflammatory cytokine gene expression. Finally, allele-specific accessibility together with deep learning models fine-tuned to predict regulatory activity from DNA sequence identify cell type-specific regulatory effects at GWAS loci for atrial fibrillation, coronary artery disease and calcific aortic valve stenosis, including a variant whose Valve Fibroblast-specific effect on the polyamine transporter *ATP13A3* is opposite in direction to the effect seen in bulk-tissue eQTL references that lack valve cells. Together with these cardiac sequence models and CardioSleuth, an AI-powered exploration engine for cardiac regulatory effects, this atlas links cardiac anatomy, gene regulation and disease genetics at single-cell resolution. By placing genetic risk in its precise cellular and anatomical context, it provides a foundation for mechanistic disease understanding, cell-state-specific therapeutic target nomination, and improved interpretation of inherited cardiovascular risk to inform future therapy and management.

## Introduction

The adult human heart is organised into anatomically specialised territories whose local cellular composition underlies regional function and disease susceptibility. Single-cell and single-nucleus atlases have established foundational catalogues of cardiac cell types across the major chambers ^1–3^. However, many clinically important cardiovascular diseases arise in structures that are under-represented in these references: valvular disease in the valve leaflets, atrial fibrillation in the pulmonary veins, and atherosclerosis in the coronary arteries. Because these regions are largely absent from atlases built around the major working-myocardial chambers, the cell states and niches that drive these prevalent conditions remain incompletely defined - as do the mechanisms through which they develop, including inflammation, fibrosis, remodelling and arrhythmogenesis. A multimodal map of these disease-relevant regions therefore creates an opportunity to identify the specific cell states and pathways that could serve as therapeutic targets.

This anatomical gap also limits interpretation of human genetics. Genome-wide association studies have identified numerous loci for cardiovascular traits, including coronary artery disease and atrial fibrillation ^4;5^, but most associated variants are non-coding and likely act through cell type-specific regulatory mechanisms. Bulk tissue datasets can obscure these effects, particularly when the relevant cell type is rare, spatially restricted or absent from the tissue that was sampled. Defining the cellular context of genetic risk therefore requires paired maps of gene expression, chromatin accessibility and spatial organisation across disease-relevant regions of the human heart.

Here we profile 28 anatomical regions of the adult human heart using paired single-nucleus RNA-seq and ATAC-seq, together with complementary spatial transcriptomic and imaging modalities. We detect a valve fibroblast–macrophage niche linked to aortic stenosis susceptibility, pulmonary vein myocardial sleeve cardiomyocytes connected to the *PITX2*/4q25 atrial fibrillation locus, and a synthetic smooth muscle cell state defining the coronary subintima, a potential substrate for atherosclerosis. We further integrate cell state-resolved chromatin accessibility with allele-specific accessibility and fine-tuned sequence-to-function models to prioritise cell type-specific genetic mechanisms. Together, these data provide an anatomical and regulatory reference for interpreting cardiovascular disease biology at single-cell resolution.

## Results

### Multimodal profiling of disease-relevant cardiac regions resolves regionally enriched cell states and niches

Many of the most prevalent cardiovascular diseases originate in anatomically specialised structures that are absent or only sparsely represented in existing single-cell and spatial atlases of the human heart ^1–3^. To resolve the cell states and tissue niches of these disease-relevant territories, we profiled 28 anatomical regions of the adult human heart, deliberately targeting sites that prior references have under-sampled, including the aortic and mitral valves, the pulmonary veins, the coronary and great (pulmonary and aortic) arteries, as well as multiple myocardial sampling sites (**Fig. 1a**; **Supplementary Table**). This sample set extends well beyond the major-chamber coverage of previous human heart atlases and captures region-restricted populations that were poorly represented in earlier datasets. For downstream analysis the 28 regions were organised into a hierarchical taxonomy (‘region_coarse’, 12 categories; ‘tissue’, 4 categories).

**Figure 1:**
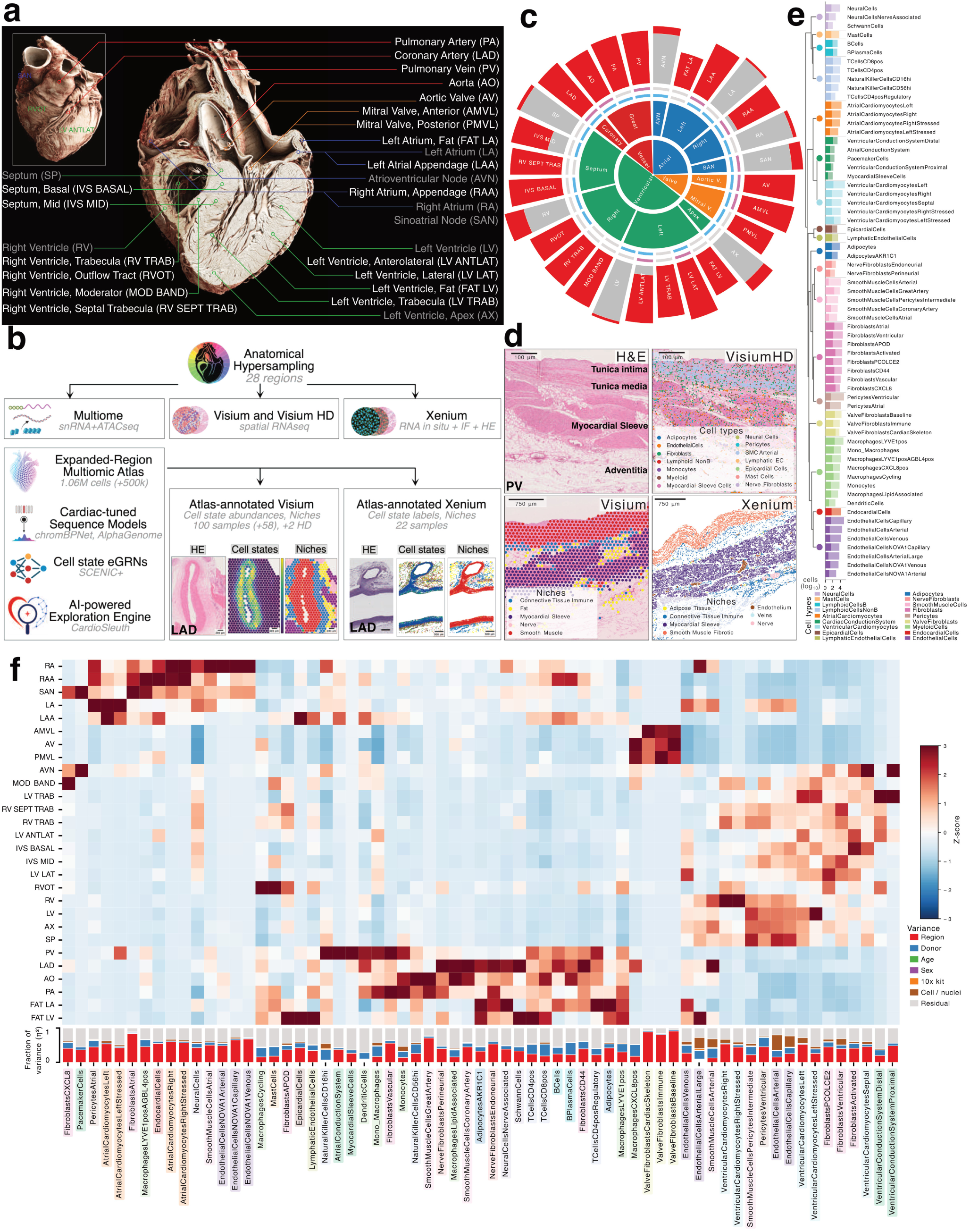
A multi-region, multi-modal atlas of the adult human heart. **a,** Overview of the 28-region sampling plan, depicted on a hierarchical phase-contrast tomography (HiP-CT) scan of a human heart at 8 *µ*m resolution. Line colours indicate four tissue categories (see also, c); grey labels denote regions included in Kanemaru *et al.* ^3^, white labels denote newly sampled regions. The HiP-CT image is for visualisation only and was not profiled in this study. **b,** Schematic of the data generation workflow. After multi-region dissection, samples were processed for single-nucleus RNA-seq and ATAC-seq (10x Multiome), spatial transcriptomics (10x Visium and Visium HD), and *in situ* gene profiling (10x Xenium) with post-Xenium immunofluorescence and H&E imaging. Single-cell data were integrated to form a 28-region RNA-seq and ATAC-seq atlas. This was used as a reference to annotate cell types and states in spatial gene expression data. Downstream resources include fine-tuned DNA sequence-to-function models, inferred gene regulatory networks, and CardioSleuth, an AI-powered exploration engine for cardiac regulatory effects. **c,** Distribution of data across 28 anatomical regions. The outer rim reflects the log_2_ number of cells per region; red indicates cells newly generated in this study, grey indicates previously published data ^3^. Pink segments indicate Xenium coverage, blue segments indicate Visium coverage. Inner segments reflect the ‘region coarse’ and ‘tissue’ level taxonomy. **d,** Multi-modal spatial profiling of the pulmonary vein. Top left: H&E histology showing the layered tissue architecture. Top right: Visium HD with spots coloured by cell type. Bottom left: Visium with spots coloured by niche identity. Bottom right: Xenium with cells coloured by NicheCompass-derived niche identity. **e,** Dendrogram linking 18 annotated cell types by transcriptional similarity. Leaves represent cell states. Bar plot shows per-cell state cell numbers, coloured by parent cell type. Bar is divided to depict the fraction of Multiome nuclei (full opacity) vs unpaired transcriptomes (partly-transparent). **f,** Regional cell state composition and variance partitioning. Top: heatmap of *z*-scored cell state proportions across 28 anatomical regions (red, enrichment; blue, depletion; both axes hierarchically clustered). Bottom: stacked bar chart showing the fraction of variance (*η*^2^) in each cell state’s proportion attributable to anatomical region, donor, age, sex, 10x kit version, cell/nuclei input types, and residual variance. Cell state labels highlighted according to their parent cell type identity (see E for legend).

From each region, paired single-nucleus RNA-seq and ATAC-seq data were generated using the 10x Genomics Multiome assay and integrated with our previously published dataset ^3^. After quality control, the single-cell atlas comprised 1,059,175 expression profiles, including 501,613 newly generated profiles, together with 452,614 nuclei for chromatin accessibility analysis (**Fig. 1b**). These data were complemented by spatial gene expression profiling: Visium (55 *µ*m; 100 samples, of which 58 were new), Xenium (377-gene panel; 19 samples), Xenium 5K (5,001-gene panel; 3 samples), and Visium HD (2 *µ*m; 2 samples), with several regions being profiled with multiple technologies (**Fig. 1b**). For Xenium experiments, post-Xenium protein immunofluorescence (using either 16-plex RareCyte or a 4-plex panel of decorin, collagen I, cardiac troponin and DAPI for nuclear staining) and H&E images were co-registered. In total, the multimodal dataset comprises samples from 36 donors (19 female, 17 male), including four donors who contributed spatial data only, across 28 anatomical regions (**Fig. 1b,c; Supplementary Fig. S1a,b**). Profiling each disease-relevant region with several assays was a deliberate design choice: rather than relying on any single technology, we sought concordant evidence across orthogonal modalities, so that the core cell states and spatial niches described below are independently corroborated rather than dependent on the assumptions of any one platform.

### Cell type annotation

Integration of single-cell gene expression data accounted for batch variation attributable to donor identity, input type (cell or nucleus), and 10x Genomics chemistry version. Cell type annotations were generated in a latent space constructed using a conditionally invariant variational autoencoder (inVAE), with anatomical ‘tissue’ as the invariant key ^6^. To confirm annotation robustness, we assessed label recoverability in an independent scVI^7^ latent space using cross-validated *k*-nearest neighbours classification, achieving a median accuracy of 83.5%, confirming that assignments are robust to the choice of integration framework (**Supplementary Fig. S1c,d,e**). We annotated 18 cell types and 65 cell states, organised in a hierarchical taxonomy (**Fig. 1e**). All major cell types from the previous atlas were recapitulated (**Supplementary Fig. S1f**). With the expanded dataset, two taxonomic refinements emerged: the previously unified Mural Cell category resolved into distinct Pericyte and Smooth Muscle Cell populations, and Endocardial Cells formed a cluster separate from Endothelial Cells (**Supplementary Fig. S1g,h**). Throughout, when referring to specific cell state annotations we use capitalisation and the original word order, for example Valve Fibroblasts Immune.

### Spatial data annotation and niche identification

Visium data were deconvolved using cell2location ^8^ to estimate cell state abundances per spot, followed by integration using inVAE and unsupervised clustering, which resolved eight spatially coherent tissue niches: Ganglia, Nerve, Myocardium, Fat, Conduction System, Connective Tissue Immune, Smooth Muscle, and Valve Fibrosa (**Supplementary Fig. S1i,j,k**).

Individual cells in Xenium data were segmented using Baysor ^9^, and cell state labels were transferred from the annotated single-nucleus RNA-seq data using CellTypist ^10^ (**Supplementary Fig. S1i**). Cross-regional integration and clustering based on NicheCompass ^11^ embedding revealed 17 niches with distinct cellular compositions and regional enrichments (**Supplementary Fig. S1l,m,n**).

While some niches, such as nerve, were consistently identified across both Xenium and Visium datasets, others (including myocardium and smooth muscle) were further resolved into multiple Xenium niches, showing the greater tissue complexity that can be resolved with a single-cell resolution method (**Supplementary Fig. S1o**). The breadth of spatial modalities is illustrated by multi-modal profiling of the pulmonary vein, where H&E histology, Visium HD, Visium, and Xenium each provide complementary views of the layered tissue architecture, its cell types and multicellular niches (**Fig. 1d**).

### Regional specificity of cardiac cell populations

Sampling multiple anatomical regions enabled the identification of regionally-enriched cell populations. To systematically quantify sources of compositional variation, we performed variance partitioning on sample-level cell type proportions using a linear model that included anatomical, donor-related (age, sex), and technical covariates. Anatomic region was the dominant source of variance for the majority of cell states, with pronounced regional specificity observed for Valve Fibroblasts, left and right Atrial Cardiomyocytes, Smooth Muscle Cell populations, and the cardiac conduction system (**Fig. 1f**). To distinguish cell populations that are regionally-restricted, we additionally quantified the regional specificity of each cell type and state using a Gini index of mean abundance across regions. This confirmed a clear gradient from highly region-restricted populations (valve fibroblasts, the cardiac conduction system, smooth muscle cell subtypes, and atrial cardiomyocytes) to broadly distributed populations such as fibroblasts and immune cells, which were relatively ubiquitous, with comparatively little regional bias (**Supplementary Fig. S1p,q**).

### Cardiac valves harbour an immune-active stromal niche linked to disease susceptibility

Inclusion of aortic and mitral valves in our expanded sampling strategy enabled the identification of Valve Fibroblasts as a transcriptionally distinct stromal population (**Fig. 2a**). Valve Fibroblasts shared pan-fibroblast markers such as *DCN* and *GSN* with conventional myocardial fibroblasts but were distinguished by expression of extracellular matrix and matrix-associated genes including *COMP*, *FMOD*, *BGN*, and *LTBP2*, and the complement-protective chaperone *CLU* (**Fig. 2b; Supplementary Fig. S2a, b**) ^12;13^. They express genes encoding targets of valvulopathy-inducing drugs, including serotonin agonists as well as MAO inhibitors (**Supplementary Fig. S2c**) ^14;15^. Within Valve Fibroblasts, three transcriptionally distinct states were resolved: a generic Baseline state and two specialised states, Cardiac Skeleton and Immune, each additionally expressing distinct marker genes superimposed on this shared fibroblast identity (*COMP* and *LTBP2* for Cardiac Skeleton; *CXCL2* and *LIF* for Immune; **Fig. 2b**).

**Figure 2:**
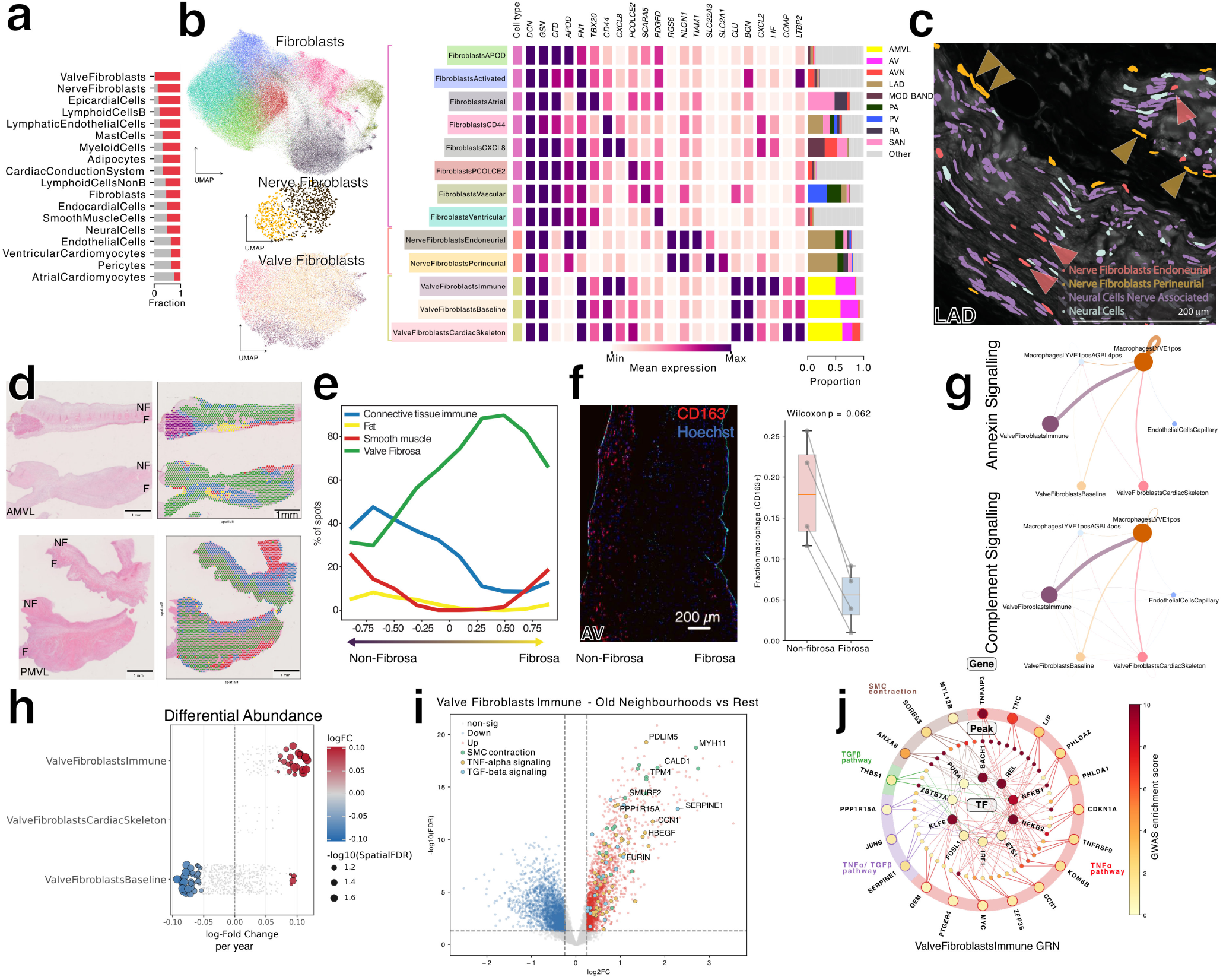
Cardiac valves harbour an immune-active stromal niche linked to disease susceptibility. **a,** Fraction of cells per cell type that are newly profiled in this atlas (red) versus present in the previous eight-region atlas (grey) ^3^. Valve Fibroblasts and Nerve Fibroblasts were largely absent from the earlier dataset. **b,** UMAP embeddings of the three fibroblast populations (Fibroblasts, Nerve Fibroblasts, and Valve Fibroblasts). Accompanying heatmap shows expression of core and marker genes, and the adjacent bar indicates the proportion of cells from each anatomical region per state (regions contributing at least 25% of any cell state shown). **c,** Xenium 5K spatial transcriptomic section of a left anterior descending (LAD) coronary artery-adjacent nerve bundle. Cell identity was defined by CellTypist label transfer from snRNA-seq. Nerve Fibroblasts Perineurial are detected at the surface of the nerve and Nerve Fibroblasts Endoneurial within the bundle interior, alongside Neural Cells and Neural Cells Nerve Associated. **d,** Left: H&E histology of mitral valve leaflet sections (anterior, AMVL; posterior, PMVL; fibrosa surface, F; non-fibrosa surface, NF). Right: spatial maps of transcriptional niches identified by unsupervised clustering of Visium data, coloured by niche identity. See panel e for colour legend. Myocardium niche is purple (not in legend). **e,** Line plot showing the percentage of Visium spots occupied by each niche as a function of position along the trans-valvular axis, from the flow-exposed surface (atrialis in mitral valve; ventricularis in aortic valve) to the fibrosa. The connective tissue–immune niche is enriched at the non-fibrosa surface. **f,** Left: RareCyte immunofluorescence of an aortic valve section stained for CD163 (macrophage marker) and Hoechst (nuclei). Right: Box plots comparing the fraction of CD163-positive cells (segmented nuclei with non-zero CD163 signal) in the fibrosa (bottom decile) versus non-fibrosa (top decile) of the trans-valvular axis (*n* = 4 sections; one-sided Wilcoxon signed-rank test, *p* = 0.062). **g,** CellChat-inferred cell–cell communication within the connective tissue–immune niche. Circle plots show signalling probabilities for the Complement pathway (left) and the Annexin pathway (right). Arrow width is proportional to communication probability; dot size indicates cell state abundance. **h,** Milo differential abundance analysis of valve fibroblast states as a function of donor age. Each dot represents a cellular neighbourhood; colour indicates log fold-change for each one-year increase in age (red, enriched in older donors; blue, enriched in younger donors). Valve Fibroblasts Immune show significant age-associated expansion. **i,** Volcano plot of differentially expressed genes in age-enriched versus remaining Valve Fibroblasts Immune neighbourhoods. Highlighted genes are coloured by pathway: TNF*α* signalling (red), TGF*β* signalling (blue), and smooth muscle contraction (green). **j,** Gene regulatory network of Valve Fibroblasts Immune visualised as a three-ring radial diagram coloured by GWAS *z*-score for aortic valve stenosis (SNP2CELL). The outer ring shows target genes grouped by pathway (TNF*α*, TGF*β*, smooth muscle cell phenotype), the middle ring shows enhancer peaks, and the inner ring shows upstream transcription factors. Curated gene sets were ranked by SNP2CELL score, with top genes retained alongside up to three corresponding enhancer peaks. TFs were selected from upstream TF–peak links, ranked by the number of selected peaks bound.

In addition to Valve Fibroblasts, Nerve Fibroblasts were captured, predominantly in coronary vessel-associated tissue (**Fig. 2b,c**). Profiling with Xenium resolved the characteristic spatial organisation of Nerve Fibroblasts Perineurial (expressing the organic cation transporter *SLC22A3* and *SLC2A1*, the latter encoding the GLUT1 transporter characteristic of the blood–nerve barrier) and Nerve Fibroblasts Endoneurial around and within nerve bundles (**Fig. 2b, c**) ^16^. Unlike other cardiac fibroblasts, they lack *TBX20* expression, which may suggest an extra-cardiac embryological origin (**Supplementary Fig. S2d**). Additionally, we observed Nerve Fibroblasts in neural ganglia alongside neuronal somata located near the coronary arteries and the sinoatrial node in both Visium and Xenium modalities (**Supplementary Fig. S2e-i**).

### A connective tissue-immune niche at the non-fibrosa valve surface

To determine the spatial organisation of valve fibroblast states, we analysed Visium spatial transcriptomics data from aortic and mitral valve leaflets. Unsupervised niche identification defined spatially coherent tissue domains including valve fibrosa, smooth muscle, fat, and a connective tissue-immune niche (**Fig. 2d; Supplementary Fig. S2j**). Analysis of niche abundance along a trans-valvular spatial axis, from the flow-exposed non-fibrosa surface (atrialis in mitral valve; ventricularis in aortic valve) to the fibrosa, revealed that the connective tissue-immune niche (harbouring Valve Fibroblasts Immune and macrophages) was preferentially localised to the non-fibrosa surface (**Fig. 2e; Supplementary Fig. S2k**). Post-Xenium RareCyte immunofluorescence showed a consistent trend at the protein level, with a higher fraction of CD163-positive macrophages in the non-fibrosa surface (*n* = 4 valve leaflet sections; Wilcoxon *p* = 0.062; **Fig. 2f; Supplementary Fig. S2l**). We also observed smooth muscle cells occupying layers near the valvular surfaces, notable since smooth muscle markers were previously described around calcified regions of stenotic aortic valves^17^ (**Fig. 2e; Supplementary Fig. S2m**).

### Complement and annexin signalling within the valve immune niche

Cell-cell communication analysis within the connective tissue-immune niche identified significant Complement and Annexin signalling interactions (**Fig. 2g; Supplementary Fig. S2n**). Valve Fibroblasts expressed the anti-inflammatory mediator *ANXA1* alongside complement component *C3* signalling to tissue-resident macrophages via receptors including *FPR1*, *FPR3*, *C3AR1*, and *ITGB2* (**Supplementary Fig. S2o, p**). This pairing suggests that valve fibroblasts may coordinate local complement signalling whilst concurrently dampening macrophage inflammatory activation via ANXA1/FPR interaction — a stromal immune regulatory role described in other tissue contexts ^18^. Among the three valve fibroblast states, Valve Fibroblasts Immune showed the highest expression of *C3*, suggesting a heightened capacity for local complement activation consistent with their pro-inflammatory transcriptional profile (**Supplementary Fig. S2p**).

### Age-associated expansion and inflammatory activation of valve fibroblasts

Differential abundance analysis of valve fibroblast states across the donor age range (from 20 to 74) revealed that the Valve Fibroblasts Immune state was significantly enriched in older donors (**Fig. 2h; Supplementary Fig. S2q, r**). Within Valve Fibroblasts Immune, neighbourhoods enriched for cells from older donors showed upregulation of TNF*α* and TGF*β* signalling, as well as smooth muscle contraction genes, relative to the remaining neighbourhoods, consistent with inflammatory and myofibroblast-like transcriptional features (**Fig. 2i; Supplementary Fig. S2s**). In parallel, older age-associated changes in Valve Fibroblasts Baseline showed enrichment of TNF*α*, chemokine, and complement pathways, supporting activation of complement signalling during ageing (**Supplementary Fig. S2t,u**). Together, these findings suggest that ageing is accompanied by both an expansion of Valve Fibroblasts Immune and increased pro-inflammatory, myofibroblast-like gene expression within this state, potentially disrupting fibroblast–macrophage crosstalk and predisposing the valve to disease.

### Genetic risk converges on Valve Fibroblasts Immune

To link these cellular states to disease susceptibility, we integrated cell state-specific gene regulatory networks (derived from SCENIC+) with GWAS summary statistics for aortic valve stenosis using SNP2CELL^19^. Disease-associated genetic signals were preferentially enriched in the regulatory networks of Valve Fibroblasts Immune, converging on an inflammation- and stress-responsive transcription factor programme centred on NF-*κ*B, IRF, and AP-1 signalling (**Fig. 2j**) ^20;21^. These results identify age-expanded Valve Fibroblasts Immune, situated within a spatially defined immune niche, as candidate mediators of genetically driven valve disease.

### Pulmonary vein Myocardial Sleeve Cells represent a specialised cardiomyocyte population

Within the single-nucleus RNA-seq data, we identified 14 cardiomyocyte cell states across the cardiac regions exhibiting atrioventricular and left–right heterogeneity (**Fig. 3a**), including specific drug target enrichments (Supplementary Fig. S3**a, b, c**). Beyond the expected atrioventricular division, atrial cardiomyocytes segregated along a left–right axis that mirrors developmental laterality ^22^. Atrial Cardiomyocytes Right were distinguished by *BMP10*, a heart-restricted TGF*β*-superfamily ligand that circulates as a biomarker of atrial fibrillation ^23;24^, and by *HAMP*, the iron-regulatory hormone hepcidin previously validated by smFISH as a right-atrial cardiomyocyte marker in the human heart ^1^. Atrial Cardiomyocytes Left reciprocally expressed *PANCR*, a left-atrial-restricted long non-coding RNA that positively regulates *PITX2c* expression ^25^ (**Fig. 3a; Supplementary Fig. S3c**). SCENIC+ enhancer-driven regulon activity in ventricular and atrial cardiomyocyte metacells reinforced these regional differences, with *HAND2* activity delineating right ventricular cardiomyocytes (Supplementary Fig. S3d, e). Further, we define stressed cardiomyocyte phenotypes within each chamber-enriched population. In the atria, the stressed populations are defined by *NPPB* and *FHL1* expression suggesting chronic stretch induced stress rather than processing-induced injury ^26;27^. Amongst the ventricular cardiomyocytes, the stressed populations were marked by *CCN1*, a gene induced by mechanical and ischaemic injury ^28^, alongside *XIRP1* and *XIRP2*, the latter associated with angiotensin II mediated stress ^29^.

**Figure 3:**
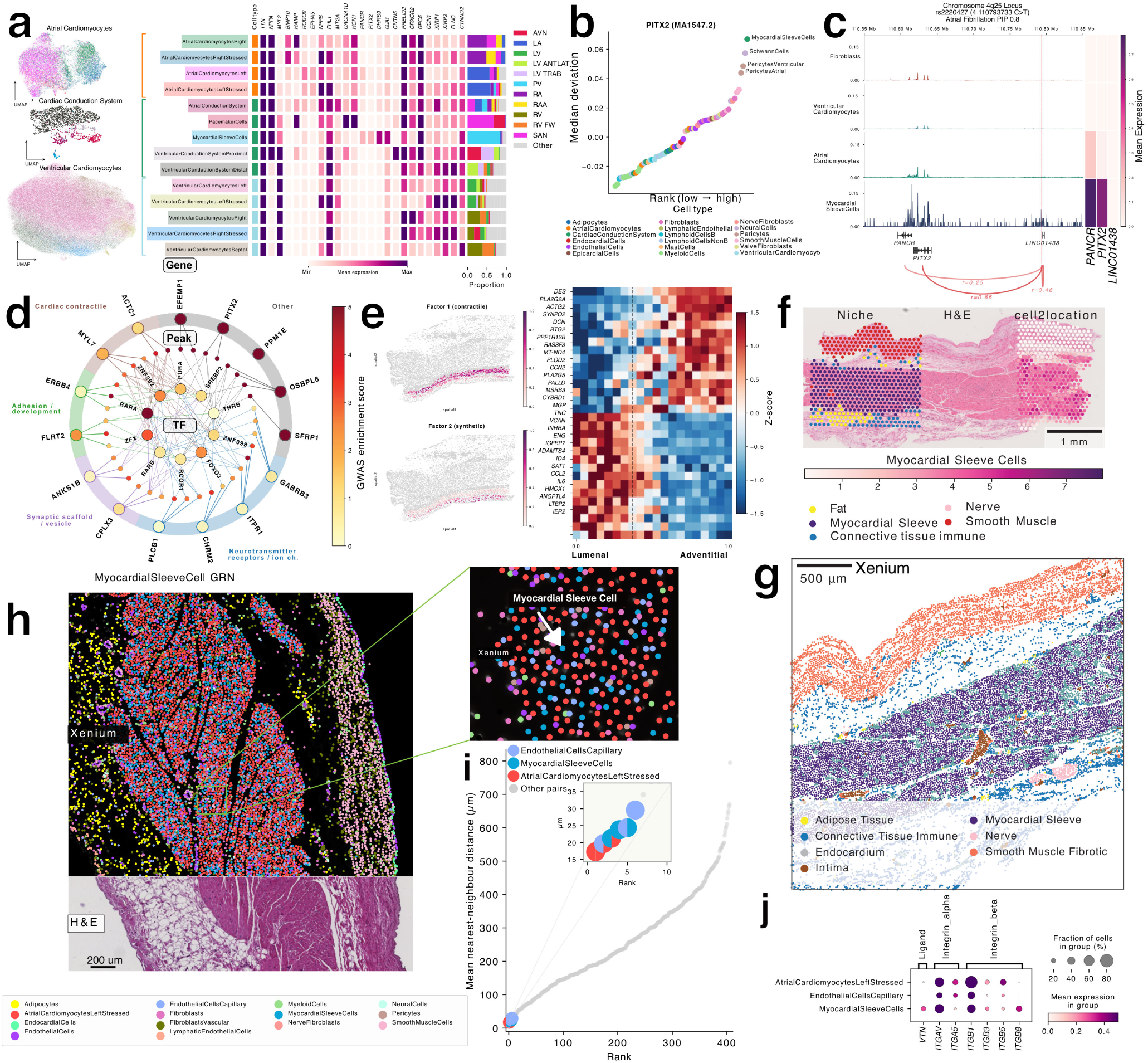
Pulmonary vein Myocardial Sleeve Cells represent a specialised cardiomyocyte population. **a,** UMAP of cardiomyocytes (atrial, ventricular and conduction system) coloured by cell state identity. Accompanying heatmap shows expression of core and marker genes, and the adjacent bar plots indicate the proportion of cells from each anatomical region per state (regions contributing at least 25% of any cell state shown). **b,** chromVAR motif enrichment for the PITX2 binding motif across all cardiac cell states. Myocardial Sleeve Cells showed the highest enrichment. **c,** Locus plot at the 4q25 atrial fibrillation locus showing fine-mapped credible-set SNPs, cell state-resolved chromatin accessibility tracks, and peak-to-gene linkages connecting the credible-set SNP (chr4:110,793,733 C*>*T, rs2220427; PIP = 0.80) to *PITX2* (*r* = 0.65), *PANCR* (*r* = 0.25), and *LINC01438* (*r* = 0.46). Heatmap shows expression of linked genes across cardiomyocyte cell states. **d,** Gene regulatory network of Myocardial Sleeve Cells visualised as a three-ring radial diagram coloured by GWAS *z*-score for atrial fibrillation (SNP2CELL). Curated gene sets labelled by sleeve cell-associated pathways were ranked by SNP2CELL score; top genes were retained with up to three corresponding enhancer peaks. TFs were selected from upstream TF-peak links, ranked by number of selected peaks bound. **e,** Visium HD analysis of pulmonary vein smooth muscle. Bin2cell-segmented smooth muscle cells were sub-clustered using NMF (*k* = 2), resolving two concentrically arranged transcriptional programmes (NMF factors 1 and 2): a synthetic programme (inner, luminal) and a contractile programme (outer, adventitial). Heatmap show expression of genes loading onto factors 1 and 2, ordered from the inner/luminal to the outer/adventitial layer. **f,** H&E staining of representative pulmonary vein (PV) sections showing smooth muscle (SM), myocardial sleeve (MS), and adventitia (Adv) (middle). Visium spatial niche annotations projected onto the same PV section (left). cell2location prediction scores for Myocardial Sleeve Cells displayed on the same section (right). **g,** Xenium spatial map of pulmonary vein tissue with cells coloured by NicheCompass-derived niche identity. **h,** Co-registered Xenium and H&E image of the pulmonary vein myocardial sleeve, with segmented cells coloured by cell state. Myocardial Sleeve Cells are intermingled with Atrial Cardiomyocytes Left Stressed and Endothelial Cells Capillary within the sleeve. **i,** Mean nearest-neighbour distance between cell states in the Myocardial Sleeve Xenium niche. **j,** Expression of the ligand (*VTN*) and receptors in Vitronectin signalling in snRNA-seq data of pulmonary veins.

The human pulmonary veins are a major source of ectopic electrical activity in atrial fibrillation ^30^ but have not previously been profiled at single-cell resolution. We generated Multiome, Visium, Visium HD, and Xenium data from this region, providing a comprehensive multi-modal characterisation of pulmonary vein tissue architecture (**Fig. 3a,e,f,g,h**).

A novel population, Myocardial Sleeve Cells, emerged almost exclusively from pulmonary vein samples (**Fig. 3a; Supplementary Fig. S3c**). Comparison with other cardiomyocyte and conduction system populations revealed a distinctive transcriptional profile. Myocardial Sleeve Cells were marked by expression of *PITX2*, a paired-like homeodomain transcription factor whose genomic locus (4q25) harbours the strongest known GWAS signal for atrial fibrillation ^31^; *DHRS9*, a retinol dehydrogenase involved in retinoic acid metabolism and the synthesis of ligands for RARA^32^ that has also been associated with atrial fibrillation ^33^; and *KCNMB2*, a regulatory subunit of large-conductance calcium-activated potassium (BK) channels (**Fig. 3a, d**) ^34^. Myocardial Sleeve Cells expressed the calcium-permeable cation channel *TRPM3*, previously reported as specific to Pacemaker Cells among cardiomyocyte states ^3^. Unlike Pacemaker Cells, however, they did not robustly express the hyperpolarisation-activated funny-current channels *HCN1* or *HCN4* (**Fig. S3c**). This raises the possibility that *TRPM3* contributes to membrane excitability in Myocardial Sleeve Cells, although a definitive cardiac role for *TRPM3* remains to be established.

Motif enrichment analysis of cell state-resolved ATAC-seq data using chromVAR confirmed that Myocardial Sleeve Cells exhibited the highest chromatin accessibility enrichment for PITX2 binding motifs among all cardiac cell types (**Fig. 3b**). This profile, combining arrhythmia-associated gene expression with pulmonary vein restriction, positions Myocardial Sleeve Cells as candidate mediators of ectopic electrical activity (Supplementary Fig. S3**c**).

Drug target enrichment analysis using drug2cell highlighted several compounds with predicted cell state-specific effects on Myocardial Sleeve Cells, including doxapram and inhalation anaesthetics targeting the TASK-1 potassium channel (encoded by *KCNK3*), potentially relating to the differential risk of post-surgical atrial fibrillation with inhalation anaesthetics versus propofol and to the anti-arrhythmic effects of doxapram (Supplementary Fig. S3f, k-m) ^35–38^. Hotspot analysis identified a Myocardial Sleeve Cell-specific gene module enriched for neuronal and synaptic gene sets (synapse, neuron projection; GSEA), spanning neurotransmitter receptors and ion channels, synaptic and vesicle-related genes, and axon-guidance molecules (Supplementary Fig. S3g-j). The inclusion of the parasympathetic receptor *CHRM2* and the atrial serotonin receptor *HTR4* raises the possibility that this programme reflects autonomic responsiveness or close neural association of the sleeve myocardium, consistent with the dense innervation of the pulmonary veins.

### A candidate causal variant at the 4q25 atrial fibrillation locus

To investigate whether disease-associated genetic variation at this locus acts through Myocardial Sleeve Cells, we intersected fine-mapped credible-set SNPs from atrial fibrillation GWAS with cell state-resolved open chromatin regions. A 95% credible-set SNP (chr4:110,793,733 C*>*T; rs2220427; PIP = 0.80) overlapped an accessible chromatin peak selectively open in Myocardial Sleeve Cells. Peak-to-gene linkage analysis identified significant correlations between this peak and the expression of *PITX2* (*r* = 0.65), *PANCR* (*r* = 0.25), and the long non-coding RNA *LINC01438* (*r* = 0.46) (**Fig. 3c**). SNP2CELL network propagation of atrial fibrillation genetic risk implicated cardiomyocytes broadly, but with the highest scores in atrial and conduction-system subtypes including Pacemaker Cells and Myocardial Sleeve Cells (**Supplementary Fig. S3q**). Further analysis of enhancer-driven gene regulatory networks (eGRNs) showed that atrial fibrillation risk variants were enriched in a nuclear-receptor-centred regulatory programme in Myocardial Sleeve Cells, including the retinoic acid receptors RARA and RARB alongside metabolic and stress-associated regulators such as SREBF2 and FOXO3 (**Fig. 3d**). Retinoic acid signalling has been implicated in the regulation of *PITX2* and the specification of pulmonary vein and left atrial cardiomyocyte identity ^39^, consistent with the developmental origin of this population.

### Spatial organisation of the pulmonary vein wall

Histological and spatial transcriptomic analysis of pulmonary vein cross-sections revealed a layered architecture comprising smooth muscle, the myocardial sleeve, and adventitia (**Fig. 3e, f, g**). To characterise heterogeneity within the smooth muscle layer, we applied bin2cell segmentation to Visium HD data and performed non-negative matrix factorisation (NMF) on smooth muscle cells. This resolved two smooth muscle cell transcriptional programmes arranged concentrically: a synthetic programme (marked by *VCAN*, *TNC*, and *ENG*) enriched at the luminal surface, and a contractile programme (marked by *DES*, *PLA2G2A*, and *ACTG2*) enriched at the adventitial surface (**Fig. 3e**).

Xenium profiling with NicheCompass-based niche identification resolved distinct spatial domains within the pulmonary vein, with the myocardial sleeve niche clearly delineated from surrounding smooth muscle and connective tissue (**Fig. 3f, g, h**). Xenium data demonstrated that the myocardial sleeve is not composed solely of Myocardial Sleeve Cells, but comprises three principal cell states: Endothelial Cells Capillary, Myocardial Sleeve Cells, and Atrial Cardiomyocytes Left Stressed (**Fig. 3h, i; Supplementary Fig. S3n**). Cell-cell communication analysis within the sleeve niche identified Vitronectin signalling between Myocardial Sleeve Cells and neighbouring Atrial Cardiomyocytes Left Stressed and Endothelial Cells Capillary (**Fig. 3j; Supplementary Fig. S3o, p**).

### Vascular smooth muscle cell diversity across cardiac arteries

We annotated four vascular cell types (Smooth Muscle Cells, Pericytes, Endothelial Cells and Endocardial Cells), comprising 15 cell states in total (**Fig. 4a**). In accordance with a recent vascular cell atlas ^40^, Endothelial Cells could be resolved into cell states distinguished by vascular bed: Capillary (*RGCC*, *CA4*), Arterial (*HEY1*), Arterial Large (*GJA5*, *SULF1*) and Venous (*ACKR1*). We additionally identified arterial, capillary and venous states marked by the neuronal splicing regulator *NOVA1*. Each retained its conventional bed-identity markers while sharing a common programme (*NOVA1*, *FGF23*, *NR4A2*, *PDK4*, *PER1*, *RGL1*, *ACER2*; **Supplementary Fig. S4a**). *NOVA1* bearing endothelial cell states were largely restricted to right atrial and sinoatrial samples (**Fig. 1f**). Pericytes (co-expressing *ACTA2*, *ABCC9* and *CD36*) separated into Atrial and Ventricular states, the former marked by *SLIT3*, a secreted guidance glycoprotein implicated in cardiac development and fibrosis ^41^. Endocardial Cells formed a distinct cluster expressing *VWF* but distinguishable from Endothelial Cells by *CDH11*, *NRG3* and *PLA2G5*.

**Figure 4:**
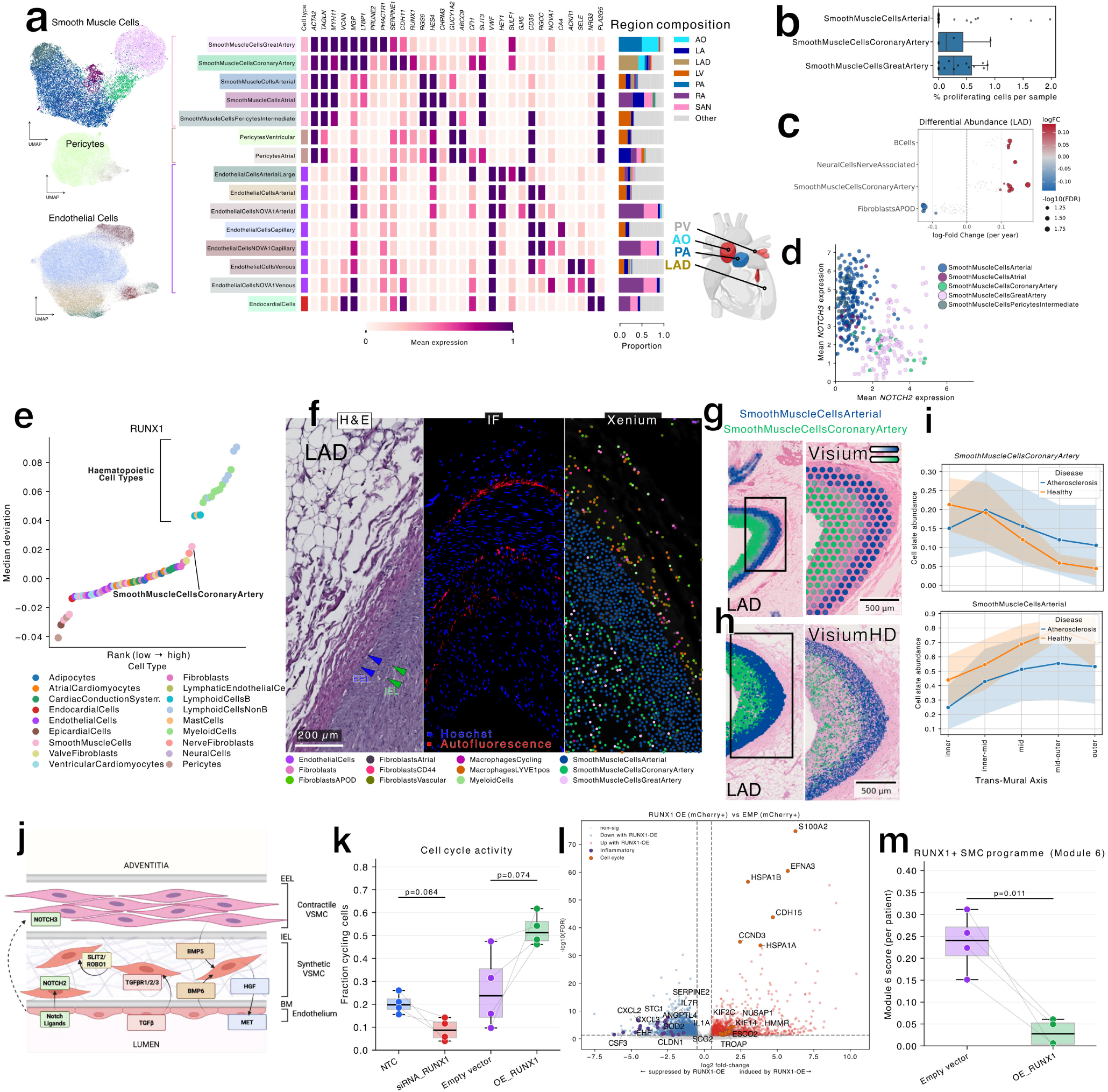
Cardiac arteries contain transcriptionally diverse, regionally specialised, and spatially zonated smooth muscle cell populations. **a,** UMAP embedding of vascular cell types (Smooth Muscle Cells, Pericytes, Endothelial Cells) coloured by cell state, resolving 15 states. Accompanying heatmap shows expression of core and marker genes, and the adjacent bar indicates the proportion of cells from each anatomical region per state (regions contributing at least 25% of any cell state shown). Endocardial Cells UMAP not shown but their gene expression profile is included in the heatmap. **b,** Proportion of cycling cells across smooth muscle cell states in single-nucleus RNA-seq data. Cells were classified as cycling on the basis of non-zero expression of at least one proliferation marker (*MKI67*, *TOP2A* or *CDK1*). Each point is one sample (sample–cell state combinations with fewer than 200 cells excluded); boxes show the median and interquartile range, with whiskers extending to 1.5× the interquartile range. Synthetic populations (Smooth Muscle Cells Coronary Artery and Smooth Muscle Cells Great Artery) showed a higher cycling fraction than contractile populations. **c,** Differential cell state abundance analysis of coronary artery (LAD) samples as a function of donor age. Beeswarm plot of per-neighbourhood log-fold change in abundance per year of age, grouped by cell state; positive values indicate increasing abundance with age. Neighbourhoods with a spatial FDR below 0.1 were considered significantly age-associated and are coloured by log-fold change, with point size encoding − log_10_(spatial FDR); non-significant neighbourhoods are shown in grey. Smooth Muscle Cells Coronary Artery neighbourhoods showed increased abundance with age. **d,** Mean log-normalised *NOTCH2* versus *NOTCH3* expression per smooth muscle cell neighbourhood, coloured by dominant cell state. Synthetic states show higher *NOTCH2* relative to *NOTCH3*, with the converse in contractile states. **e,** Median chromVAR motif accessibility deviation for the RUNX1 motif across cardiac cell states, ranked from lowest to highest and coloured by parent cell type. Smooth Muscle Cells Coronary Artery showed the highest RUNX1 motif accessibility of any non-haematopoietic cell state. **f,** Co-registered split view of a human coronary artery cross-section: histology (H&E), immunofluorescence (IF) and Xenium spatial transcriptomics of the same tissue region. In the H&E panel, green and blue arrows mark the internal and external elastic laminae (IEL and EEL), respectively. The IF panel shows nuclei (Hoechst) and autofluorescence, the latter highlighting the elastic fibres of the IEL and EEL. The Xenium panel shows segmented cells coloured by cell state: the most abundant states within the region of interest are shown individually and remaining cells are collapsed to their parent cell type. **g,** Visium spatial profiling of a coronary artery cross-section, resolving the concentric transition from an inner Smooth Muscle Cells Coronary Artery-dominated layer to an outer Smooth Muscle Cells Arterial-dominated media. Values plotted are per-spot cell2location cell-type abundances. **h,** Visium HD spatial profiling of a coronary artery cross-section, corroborating the dual-layer smooth muscle cell architecture at higher spatial resolution. Bin2cell-segmented cells are coloured by cell state, showing Smooth Muscle Cells Coronary Artery and Smooth Muscle Cells Arterial only. **i,** Transmural axis analysis comparing smooth muscle cell state distribution in healthy coronary arteries (this study) and atherosclerotic coronary arteries ^48^. There was no significant difference in transmural distribution between conditions. **j,** Schematic of cell–cell interactions in the coronary artery wall, showing key signalling axes (BMP, NOTCH, SLIT–ROBO, TGF*β* and HGF–MET) between endothelial cells, contractile Smooth Muscle Cells Arterial of the media (NOTCH3) and synthetic Smooth Muscle Cells Coronary Artery of the subintima (NOTCH2). As Notch signalling normally occurs between cells in direct contact rather than via secreted factors, the dashed arrow denotes the potential for this interaction. **k,** Fraction of cycling cells per donor across perturbation conditions: non-targeting siRNA control (NTC), RUNX1 siRNA knockdown (siRNA_RUNX1), empty-vector lentiviral control (Empty vector) and RUNX1 lentiviral overexpression (OE_RUNX1). Cells were classified as cycling on the basis of S- or G2/M-phase assignment (Scanpy score_genes_cell_cycle with the cell-cycle gene set from Tirosh *et al.* ^53^); each point is one donor, with lines connecting paired donors and boxes showing median and interquartile range. *P* values, paired *t*-test across donors. **l,** Volcano plot of pseudobulk differential expression between RUNX1 overexpression (OE_RUNX1) and empty-vector control. Each point is a gene; coloured points pass the significance thresholds (Benjamini–Hochberg-adjusted *P <* 0.05 and | log_2_ fold-change| *>* 0.5), red for up- and blue for down-regulated with RUNX1 overexpression. Cell-cycle genes were upregulated and inflammatory cytokine genes downregulated. **m,** Per-donor activity of Module 6 (the Smooth Muscle Cells Coronary Artery *in vivo* gene programme containing *RUNX1* and inflammatory cytokine genes) across perturbation conditions. Each point is one donor, with lines connecting paired donors and boxes showing median and interquartile range; module scores were computed per cell with score_genes and averaged per donor. *P* values, paired *t*-test across donors. RUNX1 overexpression reduces expression of this gene module.

Cardiac smooth muscle cells exhibited strong regional specificity. By transcriptional signature and region of origin, contractile smooth muscle cell states (*RGS6*, *HES4*, *SLIT3*) were distinguished from synthetic cell states (also termed fibromyocytes; *LTBP1*, *VCAN*, *MGP*), the latter enriched for a glycosaminoglycan-biosynthetic programme (**Supplementary Fig. S4b,c,d**). Within the contractile group we identified a general Smooth Muscle Cells Arterial state, the predominant contractile population found across all sampled arteries and cardiac regions, including the great arteries and coronary arteries; a *GUCY1A2*^+^ Smooth Muscle Cells Pericytes Intermediate population; and a Smooth Muscle Cells Atrial state, the last highly enriched in atrial samples and marked by the muscarinic M3 receptor (*CHRM3*). This is consistent with the dense parasympathetic innervation of the atria ^42^, and drug2cell analysis predicted an atropine-responsive pharmacological profile for this population (**Supplementary Fig. S4e**).

Synthetic smooth muscle cell states showed a higher fraction of cycling cells than contractile states (**Fig. 4b**). They also showed reciprocal *NOTCH2*/*NOTCH3* expression, with higher *NOTCH2* relative to *NOTCH3* in synthetic populations and the converse in contractile states (**Fig. 4d**). Synthetic cells subclustered into two populations (**Fig. 4a**): Smooth Muscle Cells Coronary Artery, enriched in (but not specific to) coronary artery (LAD) samples, and Smooth Muscle Cells Great Artery, specific to the great arteries (AO and PA). The former were enriched in coronary samples from older donors (**Fig. 4c**) and were marked by *RUNX1*, *SERPINE1* (recently proposed as a foam-cell marker ^43^) and an inflammatory cytokine gene expression module. Smooth Muscle Cells Great Artery were marked by *MYH10*, *PHACTR1* and *PRUNE2*, together with a module enriched for extracellular-matrix genes including *ELN* and *COL3A1* (**Supplementary Fig. S4c,d**). Throughout, cell-state names indicate the vessel or region in which a state is enriched or specific, not the only smooth muscle cell population present there; contractile Smooth Muscle Cells Arterial were found in all arteries, including the great arteries alongside Smooth Muscle Cells Great Artery.

Among the markers distinguishing Smooth Muscle Cells Coronary Artery, the transcription factor *RUNX1* was notable. Although canonically associated with haematopoietic development ^44^, recent work has identified RUNX1 as a regulator of vascular smooth muscle cell phenotype ^45^. Motif enrichment analysis of cell-state-resolved ATAC-seq data using chromVAR ranked Smooth Muscle Cells Coronary Artery as the most enriched for the RUNX1 motif among all non-haematopoietic cell states (**Fig. 4e**).

### Spatial organisation and cell-cell communication in the healthy coronary artery wall

Spatial gene expression profiling using Xenium, Visium and Visium HD consistently revealed a concentric organisation of smooth muscle cell states across the coronary artery wall (**Fig. 4f,g,h; Supplementary Fig. S4f**). Triple co-registered Xenium imaging (gene expression, immunofluorescence and H&E) of coronary artery cross-sections localised contractile Smooth Muscle Cells Arterial to the tunica media, whereas the synthetic Smooth Muscle Cells Coronary Artery population occupied the subintimal layer between the endothelium and the internal elastic lamina, with a minority also present in the media (**Fig. 4f**). Visium and Visium HD confirmed this dual-layer architecture, resolving a concentric transition from an inner Smooth Muscle Cells Coronary Artery layer to an outer Smooth Muscle Cells Arterial media (**Fig. 4g,h**). This subintimal localisation of Smooth Muscle Cells Coronary Artery cells, together with the synthetic transcriptional phenotype, is consistent with the histological entity conventionally termed diffuse intimal thickening (DIT) — a common feature of healthy human coronary arteries that is considered a substrate for atherosclerotic plaque development ^46;47^. Our data therefore provide a molecular and cellular identity for coronary DIT, defining it as a layer of *RUNX1*-marked, coronary-enriched fibromyocytes.

To assess whether this transmural organisation was altered in established coronary artery disease, we transferred our cell state labels to published Visium data from atherosclerotic coronary arteries ^48^ and applied the same inner-to-outer axis analysis. Neither Smooth Muscle Cells Coronary Artery nor Smooth Muscle Cells Arterial showed a statistically significant difference in transmural distribution between healthy and atherosclerotic vessels (**Fig. 4i**), suggesting that Smooth Muscle Cells Coronary Artery represent a constitutive feature of normal coronary biology, rather than a consequence of disease.

Next, we performed cell–cell interaction analysis using CellPhoneDB ^49^ between smooth muscle and endothelial cell states in the coronary artery niche, where Endothelial Cells Arterial Large are the predominant endothelial state (**Supplementary Fig. S4h**). This revealed several potential signalling axes between the two compartments, including BMP, NOTCH, SLIT–ROBO, TGF*β* and HGF–MET (**Fig. 4j**; **Supplementary Fig. S4g**). HGF–MET signalling was predicted between Smooth Muscle Cells Coronary Artery (sender) and Endothelial Cells Arterial Large (receiver), with *HGF* expression largely restricted to Smooth Muscle Cells Coronary Artery (**Supplementary Fig. S4i**). As circulating HGF is an early biomarker of subclinical coronary atherosclerosis ^50;51^, our data raise the possibility that the synthetic smooth muscle cells of the coronary subintima are a source of this signal. Both Smooth Muscle Cells Coronary Artery and Smooth Muscle Cells Great Artery expressed multiple BMP receptors (*BMPR1A*, *BMPR1B*, *ACVR1* and *BMPR2*), rendering them competent to respond to *BMP5* from the contractile Smooth Muscle Cells Arterial of the media and *BMP6* from the overlying Endothelial Cells Arterial Large (**Supplementary Fig. S4i**). Loss-of-function mutations in *BMPR2* drive smooth muscle cell proliferation in pulmonary arterial hypertension ^52^; by analogy, intact BMP reception on coronary synthetic smooth muscle cells may represent a homeostatic brake on proliferation in the healthy vessel wall.

### RUNX1 perturbation confirms a proliferative and immunomodulatory role in vascular smooth muscle

To functionally test the role of RUNX1 in smooth muscle cell biology, we performed gain and loss of function perturbations in primary human smooth muscle cells using lentiviral overexpression and siRNA-mediated knockdown of RUNX1, followed by single-cell RNA sequencing (**Supplementary Fig. S4j,k,l,m**). Overexpression of RUNX1 promoted cell cycling, while knockdown reduced it, identifying RUNX1 as a driver of smooth muscle cell proliferation (**Fig. 4k**). Differential expression analysis revealed two principal transcriptional responses: upregulation of cell cycle genes, and downregulation of inflammatory cytokine programmes (**Fig. 4l; Supplementary Fig. S4n,o**). To separate effects that scaled with transgene dosage from those that did not, we modelled the expression of each differentially expressed gene against the mCherry reporter as a proxy for RUNX1 dose, controlling for donor. The proliferative response was dose-independent, with cell-cycle genes showing no significant relationship to transgene level, consistent with a largely threshold (switch-like) effect; in contrast, suppression of inflammatory cytokine expression covaried with mCherry, indicating a graded, dose-dependent response (**Supplementary Fig. S4p,q**). RUNX1 overexpression significantly reduced the activity of the gene module specific to Smooth Muscle Cells Coronary Artery *in vivo* (Module 6), which contains multiple inflammatory cytokine genes as well as *RUNX1* itself (**Fig. 4m; Supplementary Fig. S4r**). This suggests a negative-feedback architecture: RUNX1 is a component of the Smooth Muscle Cells Coronary Artery programme, and elevated RUNX1 activity drives proliferation while simultaneously dampening the inflammatory limb of that programme.

Taken together, these data position Smooth Muscle Cells Coronary Artery as a constitutive, age-accumulating synthetic smooth muscle population that constitutes the cellular identity of diffuse intimal thickening in the healthy coronary artery, maintained in a proliferative but inflammation-restrained state by RUNX1. Their expansion with age, competence for BMP-mediated proliferative control, and active suppression of inflammatory output are consistent with a homeostatic, potentially protective role; whether loss of this RUNX1-enforced restraint permits the same population to contribute to atherosclerotic plaque, of which diffuse intimal thickening is a recognised substrate, remains to be tested.

### Cell type-specific genetic effects shed light on common variant disease mechanisms

#### Genotype profiling and allele-specific accessibility using Multiome data

To identify variants likely to act through cell type-specific regulatory mechanisms, we developed an allele-specific accessibility (ASA) framework leveraging our single-nucleus Multiome dataset (**Fig. 5a**). We selected 655 GWAS SNPs with posterior inclusion probability (PIP) *>* 0.4 across three cardiovascular traits relevant to the anatomical regions profiled in our other analyses: atrial fibrillation (AF), coronary artery disease (CAD), and calcific aortic valve stenosis (CAVS). A total of 26 donors were genotyped at each SNP using array-based (*n* = 15) or read-based (RNA and ATAC; *n* = 11) approaches, with genotyping coverage at the majority of SNP sites (**Methods**). Read- and array-based genotypes were highly concordant (96.2%, 531/552 donor-SNP genotype calls, using combined RNA and ATAC reads; **Supplementary Fig. S5a**). To mitigate reference-mapping bias, we performed allelic remapping using WASP^54^ before UMI-deduplicated, per-barcode allele counting (**Supplementary Fig. S5b**).

**Figure 5:**
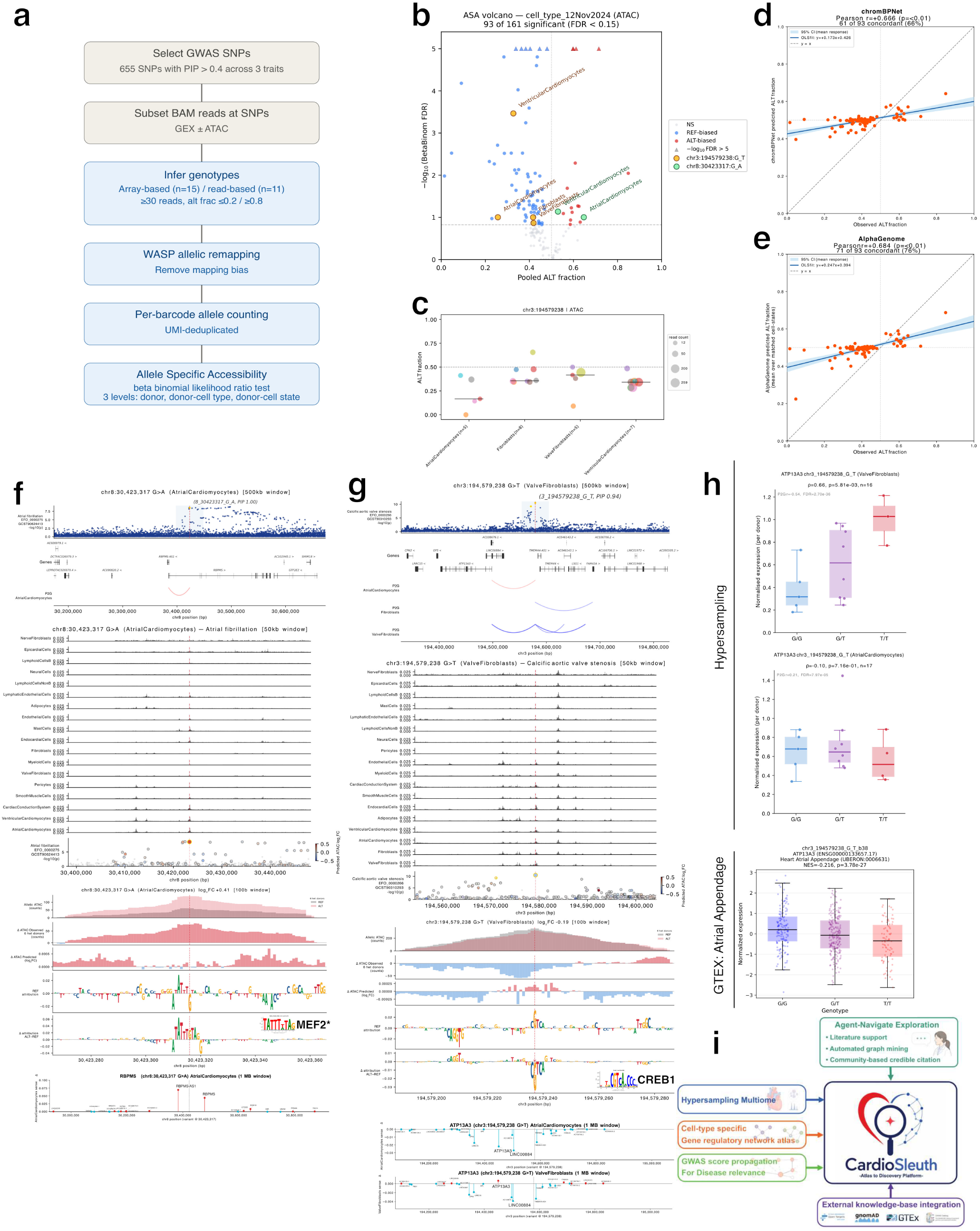
Cell type-specific genetic effects shed light on common variant disease mechanisms. **a,** Allele-specific accessibility (ASA) workflow. 655 GWAS SNPs (PIP *>* 0.4 across three cardiovascular traits) were genotyped using array-based (*n* = 15) or read-based (*n* = 11) approaches, followed by WASP allelic remapping ^54^, per-barcode allele counting (UMI-deduplicated for RNA reads, position-deduplicated for ATAC), and beta-binomial likelihood-ratio testing at donor (bulk, across all cell types), donor-cell type, and donor-cell state levels. **b,** ASA volcano plot at the donor-cell type level. The *x* axis shows the pooled ALT fraction, the fraction of reads carrying the alternate allele, pooled across donors for each SNP-cell type pair, and the *y* axis shows − log_10_ of the beta-binomial FDR, capped at 5. The vertical dotted line at 0.5 marks allelic balance; points left of it are REF-biased (blue) and points right are ALT-biased (red), with the horizontal dashed line at the FDR *<* 0.15 significance threshold. Of 161 testable SNP-cell type pairs, 93 were significant (FDR *<* 0.15). Highlighted are the calcific aortic valve stenosis (CAVS)-associated variant rs1706003 (chr3:194,579,238 G*>*T, yellow) and the atrial fibrillation-associated variant rs2979489 (chr8:30,423,317 G*>*A, green), each labelled by cell type. **c,** Bubble plot of per-donor ALT allele fraction for rs1706003 across the cell types with significant allelic imbalance (FDR *<* 0.15). Each point is one heterozygous donor; point size encodes the number of supporting reads and colour encodes donor identity. The black horizontal line marks the per-cell type median across donors and the dashed line at 0.5 marks allelic balance. The number of donors (n) and the beta-binomial FDR are shown for each cell type. **d,** Observed versus chromBPNet-predicted ALT allele fractions for the 93 significant ASA SNP-cell type pairs with a valid prediction. The dashed grey line is the identity (*y* = *x*); the blue line is an ordinary least-squares fit with its 95% confidence band, and dashed reference lines mark allelic balance (0.5) on each axis. Observed and predicted fractions are positively correlated (Pearson *r* = 0.67, *p* = 2.4 × 10^−13^), with 61 of 93 pairs (66%) concordant. **e,** As in **d**, for AlphaGenome predictions (Pearson *r* = +0.684, *p <* 0.01; 71 of 93 pairs (76%) concordant). **f,** Three-level dissection of the atrial-fibrillation-associated variant rs2979489 (chr8:30,423,317 G*>*A; PIP = 1) in Atrial Cardiomyocytes. **Top** (500 kb window, SNP marked by the red dashed line): GWAS association (− log_10_ *p*) for atrial fibrillation; variants in the 95% credible set are gold, all others blue. Below is a gene track (exons as filled blocks, introns as lines, strand shown by chevrons), and beneath it cell type-specific ArchR peak-to-gene (P2G) arcs from the Multiome data, linking the rs2979489-containing peak to candidate target genes; arc colour denotes the peak–gene correlation (red positive, blue negative). **Middle** (50 kb window): cell type-specific ATAC accessibility tracks, with a lower scatter of common variants (gnomAD v4.1) coloured by their chromBPNet-predicted accessibility effect (red increase, blue decrease; gold edge marks credible-set membership; grey points lack a prediction). **Bottom** (100 bp window): allelic ATAC pileup over donors heterozygous for rs2979489 (ALT in red, REF in grey, superimposed) with the observed ALT−REF difference below; the chromBPNet-predicted ALT−REF difference (log_2_FC); and base-resolution contribution scores for the REF allele followed by the ALT−REF difference (height above the axis indicates increased predicted accessibility, below decreased). rs2979489 has the strongest predicted accessibility effect within the 50 kb window and is predicted to create a MEF2-family motif (motif inset; *p* = 2.7 × 10^−7^). **Lowermost**: AlphaGenome-predicted gene-expression effects across a 1 Mb window centred on the variant. Genes with larger predicted effects are highlighted. **g,** As in **f**, for the calcific aortic valve stenosis-associated variant rs1706003 (chr3:194,579,238 G*>*T) in Valve Fibroblasts. The variant is associated with reduced accessibility, and attribution analysis indicates destruction of a motif most closely matching CREB1 (motif inset; *p* = 2.8 × 10^−6^). The peak-to-gene arcs show the divergent cell type-specific links to *ATP13A3* (positive in Atrial Cardiomyocytes, negative in Valve Fibroblasts). **Lowermost**: AlphaGenome-predicted gene-expression effects across a 1 Mb window in Valve Fibroblasts and Atrial Cardiomyocytes, predicting reduced expression of several genes in both cell types but increased *ATP13A3* expression specifically in Valve Fibroblasts. **h,** Genotype–expression associations for *ATP13A3* at rs1706003. **Top** and **middle**: per-donor mean expression versus ALT allele dosage from the Multiome data, showing a significant positive association in Valve Fibroblasts (*ρ* = 0.66, *p* = 5.81 × 10^−3^, *n* = 16) and a non-significant trend toward reduced expression in Atrial Cardiomyocytes (*ρ* = −0.10, *p* = 0.716, *n* = 17). Bottom: GTEx single-tissue eQTL in heart atrial appendage, where the ALT allele is associated with reduced *ATP13A3* expression (NES = −0.216, *p* = 3.78 × 10^−27^). The opposing directions illustrate a cell type-specific effect in valve fibroblasts, a tissue absent from GTEx. **i,** Schematic of CardioSleuth, an AI-powered exploration engine for cardiac regulatory effects integrating Multiome data, cell state-specific gene regulatory networks, GWAS score propagation (SNP2CELL), and external knowledge bases. An agent-navigated interface supports literature-guided interpretation and graph mining, enabling interactive generation of cell state-resolved hypotheses for cardiovascular disease mechanisms. Abbreviations: ASA, allele-specific accessibility; CAVS, calcific aortic valve stenosis; PIP, posterior inclusion probability; P2G, peak-to-gene; GTEx, Genotype-Tissue Expression; eQTL, expression quantitative trait locus; NES, normalised effect size.

We then applied a beta-binomial likelihood-ratio test for ASA at three hierarchical levels: donor (bulk across all cell types), donor-cell type, and donor-cell state (**Supplementary Fig. S5c**). SNP-group pairs (a SNP tested within a given donor, cell type, or cell state) were tested only if they had ≥10 reads in each of ≥4 heterozygous donors, with a between-donor agreement fraction (the proportion of donors whose allelic direction agreed with the pooled estimate) ≥0.7. At the cell type level, 93 of 161 testable SNP-cell type pairs were significant (at a permissive FDR *<* 0.15) and were prioritised for downstream interpretation, including a subset of SNPs exhibiting ASA in more than one cell type (**Fig. 5b,c**; **Supplementary Fig. S5d**).

Although residual reference-mapping bias cannot be fully excluded, it acts toward the reference allele independently of cell type and therefore cannot account for SNPs we observed with opposing allelic direction between cell types (REF-biased in some, ALT-biased in others; **Supplementary Fig. S5e**). Supporting genuine cell type-specific regulation, 12 SNPs did not show significant ASA at the donor level, but were significant at the donor-cell type level (**Supplementary Fig. S5e,f**).

#### Predicting cell type-specific variant effects with deep-learning sequence models

Recent advances in deep-learning DNA sequence-to-function models have enabled the inference of variant effects, but regulatory effects are frequently cell type-specific ^55^. We therefore fine-tuned two state-of-the-art models on our Multiome dataset: ChromBPNet ^56^ to predict base-resolution chromatin accessibility, and AlphaGenome ^57^ to model both RNA and ATAC profiles. When benchmarking two AlphaGenome fine-tuning strategies, a linear-probe model achieved comparable prediction accuracy to the more costly LoRA-based model (**Supplementary Fig. S5g**); we therefore adopted the linear model for downstream analyses. Fine-tuning with cell type-matched data improved variant effect prediction for both ChromBPNet and AlphaGenome (**Supplementary Fig. S5h-j**). The cell type-matched ChromBPNet model outperformed both a naive Tn5-bias-only baseline and mismatched cell type models (**Supplementary Fig. S5h,i**).

Predicted allelic effects from both models correlated with observed ASA (Pearson *r* = 0.67 and 0.68 for ChromBPNet and AlphaGenome, respectively; **Fig. 5d,e**), with directional concordance of 66% and 76%, respectively. Concordance was likely limited by the modest effect sizes of many common variants, whose allelic direction is harder to call as the predicted effect approaches the null. Together, these results indicate that our cardiac fine-tuned models capture cell type-specific regulatory effects of genetic variation.

#### Mechanistic dissection of GWAS variants

We used the 93 SNP-cell type pairs where we had observed ASA in our human cardiac Multiome to investigate potential disease mechanisms. The cell type specificity of the observed ASA effects provided a basis to narrow the search for effector cell types at each locus. We highlight three illustrative examples.

The atrial fibrillation-associated SNP rs2979489 (chr8:30,423,317 G*>*A) mapped within an ATAC peak accessible in multiple cell types but linked to *RBPMS* expression by ArchR peak-to-gene (P2G) analysis in Atrial Cardiomyocytes only (**Fig. 5f**). ChromBPNet predicted this variant to have the largest effect on chromatin accessibility of any common variant (gnomAD v4.1) in its 50-kb window, consistent with the strong ALT allele bias observed in our data (log_2_FC +0.41). Attribution analysis indicated that the ALT allele creates a MEF2 family motif (*p* = 2.7 × 10^−7^; **Supplementary Fig. S5k**), while AlphaGenome predicted increased expression of *RBPMS* and *RBPMS-AS1* (**Fig. 5f**, **Supplementary Fig. S5l**). *RBPMS* encodes an RNA-binding protein that regulates alternative splicing of sarcomere genes, and its cardiac-specific deletion in mice causes severe contractile dysfunction and dilated cardiomyopathy ^58^. At a locus recently associated with atrial fibrillation ^59^, these findings support *RBPMS* as a plausible effector gene, and our data link the regulatory effect specifically to Atrial Cardiomyocytes.

A calcific aortic valve stenosis-associated variant, rs1706003 (chr3:194,579,238 G*>*T), mapped to a peak with divergent cell type-specific ArchR-derived Peak-to-Gene (P2G) links: accessibility correlated positively with *ATP13A3* expression in Atrial Cardiomyocytes (P2G *r* = 0.21) but negatively in Valve Fibroblasts (P2G *r* = −0.54). In agreement with model predictions, this SNP was observed to associate with reduced accessibility, significant in four cell types including Valve Fibroblasts and Atrial Cardiomyocytes (**Fig. 5b,c**). Attribution analysis indicated destruction of a motif most closely matching CREB1 (*p* = 2.8 × 10^−6^; **Supplementary Fig. S5k**). AlphaGenome predicted reduced expression of several genes in both Atrial Cardiomyocytes and Valve Fibroblasts, but increased expression of *ATP13A3* specifically in Valve Fibroblasts (**Fig. 5g; Supplementary Fig. S5m**). Consistent with this prediction, using our Multiome data, ALT allele dosage showed a significant positive association in Valve Fibroblasts with *ATP13A3* expression (*ρ* = 0.66, *p* = 5.81 × 10^−3^, *n* = 16; **Fig. 5h**), with a non-significant trend toward reduced expression in Atrial Cardiomyocytes (*ρ* = −0.10, *p* = 0.716, *n* = 17). The latter effect is supported by GTEx, where this SNP is a known eQTL for *ATP13A3* in atrial appendage tissue, showing reduced expression with ALT allele dosage (NES = −0.216, *p* = 3.78×10^−27^; **Fig. 5h**). Our data are thus consistent with the GTEx finding but reveal that this aortic stenosis SNP may have a directionally opposite effect on *ATP13A3* expression in Valve Fibroblasts, a cell type absent from GTEx (which lacks heart valve tissue). The opposing directions of effect are compatible with the context-dependent, bidirectional activity of CRE/CREB-family elements, which can activate or repress transcription depending on isoform and cofactor context ^60^, such that loss of this element may reduce accessibility while relieving repression of *ATP13A3* in Valve Fibroblasts. *ATP13A3* encodes a polyamine transporter recently implicated in pulmonary arterial hypertension ^61^, making it an intriguing candidate effector gene for calcific aortic valve stenosis.

A third variant, rs1886512 (chr13:73,946,049 T*>*A), also associated with atrial fibrillation and a known eQTL for *KLF12* in heart tissue, showed particularly strong ASA in Ventricular Cardiomyocytes (log_2_FC +0.46; **Supplementary Fig. S5n**). The effect on *KLF12* expression in Ventricular Cardiomyocytes was also significant in our single-nucleus data (*ρ* = 0.56, *p* = 2.92 × 10^−2^, *n* = 15; **Supplementary Fig. S5o**). *KLF12* encodes a Krüppel-like transcription factor recently shown to promote angiotensin II-induced cardiac remodelling and fibrosis through transcriptional repression of *Smad7* and consequent activation of TGF*β*–Smad3 signalling ^62^. As cardiac fibrosis is a recognised substrate for atrial fibrillation, the genetic link between this variant, *KLF12* expression in Ventricular Cardiomyocytes, and an AF-associated locus suggests a remodelling-mediated mechanism.

Beyond individual loci, AlphaGenome predictions across the analysed SNPs provided insight into the overall predicted trait contributions of different cell types. For instance, coronary artery disease variants showed stronger predicted gene-expression effects in Myeloid Cells, whereas atrial fibrillation variants showed stronger effects in cardiomyocytes (**Supplementary Fig. S5p**). Consistent with prior functional and genetic evidence, *LIPA* showed a prominent myeloid-specific effect ^63^, while *RPL3L* showed cardiomyocyte-enriched effects ^64^ (**Supplementary Fig. S5q**). These results demonstrate that variant-effect predictions can be integrated with cell type-resolved multiomic data to nominate plausible trait-relevant cellular contexts.

#### CardioSleuth: an AI-powered exploration engine for cardiac regulatory effects

To enable exploration of the cell state-resolved gene regulatory networks, variant-effect predictions, and GWAS enrichments identified across the 28 regions and associated cell states, we developed CardioSleuth, an agent-interactive platform for exploring cardiac regulatory effects (**Fig. 5i**). CardioSleuth integrates cell state-specific regulatory networks, GWAS score propagation, precomputed AlphaGenome variant-effect predictions, and external knowledge-base annotations. A literature-grounded agent draws on these resources to support context-aware interpretation of genes, variants, and transcription factors, and enables users to interactively evaluate and curate citations for their specific research questions. CardioSleuth provides an open, agent-assisted framework for translating this heart cellular map into interpretable, cell state-specific hypotheses of cardiovascular disease mechanisms.

## Discussion

We present a multi-region, multi-modal atlas of the adult human heart that extends earlier references ^1–3^ by profiling the anatomically specialised territories in which common cardiovascular disease arises. By sampling the cardiac valves, arteries, pulmonary veins, and multiple myocardial sites, each profiled with orthogonal single-cell and spatial assays, we defined region-enriched cell states, placed them within their native tissue niches, and connected them to non-coding genetic risk. Three findings illustrate this approach: a valve fibroblast–macrophage niche linked to aortic stenosis, a pulmonary vein cardiomyocyte population carrying the principal atrial fibrillation risk signal, and a RUNX1-marked smooth muscle state that constitutes the healthy coronary subintima. In each case, anchoring a disease-associated cell state in its anatomical and regulatory context nominates specific pathways and candidate effector genes for mechanistic dissection and, ultimately, therapeutic targeting. We discuss each in turn, then show how cell type-resolved genetic analysis extends variant interpretation across the atlas.

### A valve-intrinsic immune niche linked to aortic stenosis susceptibility

Calcific aortic valve stenosis is an active fibro-inflammatory and osteogenic disease with a substantial heritable component ^65^, yet it remains treatable only by valve replacement, with no disease-modifying medical therapy. Most cellular insight has come from diseased or stenotic valves, leaving the organisation of the healthy valve poorly defined. By profiling non-diseased tissue, we show that immune activity is not solely a late feature of stenotic remodelling: we resolve Valve Fibroblasts as a distinct stromal compartment and identify Valve Fibroblasts Immune, a subpopulation co-localising with resident macrophages in a connective tissue–immune niche that is spatially polarised to the non-fibrosa aspect of both aortic and mitral valves. Within this niche, Valve Fibroblasts express *ANXA1* and complement components including *C3*, with predicted Annexin and Complement signalling to macrophages, consistent with an established role for innate immune signalling in valve calcification ^66^. In healthy valves these signals need not be pathogenic; alongside the age-associated expansion of Valve Fibroblasts Immune and its increased TNF*α*, TGF*β* and myofibroblast-like expression, they instead suggest a homeostatic stromal–immune niche poised to shift towards fibro-inflammatory remodelling with age.

Integrating cell-state regulatory networks with aortic stenosis GWAS loci prioritised Valve Fibroblasts Immune as a disease-relevant regulatory context, with genetic signal converging on NF-*κ*B, IRF and AP-1 programmes within the fibroblasts themselves rather than immune cells alone. This represents genetic prioritisation rather than proof of causality, but it is reinforced by the cell type-specific rs1706003–*ATP13A3* effect we observe in Valve Fibroblasts. Together these findings recast the healthy valve as harbouring a spatially polarised, age-responsive fibroblast–macrophage niche that is genetically linked to aortic stenosis. Importantly, the pro-resolving ANXA1–formyl peptide receptor axis that organises this niche is pharmacologically tractable in the heart, with selective FPR agonists already advancing in cardiovascular development ^67;68^. Should this niche prove to expand or change regulatory state in diseased valves, the ANXA1/FPR and complement axes would represent candidate entry points for anti-inflammatory intervention upstream of overt calcification, addressing an unmet need in a disease for which no disease-modifying therapy currently exists.

### Myocardial Sleeve Cells define a pulmonary vein cardiomyocyte substrate for atrial fibrillation genetics

Pulmonary veins have long been recognised as the principal source of ectopic electrical activity triggering atrial fibrillation, and pulmonary vein isolation is the cornerstone of catheter ablation ^30;69^. Yet the cellular identity of the human pulmonary vein myocardium has remained poorly resolved, leaving models of sleeve arrhythmogenesis reliant on anatomy, electrophysiology and genetics rather than direct molecular profiling. We identify Myocardial Sleeve Cells, a specialised cardiomyocyte population restricted to the pulmonary veins, marked by *PITX2*, *DHRS9* and *KCNMB2* and showing the strongest PITX2 motif enrichment of any cardiac cell state. This is notable because the *PITX2* locus at 4q25 carries the most reproducible common-variant signal for atrial fibrillation ^5;31^, and *PITX2* is itself central to pulmonary myocardial development and left atrial electrophysiological identity, with reduced dosage promoting arrhythmia susceptibility through left-sided pacemaker-like specification and atrial transcriptional remodelling ^70;71^.

Linking this population to atrial fibrillation genetic risk, the 4q25 locus credible-set SNP rs2220427 overlapped chromatin selectively accessible in Myocardial Sleeve Cells, with peak-to-gene links to *PITX2*, *PANCR* and *LINC01438*. While this does not establish causality, it places a non-coding AF risk signal in a defined cellular context: pulmonary vein sleeve cardiomyocytes. The sleeve cardiomyocyte identity also bore a developmental signature: AF variants were further enriched in a nuclear-receptor regulatory module including retinoic acid receptors, alongside expression of the retinoid-metabolism gene *DHRS9*. Spatially, Myocardial Sleeve Cells did not form a discrete layer but were intermingled with left atrial cardiomyocyte states, suggesting that ectopy and conduction heterogeneity emerge from interactions among sleeve cells, neighbouring myocardium, innervation and matrix rather than from any single cell type in isolation. Consistent with this, we inferred vitronectin signalling between Myocardial Sleeve Cells and neighbouring stressed atrial cardiomyocytes, one example of a local niche interaction that may shape sleeve tissue organisation, though it awaits functional validation.

Clinically, pulmonary vein isolation already targets this region empirically, ablating it without a molecular definition of what is being treated. By resolving the sleeve cardiomyocyte programme and the regulatory circuits through which 4q25 risk may act, our data provide a molecular handle on this substrate that could help explain variation in ablation success and motivate more selective, cell-state-directed antiarrhythmic strategies. That such a transition is feasible is illustrated by TASK-1 (*KCNK3*), an atrial-selective potassium channel whose targeted inhibition can cardiovert atrial fibrillation ^36^. In our data *KCNK3* is selectively expressed by the two cardiomyocyte populations of the sleeve, Myocardial Sleeve Cells and Atrial Cardiomyocytes Left Stressed, placing a tractable antiarrhythmic target within this very substrate.

### A RUNX1-marked smooth muscle state defines the healthy human coronary subintima

Previous single-cell studies have largely characterised modulated smooth muscle cells in established atherosclerosis, including TCF21-associated fibromyocytes ^72;73^ and FAP^+^ intimal smooth muscle cells ^74^. We identify Smooth Muscle Cells Coronary Artery, a synthetic (fibromyocyte) state present in coronary arteries from donors without known cardiovascular disease and localised by spatial profiling to the subintima, where it forms a reproducible inner layer adjacent to the contractile media. This provides molecular resolution for diffuse intimal thickening, a long-recognised feature of human coronary arteries and a proposed substrate for early atherogenesis ^75;76^: rather than an undifferentiated accumulation of medial cells, it comprises a transcriptionally specialised, RUNX1-marked smooth muscle state. Its transmural distribution is preserved in publicly available atherosclerotic coronary data, indicating a normal component of arterial architecture on which disease programmes may later be imposed rather than a product of advanced disease.

Smooth Muscle Cells Coronary Artery expressed *RUNX1* alongside an inflammatory cytokine programme, and showed the strongest RUNX1 motif enrichment of any non-haematopoietic cardiac cell state, extending work implicating RUNX1 in vascular remodelling ^45^ to a steady-state human subintimal population rather than to injury or overt plaque alone. Bidirectional perturbation in primary human smooth muscle cells supported a regulatory role: RUNX1 promoted cell-cycle entry while its knockdown reduced proliferation, and RUNX1 overexpression suppressed the cell-state-defining inflammatory module of which *RUNX1* is itself a member. This points to a negative-feedback architecture in which RUNX1 couples expansion of the subintimal compartment to restraint of its own inflammatory output.

Together these data recast the healthy coronary subintima as a molecularly defined, RUNX1-controlled smooth muscle niche that accumulates with age ^75^. Whether this accumulation is primarily adaptive, maintaining endothelial–medial integrity and limiting inflammation, or whether it renders the intima permissive to lipid retention and plaque initiation when homeostatic control fails, is the central open question, and it carries a direct therapeutic corollary: if RUNX1 activity proves protective it should be reinforced in early coronary disease, whereas if it proves permissive it represents a target for restraint, for which small-molecule RUNX1 inhibitors already exist ^77^.

### Cell type-resolved genetic effects and variant-effect prediction

Most cardiovascular GWAS variants are non-coding and act in cellular contexts that bulk-tissue assays obscure ^55^. Our allele-specific accessibility framework addresses this by comparing alleles within heterozygous donors, resolving regulatory effects even at modest donor numbers and recovering a subset detectable only at cell-type resolution. Coupling these measurements with sequence-to-function models fine-tuned on our data (ChromBPNet and AlphaGenome) let us implicate transcription factor motifs and nominate target genes in the same context ^73;78^. Anchoring non-coding variants in defined cell types in this way converts statistical associations into specific, testable mechanistic hypotheses, as three loci illustrate.

The *ATP13A3* locus illustrates the value of this resolution. The CAVS-associated variant rs1706003 showed reduced accessibility in Valve Fibroblasts with attribution to a disrupted CREB1-like motif, and AlphaGenome predicted increased *ATP13A3* expression specifically in Valve Fibroblasts, corroborated by allele-dosage analysis in our single-nucleus RNA-seq data. By contrast, GTEx atrial appendage data show the same allele reducing *ATP13A3* expression in bulk tissue. This divergence is informative: the variant appears to act in opposite directions across cardiac cell types, and the Valve Fibroblast effect is invisible to GTEx, which lacks valve tissue. *ATP13A3* encodes a P5B-type ATPase of the polyamine transport system ^79^ whose loss-of-function causes pulmonary arterial hypertension through disrupted polyamine handling ^61^. Since valve fibroblasts are the central effectors of fibro-calcific remodelling, our data nominate *ATP13A3*, and polyamine homeostasis more broadly, as a candidate mechanism in aortic stenosis, a pathway that is pharmacologically accessible through existing polyamine-targeting agents ^61^ and that would otherwise have been overlooked by bulk-tissue analysis.

Cell type resolution similarly clarified two atrial fibrillation loci. At the *RBPMS* locus, recently implicated in atrial fibrillation ^59^, we localised the regulatory effect specifically to Atrial Cardiomyocytes; *RBPMS* encodes a cardiac RNA-binding protein required for contractile function through sarcomeric splicing ^58^, providing a plausible route from noncoding risk to atrial structural remodelling. At the *KLF12* locus the regulatory effect instead localised to Ventricular Cardiomyocytes, and given the role of *KLF12* in TGF-*β*–SMAD fibrotic signalling, suggests that some AF risk may act through myocardium-wide structural or stress-response pathways rather than a primarily atrial route.

Together, our Hypersampling strategy establishes a unified molecular view of the adult human heart across 28 anatomical regions. This approach reveals biological differences among related cell types from distinct cardiac regions that are not readily apparent in isolated sampling schemes, enabling refined cell-state annotation and the identification of region-specific cellular niches. By integrating these maps with human genetics, we decompose the cellular contexts through which disease-associated variants may act, nominating candidate effector genes, including *ATP13A3*, *RBPMS* and *KLF12*, and the cell states in which they operate. Beyond cataloguing cardiac cell diversity, anchoring pathways and candidate targets in defined anatomical and regulatory contexts provides a basis for mechanistic study, target prioritisation, and cell-state-informed interpretation of inherited cardiovascular risk to guide future therapy and management.

### Resource availability

This atlas provides several resources for the cardiovascular community: single-nucleus and spatial transcriptomic data across 28 regions, freely available as a Human Cell Atlas ^80^ contribution; fine-tuned ChromBPNet and AlphaGenome models, released to remove a substantial computational barrier for variant-effect prediction; and CardioSleuth, an AI-powered exploration engine for cardiac regulatory effects that integrates gene regulatory networks, GWAS propagation scores, and variant-effect predictions with a literature-grounded conversational agent for hypothesis generation. By linking cellular atlases, regulatory models, human genetics and interactive hypothesis generation, these resources transform the atlas from a static reference into an active discovery platform for cardiovascular biology. All resources are available in interactive form at https://www.heartcellatlas.org/v3/.

### Limitations and future directions

Our donor cohort (36 individuals in total, 4 of them profiled by spatial assays alone) provides broad coverage but does not capture the full spectrum of sex-, ancestry- and age-dependent variation. Spatial technologies involve trade-offs between gene coverage and resolution, and our genetic findings remain largely associative: beyond the functional validation of RUNX1, the cell-type-specific variant effects we describe await experimental confirmation. Integration with disease cohort data, particularly from heart failure, cardiomyopathy, and stenotic valve tissue, will be essential to determine how the cell states and regulatory circuits we describe are perturbed in disease. Computational drug-target predictions (drug2cell) for populations such as Myocardial Sleeve Cells may provide starting points for repurposing screens, although understanding how these cells change in disease remains fundamental. The framework we establish, combining spatially resolved multiomics, cell state-aware variant-effect models, and agent-assisted interpretation, offers a template for similar efforts across organ systems.

## Methods

### Research ethics for donor tissues

All heart tissue samples were obtained from transplant donors following Research Ethics Committee approval and written informed consent from donor families, as previously described ^1^. Additional heart tissue donors were covered by the following ethics approvals: A61, A70 and A72 (REC reference 15/EE/0152, East of England Cambridge South Research Ethics Committee); AH7, AH8, AH9, AH10, AH13 and AH14 (REC reference 16/LO/1568, London, London Bridge Research Ethics Committee); and AV11 and AV12 (REC reference 16/NE/0230, North East, Newcastle & North Tyneside Research Ethics Committee).

### Procurement of hearts

Cardiovascular history was unremarkable for all donors **(Supplementary Table)**. Hearts were procured following donation after circulatory death (D8, A61, A70, A72, AH1, AH2, AH5, AH8, AH9, AH10, AH13, AH14, AV3, AV11 and AV12) or donation after brain death (AH6, AH7, AV1, AV10, AV13 and AV14). For donation after circulatory death donors, sternotomy was performed after confirmation of death and a mandatory 5-min stand-off period. In all cases, the aorta was cross-clamped and cold cardioplegia solution was administered into the aortic root before cardiotomy.

Samples AV1, AV3, AV10, AV11, AV12, AV13 and AV14 were procured using standard procedures and then immediately preserved and transported on a hypothermic perfusion machine. Sample AH2 was similarly preserved, but using immediate normothermic perfusion. It underwent 4 h of normothermic perfusion before tissue samples were collected.

### Dissection

Hearts were stored, transported, and dissected on ice until freezing or fixation to minimise transcriptional degradation. Two operators (JC and LM) performed the dissections according to a predefined dissection plan **(Supplementary Table)**, while other operators (KK, SB, AMAM and SP) preserved samples in frozen OCT medium or 4% formalin. Samples were either frozen with or without OCT embedding and stored at −80*,^◦^* C, or formalin-fixed and subsequently embedded in paraffin blocks.

### Anatomical sampling and region grouping

To capture regional cellular and molecular diversity, we substantially expanded the anatomical sampling relative to our previous eight-region heart atlas ^3^, collecting 28 distinct sampling sites spanning the atria, ventricles, valves and vessels. To support analyses at multiple anatomical scales, each sampling site was placed within a hierarchical taxonomy: individual sites were grouped into 12 coarse anatomical regions (region_coarse), which were in turn nested within four broad tissue compartments (tissue): atrial, ventricular, valve and vessel. The full list of sampling sites, their region and tissue assignments, abbreviations and dissection notes is provided in the **Supplementary Table**.

### Single nuclei isolation

Single nuclei were isolated from flash-frozen tissue by cryosectioning and mechanical homogenisation, as previously described ^3^. Frozen tissue slices of 5–10 mm thickness were sectioned at 50 *µ*m on a cryostat. Sections from each sample were homogenised in a 7,ml glass Dounce tissue grinder set (Merck) with 8–10 strokes of loose pestle A followed by 8–10 strokes of tight pestle B in homogenisation buffer: 250,mM sucrose, 25,mM KCl, 5,mM MgCl_2_, 10,mM Tris-HCl, 1,mM dithiothreitol (DTT), 1× protease inhibitor, 0.4,U,*µ*l^−1^ RNaseIn, 0.2,U,*µ*l^−1^ SUPERaseIn and 0.1% Triton X-100 in nuclease-free water.

Homogenates were filtered through a 40,*µ*m cell strainer (Corning) and centrifuged at 500,*g* for 5,min at 4,*^◦^*C. Supernatant was removed and nuclei were resuspended in storage buffer containing 1× PBS, 4% BSA and 0.2,U,*µ*l^−1^ Protector RNaseIn. Nuclei were stained with 7-AAD viability staining solution (BioLegend), and 7-AAD-positive single nuclei were purified by fluorescence-activated cell sorting using an MA900 Multi-Application Cell Sorter (Sony) with Cell Sorter software v.3.1.1. Nuclei purity and integrity were confirmed by microscopy.

### 10x Multiome library preparation

For 10x Multiome library preparation, nuclei were manually counted by Trypan blue exclusion, diluted to 1,000–3,000 nuclei per microlitre and loaded onto a Chromium Controller (10x Genomics), targeting recovery of 5,000–10,000 nuclei per reaction. Paired 3*^′^* gene expression and ATAC libraries were prepared using the Chromium Single Cell Multiome ATAC + Gene Expression kit (10x Genomics), according to the manufacturer’s instructions. cDNA and final library quality control were performed using a Bioanalyzer High Sensitivity DNA assay or 4200 TapeStation system (Agilent). Libraries were sequenced on a NovaSeq 6000 platform (Illumina) at the Wellcome Sanger Institute to a minimum depth of 20,000–30,000 read pairs per nucleus.

### Visium and Visium HD library preparation

Fresh-frozen samples were embedded in OCT medium and frozen in a dry ice-cooled isopentane bath at −45 *^◦^*C. OCT-embedded samples were sectioned at 10 *µ*m using a cryostat (Leica CM3050S).

For formalin-fixed paraffin-embedded (FFPE) samples, fresh tissue was fixed in *>*5 times its volume of 4% v/v formalin at ambient temperature for 24 h before processing to paraffin using a Tissue-Tek Vacuum Infiltration Processor 5 (Sakura Finetek). FFPE blocks were sectioned at 5 *µ*m using a microtome (Leica RM2125RT).

Microanatomical regions of interest (ROIs) were selected on the basis of morphology, expert review (SYH), tissue orientation assessed by H&E staining, and RNA quality. RNA integrity number was used for fresh-frozen samples and DV200 for FFPE samples, measured using an Agilent 2100 Bioanalyzer. FFPE tissues were additionally assessed for potential detachment using 10x Genomics Adhesion Test slides.

Fresh-frozen Visium Spatial Gene Expression experiments were performed according to the manufacturer’s protocol (10x Genomics). Tissue optimisation was first performed to determine the optimal permeabilisation time, which was 45 min. FFPE Visium Spatial Gene Expression experiments were also performed according to the manufacturer’s protocol (10x Genomics).

For selected FFPE sections, Visium CytAssist Spatial Gene Expression v2 was performed according to the manufacturer’s protocol (10x Genomics). FFPE tissue sections on standard glass slides were deparaffinised, H&E-stained, imaged and decrosslinked before probe-based library preparation. After probe hybridisation, tissue slides and Visium CytAssist slides were loaded onto the CytAssist instrument, which aligned tissue sections with the Visium capture areas and transferred analyte-derived molecules to spatially barcoded oligonucleotides on the Visium slide. Probe extension and downstream library construction were then performed according to the manufacturer’s protocol.

Two FFPE samples were processed using the Visium HD Spatial Gene Expression protocol according to the manufacturer’s instructions (10x Genomics). FFPE tissue sections on standard glass slides were deparaffinised, H&E-stained, imaged, destained and decrosslinked before whole-transcriptome probe hybridisation and ligation. Tissue slides and Visium HD slides were loaded onto the CytAssist instrument for CytAssist-enabled probe release and capture by spatially barcoded oligonucleotides on the Visium HD slide. Captured probes were extended to incorporate spatial barcodes and unique molecular identifiers, followed by probe elution, pre-amplification, sample indexing, SPRIselect cleanup and library quality control according to the manufacturer’s protocol.

H&E-stained Visium Gene Expression slides were imaged at 40× magnification using a Hamamatsu NanoZoomer S60. After transcript capture, Visium libraries were prepared using the 10x Genomics Visium Library Preparation protocol. Eight cDNA libraries were diluted and pooled to a final concentration of 2.25 nM in a 200 *µ*l volume, and sequenced on two SP flow cells using an Illumina NovaSeq 6000.

### Xenium in situ gene expression

#### Xenium and Xenium 5K assays

Xenium Spatial Transcriptomic assays were performed using Human Multi-Tissue and Cancer panel of 377 genes on 8 slides across 4 runs. The anatomical regions included LV apex (2), LV lateral wall (1), LV anterolateral wall (1), left atrial appendage (1), right atrial appendage (1), sinoatrial node (1), pulmonary vein ostia (3), proximal left anterior descending artery (3), proximal ascending aorta (1), aortic valve (2) and anterior mitral valve leaflet (2).

#### Post-Xenium immunofluorescence and H&E imaging

Multiplex immunofluorescence staining and single-round 16-plex imaging were performed on Formalin-fixed paraffin-embedded (FFPE) human heart tissue blocks which were sectioned at 5 µm thickness using a microtome (Leica RM2235) and mounted onto SuperFrost Plus slides. Slides were dried at 60 °C for 60 min to ensure tissue adherence. Tissue sections were deparaffinized and subjected to antigen retrieval using the BioGenex EZ-Retriever system (95 °C for 5 min followed by 107 °C for 5 min). To reduce tissue autofluorescence, slides were treated with AF quench buffer (4.5% HO, 24 mM NaOH in PBS). Slides were quenched for 60 min using the high setting with strong white-light exposure followed by a further 30 min quenching step using the 365 nm high setting with a UV transilluminator. Slides were rinsed with 1× PBS and incubated with 300 µl Image-iT FX Signal Enhancer (Thermo Fisher Scientific, I36933) for 15 min. Following PBS washes, 300 µl of a 16-plex fluorophore-conjugated primary antibody cocktail was applied to each tissue section and incubated for 120 min in the dark within a humidified chamber. Antibodies were used according to manufacturer recommendations. Slides were subsequently washed with Orion surfactant wash buffer and incubated with 300 µl nuclear stain diluted in goat diluent for 30 min in the dark. Following additional washes, slides were maintained in 1× PBS and coverslipped using ArgoFluor mounting medium. Slides were allowed to dry overnight in the dark at room temperature. Whole-slide multiplex images were acquired the following day using the RareCyte Orion imaging platform equipped with a 20× objective lens. Spectral unmixing, autofluorescence correction, image registration and image processing were performed using Artemis software. Processed images were subsequently used for downstream cell segmentation, phenotyping and spatial analyses.

#### Xenium co-registration

For each Xenium section, the post-Xenium haematoxylin and eosin (H&E) and multiplexed immunofluorescence images were registered to the Xenium coordinate system, using the Xenium DAPI morphology image as the reference. Each section was linked to its corresponding H&E and immunofluorescence files through a manually curated imaging metadata table. Corresponding landmarks were placed by hand on each image pair in BigWarp ^81^ within Fiji ^82^, and from these landmark correspondences an affine transformation was estimated by least squares for the H&E-to-immunofluorescence and the immunofluorescence-to-Xenium mappings with scikit-image ^83^. Landmark coordinates were expressed in the pixel space of a defined pyramid level of each image, whose physical pixel size was read from the image metadata, and registration accuracy was summarised as the mean and maximum landmark residual. The H&E-to-Xenium transformation was obtained by composing the H&E-to-immunofluorescence and immunofluorescence-to-Xenium affines, and the resulting alignments were inspected visually as checkerboard overlays of each warped image pair.

To transfer the imaging data onto the Xenium cells, the segmented Xenium cell centroids (in microns) were converted to the pixel grid of each modality at a common pyramid level by inverting the corresponding affine transformations. For every cell, the mean immunofluorescence intensity of each channel and the mean Xenium morphology (DAPI) intensity were sampled within a 5 *µ*m-radius disc, and a 10 *µ*m H&E patch was summarised by its per-channel colour statistics and grey-level co-occurrence texture features; the three modalities were assembled into a single MuData object carrying both native and Xenium-aligned coordinates for each cell. The alignment was validated by confirming that, across matched antibody–transcript pairs, immunofluorescence protein intensity was higher in Xenium cells expressing the corresponding gene (quantified as the area under the receiver-operating-characteristic curve), and that the independently measured Xenium DAPI and immunofluorescence Hoechst nuclear signals were correlated per cell.

### Immunofluorescence staining

Fresh frozen anterior mitral valve leaflet (n=3) and aortic valve leaflet (n=3) tissue sections from independent donor hearts were used. To delimit the area of staining hydrophobic pen was used on all tissue slides. Optimisation of final concentration of primary antibodies was performed. Slides were directly transferred from −80°C to 4°C 4% paraformaldehyde (PFA) solution and incubated for 5 mins. Slides were blocked in 4% bovine serum albumin (BSA) (Sigma Aldrich, A3059), 1% donkey serum and 0.2% Triton X-100 (Thermo Fisher, 85111) for 30 mins at room temperature. Anti-MYH11 (Santa Cruz, sc-79079) and anti-COMP (Abcam, ab11056) were diluted 1:100 in 4% BSA, 1% donkey serum, 0.2% Triton X-100 and incubated overnight at 4°C. Nuclei were stained with DAPI (Thermo Scientific, 62248) with 1:1000 dilution in mounting medium and incubated overnight at 4°C. Secondary antibodies Alexa Fluor 488 donkey anti-goat (Invitrogen, A11055) and Alexa Fluor 594 donkey anti-rat (Invitrogen, A21209) were diluted 1:200 in filtered Phosphate Buffered Saline (PBS) and incubated for 2 hours at 4°C. Isotype control and secondary antibody only staining were performed as negative controls. Rat IgG2a isotype (BD Biosciences, 553930) and goat IgG isotype (R&D Systems, AB-108-C) with matched concentrations to the corresponding primary antibodies were used. For secondary antibody staining, samples were incubated with 4% BSA, 1% donkey serum, 0.2% Triton X-100 followed by the secondary antibody incubation.

### Read mapping and donor demultiplexing

After sequencing, samples were demultiplexed and stored as CRAM files. scRNA-seq and snRNA-seq samples were mapped to the human reference genome GRCh38-3.0.0, provided by 10x Genomics, using Cell Ranger v.3.0.2 with default parameters. For single-nucleus samples, a pre-mRNA reference was generated according to the 10x Genomics instructions. Multiome and Visium samples were mapped to 10x Genomics human reference genomes using Cell Ranger ARC v.2.0.0 or Space Ranger v.1.1.0, respectively, with default parameters. Multiome samples were mapped to GRCh38-2020-A-2.0.0 and Visium samples to GRCh38-3.0.0. Visium HD samples were processed using Space Ranger v.3.0.0. For Visium datasets, Space Ranger was also used to align paired histology images with mRNA capture spot positions on the Visium slides. For cost-efficient experimental design, cells or nuclei from multiple donors of the same anatomical region were pooled before library preparation. Donor identities were subsequently assigned computationally using souporcell ^84^, which clusters single-cell or single-nucleus profiles by genotype at common variant sites without requiring prior per-donor reference genotypes. Genotyping was performed against a set of common variants from the 1000 Genomes Project ^85^ (GRCh38), restricted to the cell barcodes retained after CellBender filtering, with the number of genotype clusters set to the number of pooled donors. For Multiome libraries, the gene-expression and ATAC read alignments were merged prior to genotyping, so that genotype-informative variants from both modalities contributed to donor assignment. The anonymous genotype clusters obtained for each pool were then resolved to donor identities by comparing cluster genotypes, namely the variant allele frequencies at shared variant sites, against single-donor reference libraries where available and against genotype clusters from other pools where no single-donor reference was available, using the GenotypeMixtures package (https://github.com/bjstewart1/GenotypeMixtures); the resulting cluster-to-donor assignments were curated manually. Singlets were retained for downstream analysis, whereas cross-donor doublets and cells or nuclei that could not be confidently assigned to a donor were excluded.

### Quality control and preprocessing

For scRNA-seq, snRNA-seq and Multiome gene expression data, ambient and background RNA were removed from each raw count matrix using CellBender v0.2.2 remove-background ^86^. CellBender was run per sample on raw Cell Ranger or Cell Ranger ARC count matrices. The expected number of cells or nuclei was estimated from the corresponding Cell Ranger metrics, and the number of droplets included for background modelling was set to this value plus 10,000 after manual inspection of UMI knee plots. Quality control and downstream preprocessing were performed in Python using Pandas, NumPy, Matplotlib, Scanpy ^87^ and Scrublet ^88^. Very low-quality barcodes were first removed by requiring at least 100 detected genes and 100 counts. Putative doublets were identified using Scrublet v.0.2.3, which was run separately for each sample ^88^. Scrublet scores were incorporated into the downstream quality control workflow, with an initial maximum threshold of 0.3. Quality control metrics were then calculated for each cell or nucleus, including total counts, number of unique genes, percentage of mitochondrial reads, percentage of ribosomal reads, percentage of haemoglobin reads and Scrublet doublet score. Cells and nuclei were filtered using the automated quality control workflow autoQC ^89^. First, cell-level QC calls were generated using Gaussian mixture models fitted to QC metric distributions. The initial metric bounds were 200–6,000 detected genes, 500–20,000 counts, less than 25% mitochondrial reads for cells or 5% for nuclei, less than 30% ribosomal reads for cells or 5% for nuclei, less than 2% haemoglobin reads and a Scrublet score below 0.3. QC-space clusters were then generated from total counts, detected genes, mitochondrial percentage, ribosomal percentage, haemoglobin percentage and Scrublet score. Cluster-level QC calls were assigned using autoQC, with clusters retained if at least 50% of constituent cells or nuclei passed cell-level QC. Cells or nuclei failing cell-level QC, assigned to failing QC clusters, or identified as putative doublets were excluded from downstream analysis. Downstream preprocessing was performed using Scanpy.

For Multiome ATAC data, fragment files generated by Cell Ranger ARC were further analysed using ArchR v.1.0.2^90^. ArchR was run using the hg38 genome. Arrow files were generated for each sample using createArrowFiles, requiring a minimum transcription start site enrichment score of 4 and at least 1,000 fragments per cell, with TileMatrix and GeneScoreMatrix generation enabled. Sample-level ATAC quality control was performed by inspecting transcription start site enrichment and log_10_ fragment count distributions. Additional ATAC quality metrics considered during preprocessing included nucleosomal banding patterns, the number and fraction of fragments in peaks, reads overlapping ENCODE blacklist regions and ArchR doublet scores, where available.

For high-quality cells, chromatin accessibility was summarised in 500-bp genome-wide bins using the ArchR TileMa-trix. Gene scores were computed from chromatin accessibility around genes and compared with paired gene expression profiles from the same Multiome nuclei. Cell type and cell state annotations derived from the paired gene expression data were added to the ArchR project and used for downstream ATAC analysis.

Before peak calling, pseudo-bulk replicates were generated for each fine-grained cell state using addGroupCoverages. Three pseudo-bulk replicates were generated per cell state, with 300–1,000 cells per replicate and a maximum of 25×10^7^ fragments. Reproducible peaks were called for each cell state using addReproduciblePeakSet, requiring reproducibility in at least two pseudo-bulk replicates. Cell-state peak sets were then merged to obtain a unified peak set, and a cell-by-peak count matrix was generated using addPeakMatrix.

For Visium data, the Scanpy toolkit was used for quality control and downstream processing. Visium spots of each sample were filtered for more than 500 UMI counts and 300 genes.

### Single-cell data integration and cell-type annotation

#### Single-cell gene expression integration

Batch-corrected latent representations of the gene-expression data were learned with inVAE^6^, which served as the primary integration method. A factorised inVAE model with a negative-binomial decoder was trained on the raw counts of the highly variable genes, separately for each major cell type and for the full (global) dataset. Anatomical region (region_-coarse) was supplied as the invariant (biological) categorical covariate, and a composite batch_key, combining donor identity, assay input type (whole cell versus single nucleus) and 10x Genomics chemistry, as the spurious (technical) categorical covariate. The covariates were injected into the latent space, so that a single 20-dimensional invariant latent representation was learned, and retained as the integrated, batch-corrected embedding. Models were trained for 500 epochs (200 for the global dataset) with a learning rate of 10^−3^ and weight decay of 10^−4^.

A secondary latent representation was generated with scVI^91^, trained per cell type on the raw counts of the highly variable genes using the same composite batch_key as the batch covariate; total counts, the percentage of mitochondrial reads and the percentage of ribosomal reads were supplied as continuous covariates, and gene- and batch-specific dispersion was used. This model used a 30-dimensional latent space and a three-layer encoder (increased to 4 layers for the global dataset). This representation was used only to corroborate the robustness of the annotations.

For each cell-type object, a *k*-nearest-neighbour graph and a UMAP embedding (minimum distance 0.1) were computed on the inVAE invariant latent in Scanpy ^87^, and Leiden clustering was performed on this graph across a range of resolutions. These inVAE-derived neighbour graphs and clusterings provided the basis for the cell-type and cell-state annotation described below.

#### Cell type and cell state annotation

Annotation proceeded in two stages: broad cell-type classification across the full dataset, followed by cell-state annotation within each cell type. The integrated global object was clustered with Leiden on the inVAE neighbour graph, and the resulting clusters were assigned to 18 major cell types by canonical marker expression. Each major cell type was then taken forward as a separate object for cell-state annotation. From the Fibroblasts object, a clearly separate cluster of Nerve Fibroblasts was identified and annotated with cell states separately.

Within each cell-type object, genes detected in fewer than three cells and donor batches containing fewer than five cells were removed, counts were normalised to 10,000 per cell and log-transformed, and highly variable genes were identified. Cells were first over-clustered by Leiden clustering on the inVAE neighbour graph to identify small clusters representing potential doublets, low-quality cells or lineage contaminants, which were removed using a rules-based framework based on contaminant-lineage marker panels, canonical markers of the target lineage, expressing-cell-proportion thresholds and Scrublet scores.

The cleaned cell-type subset was then re-clustered on the inVAE neighbour graph across multiple resolutions to define biologically interpretable substructure, with the working resolution selected on the basis of canonical marker expression, anatomical enrichment and separation in the inVAE and scVI latent spaces. Final cell-state labels were assigned by marker-based curation, and any remaining unassigned clusters were merged into the nearest annotated cell state by Euclidean distance between cluster centroids in the latent space. Final annotations were assessed using differential expression analysis, marker dot plots, dendrograms, UMAP visualisation and anatomical composition plots.

### Annotation robustness assessment

To confirm that the cell type and cell state annotations were not specific to the integration framework in which they were defined, their recoverability was tested in an independent scVI latent space, which does not use the invariant covariate employed by inVAE. Using five-fold cross-validated *k*-nearest-neighbour classification (*k* = 15; scikit-learn ^92^), we quantified how accurately the inVAE-derived labels could be recovered: cell type labels for the global object, and cell state labels within each individual cell type subset. For subsets exceeding 50,000 cells, accuracies were computed on a stratified random subsample of 50,000 cells (fixed seed) to ensure tractability. The same cross-validated classification was also performed within the inVAE latent space itself, so that the per-subset recovery accuracies of the two embeddings could be compared. As a complementary, label-independent measure of structural similarity between the embeddings, the mean Jaccard overlap of each cell’s 15 nearest neighbours was computed between the scVI and inVAE latent spaces.

### Variance partitioning of cell state composition

For each cell type and cell state, the sources of variation in its abundance across samples were quantified by variance partitioning. Per-sample compositions were obtained by dividing the number of cells assigned to a given type or state in each sample by that sample’s total cell count, and the resulting proportions were arcsine-square-root transformed to stabilise the variance and to accommodate zero values. For each cell type or state in turn, the transformed proportion was modelled across the 304 samples (31 donors spanning 28 anatomical regions) by ordinary least squares as a function of anatomical region, donor, sex, 10x Genomics kit version and the cell-versus-nucleus protocol, entered as sum-coded categorical covariates, together with donor age as a continuous covariate, with all terms treated as fixed effects. Type II sums of squares were computed from each fitted model with statsmodels, and the fraction of variance attributable to each covariate, and to the residual, was reported as *η*^2^, the term sum of squares divided by the total sum of squares. Samples with missing covariate values were excluded, and the mean proportion of each cell type or state within each anatomical region was additionally z-scored across regions for visualisation.

### Regional specificity of cell state composition

To distinguish cell populations that are spatially restricted from those that are broadly distributed but modulated in abundance across regions, the regional specificity of each cell type and cell state was summarised with a Gini index. For each cell type or state, the per-sample proportions described above were averaged within each anatomical region to give a vector of mean proportions across regions, and the Gini coefficient of this vector was computed. The Gini index ranges from 0, when a population is distributed evenly across all regions (ubiquitous), to 1, when its abundance is concentrated in a single region (maximally restricted). The index was computed on mean proportions rather than on z-scored values, because z-scoring centres each population at zero and removes the differences in absolute abundance that distinguish restricted from ubiquitous populations. Cell types and cell states were ranked by their Gini index to order them from most to least regionally restricted. This specificity measure is complementary to the variance partitioning above: variance partitioning quantifies whether anatomical region explains a reproducible fraction of the variation in a population’s abundance, whereas the Gini index quantifies how unevenly that abundance is distributed across regions, so that a ubiquitous population subject to consistent regional modulation can score highly in the former while remaining low in the latter.

### Cell cycle scoring

Cells were defined as proliferating if they had a non-zero count for at least one of the canonical proliferation markers *MKI67*, *TOP2A* and *CDK1*. For each smooth muscle cell state, the proliferating fraction was calculated per sample as the number of proliferating cells divided by the total number of cells of that state in the sample, and sample–state combinations containing fewer than 200 cells were excluded to stabilise the proportion estimates.

Differences in the proliferating fraction between smooth muscle cell states were assessed with a binomial generalised linear model with a logit link, modelling the per-sample proliferating proportion as a function of cell state and weighting each observation by the number of cells contributing to it; arterial smooth muscle cells were set as the reference state, and model coefficients were exponentiated to odds ratios with 95% confidence intervals. Analyses were performed with statsmodels.

### Gene expression module analysis

Coordinated gene-expression programmes within individual cell types, including the cardiac conduction system and smooth muscle cells, were identified with Hotspot v1.1.3^93^. For each cell type, Hotspot was run on the cell-type-specific expression object using the normal gene model with the per-cell total counts as the size factor, restricting to the top 5,000 highly variable genes. A *k*-nearest-neighbour graph was built directly on the inVAE invariant latent representation of the cells, the autocorrelation of each gene over this graph was tested, and genes with statistically significant autocorrelation (Benjamini–Hochberg FDR) were carried forward. Pairwise local correlations were computed among the retained genes, and co-expression modules were defined from the local-correlation Z-scores, requiring a minimum of 30 core genes per module and a false-discovery-rate threshold of 10^−8^. Per-cell module scores were then computed.

Module scores were summarised as the median score per cell state and displayed as a hierarchically clustered heatmap, projected onto a UMAP of the inVAE latent representation, and the genes of selected modules were shown across cell states as dot plots of per-gene-scaled mean expression in Scanpy ^87^. For each module of interest, the constituent genes were tested for functional enrichment with g:Profiler ^94^, and significant terms from the Gene Ontology (biological process and cellular component), KEGG, Reactome and WikiPathways were reported, with significance assessed from the g:SCS-corrected *P* value.

### Differential abundance and differential expression analysis

Age-associated changes in valve fibroblast and smooth muscle cell composition were quantified using the Milo ^95^ implementation in Pertpy (v1.0.3) ^96^. Cells were embedded using the latent space of inVAE, and a k-nearest-neighbour graph was constructed on this embedding. Representative neighbourhoods were sampled across the graph, and cells were counted per neighbourhood for each donor. Differential abundance across age was tested with a negative-binomial generalised linear model using edgeR ^97^. Each neighbourhood was annotated to the majority cell state. Neighbourhoods with a spatial false-discovery rate below 0.1 were considered significantly age-associated, and log-fold changes were displayed as beeswarm plots grouped by cell state.

For cell states showing significant age-associated differential abundance, donor-pseudobulk differential expression between age groups was performed with edgeR (v4.8.2). Differentially expressed genes were then tested for functional over-representation using Enrichr (gseapy v1.1.11) ^98^ against the MSigDB Hallmark 2020, GO Biological Process, KEGG, and Reactome gene-set libraries, with enrichment significance assessed by the Benjamini–Hochberg-adjusted P value.

### drug2cell analysis

The targets of the drugs in the ChEMBL database were identified in Visium SD Spatial data and snRNAseq data using per-spot / per cell scoring using drug2cell ^3^. Individual genes encoding targets for drugs of interest were explored in Visium and snRNAseq data using scanpy.

### Cell-cell communication analysis (CellPhoneDB)

Cell–cell communication within the coronary artery niche was inferred from the single-nucleus RNA-seq data with Cell-PhoneDB v5.0.0^49^. The atlas was subset to the smooth muscle and endothelial cell states comprising the niche –arterial and coronary-artery smooth muscle cells, and arterial, large-arterial and capillary endothelial cells – and the log-normalised expression of these cells, labelled by cell state, was analysed with the statistical-inference method. All selected states were assigned to a single microenvironment so that interactions were tested among them; genes were matched by Ensembl identifier, a ligand or receptor subunit was required to be expressed in at least 10% of the cells of a state, statistical significance was assessed from 1,000 random permutations of the cell-state labels with a threshold of *P <* 0.05 (random seed 42), and interaction scores were computed.

Statistically significant interactions between endothelial and smooth muscle cell states were then examined for selected signalling axes and displayed as dot plots of interaction mean and permutation significance using ktplotspy. The expression of the corresponding ligand and receptor genes was additionally shown across the niche cell states as a dot plot of mean expression scaled per gene in Scanpy ^87^.

### NOTCH2/NOTCH3 neighbourhood analysis

To compare *NOTCH2* and *NOTCH3* expression across smooth muscle cell states while mitigating single-cell dropout, cells were partitioned into transcriptional neighbourhoods by Leiden clustering on the inVAE latent space (n_neighbors = 20), with the clustering resolution tuned to yield a mean neighbourhood size of approximately 50 cells. Counts were normalised to 10,000 per cell and log-transformed, and mean *NOTCH2* and *NOTCH3* expression was computed per neighbourhood. Each neighbourhood was assigned to a dominant cell state where a single state comprised more than 75% of its cells; neighbourhoods not meeting this threshold were classed as mixed and excluded from the plot.

### Characterisation of *NOVA1*-expressing endothelial states

Endothelial cells were subset from the integrated object and re-normalised (scanpy normalize_total to 10^4^ counts followed by log1p). Because the *NOVA1*-expressing states were enriched among single-nucleus libraries, marker analysis was restricted to single-nucleus profiles to avoid confounding by dissociation-induced immediate-early and heat-shock genes. To control for ambient cardiomyocyte RNA characteristic of atrial tissue, differential expression was performed within atrial nuclei only, comparing each *NOVA1* endothelial state against all non-*NOVA1* atrial endothelial cells (rank_-genes_groups, Wilcoxon rank-sum test, Benjamini–Hochberg correction). Genes shared across all three states (log_2_ fold-change *>* 1, adjusted *p <* 0.05, expressed in *>* 10% of cells) were taken as the common *NOVA1* endothelial programme. Cell-type specificity of *NOVA1* within the right atrium was assessed by comparing mean expression and the fraction of expressing cells across all annotated cell types in RA and SAN samples.

### Motif enrichment analysis (chromVAR)

Transcription-factor motif activity was estimated from the single-nucleus ATAC peak matrix of the Multiome data with chromVAR^99^, as implemented in ArchR ^90^ (hg38). Two motif collections were used: the cisBP collection ^100^ bundled with ArchR, from which the RUNX1 result was taken, and the human CORE collection of JASPAR2024 ^101^, from which the PITX2 result was taken; the JASPAR2024 motif was used for PITX2 because its position weight matrix more faithfully represented the PITX2 binding preference than the cisBP entry. Motif annotations were added to the peak set with addMotifAnnotations (for JASPAR2024, the CORE human position weight matrices were supplied as a custom motif set, with a motif-matching *P*-value cutoff of 5× 10^−5^ and a scanning width of 8 bp), and per-cell bias-corrected deviations and deviation z-scores were then computed across the peak set with addDeviationsMatrix.

To resolve motif activity by cell state, the per-cell deviations were summarised by taking the median deviation within each cell state, restricting to cell states represented by at least 30 cells (sparser states were considered to have unreliable chromatin profiles). For a given transcription factor, cell states were ranked by their median deviation for that motif and displayed as a ranked scatter plot, with each cell state coloured by its parent cell type.

### Peak-to-gene linkage

Peak-to-gene links were computed from the paired single-nucleus ATAC and gene-expression Multiome data in ArchR ^90^ (hg38). For the full dataset, an iterative LSI reduced dimensionality was first computed from the tile matrix (two iterations, 25,000 variable features, 30 dimensions; clustering resolution 0.2), and addPeak2GeneLinks was then run on this embedding together with the gene-expression matrix, correlating the accessibility of each peak with the expression of genes within the default 250-kb window across the dataset to yield a genome-wide table of peak–gene correlations.

To obtain cell-type-resolved links, the same procedure (iterative LSI followed by addPeak2GeneLinks) was repeated within each cell type and, separately, within each cell state, after subsetting the project to the corresponding cells; cell types or states represented by fewer than 50 cells were excluded, as their chromatin profiles were considered too sparse for reliable estimation. This produced, for each population, a separate set of peak–gene correlations that could be compared across cell types and states.

### Visium spatial analysis

Visium spatial samples from 8 regions of human hearts that were previously produced and published by our group were included. ^3^ Majority of the samples were processed using fresh frozen, OCT embedded tissue with version 1 Visium Spatial chemistry. SAN node samples from 1 donor were processed from FFPE tissue using FF Visium Spatial Gene Expression version 1 chemistry. Visium experiments from samples from additional regions were obtained using Visium Spatial FFPE version 2 chemistry according to manufacturer’s protocol. SpaceRanger was used to align paired histology images with the positions of mRNA capture spots on the Visium slides.

#### Structural annotation

Histological structures of all tissue samples were manually annotated by a clinician (LM) and reviewed by an expert anatomist (SYH) and two clinical histopathologists (MM & JLR) prior to data integration and analysis using Loupe Browser (10X Genomics). Each 55 *µ*m spot was assigned one of the following labels: ‘AV_bundle’, ‘Cardiac_skele-ton’, ‘Endocardium’, ‘Epicardium_subepicardium’, ‘Fat’, ‘Fibrosis_interstitial_atrial’, ‘Fibrosis_interstitial_ventricular’, ‘Fi-brosis_subendocardial’, ‘Ganglia’, ‘Haemorrhage’, ‘Membraneous_septum’, ‘Myocardial_sleeve’, ‘Myocardium_atrial’, ‘Myocardium_fat_atrial’, ‘Myocardium_fat_ventricular’, ‘Myocardium_fibrosis_interstitial_atrial’, ‘Myocardium_fibrosis_-interstitial_ventricular’, ‘Myocardium_ventricular’, ‘Nerve’, ‘Node’, ‘Purkinje_cell’, ‘Smooth_muscle_CS’, ‘Smooth_mus-cle_ISV’, ‘Smooth_muscle_IVS’, ‘Smooth_muscle_atrial’, ‘Valve’, ‘Valve_endocardium’, ‘Valve_fat’, ‘Valve_hamorrhage’, ‘Valve_myocardium’, ‘Valve_smooth_muscle’, ‘Vessel’, ‘Vessel_adventitia’, ‘Vessel_intima’, ‘Vessel_media’. Subsequently, the annotations were nested in the following broader categories “new_structural_annotation_broad”: ‘Conduction_sys-tem’, ‘Connective_tissue’, ‘Endocardium’, ‘Epicardium_subepicardium’, ‘Fat’, ‘Ganglia’, ‘Haemorrhage’, ‘Myocardial_-sleeve’, ‘Myocardium_atrial’, ‘Myocardium_ventricular’, ‘Nerve’, ‘Smooth_muscle_non_vascular’, ‘Valve’, ‘Vessel’, ‘Ves-sel_adventitia’, ‘Vessel_intima’, ‘Vessel_media’.

#### cell2location deconvolution

cell2location was used to deconvolve oligocellular 55 *µ*m spots of Visium SD data as described previously. ^3;8^ Reference signatures of all cell states identified in sc/snRNAseq data were obtained using cell2location package and their location predicted in Visium Spatial spots. For downstream analyses, raw cell2location outputs for each cell state in each spot were normalised to the sum of cell2location outputs for all cell states predicted in each spot.

#### Visium data integration and niche identification

Visium spots not overlying tissue or without identifiable histological identity (debris) were removed prior to integration. For the purpose of data integration, the gene expression space was limited to 17,987 genes that were measured across all 3 Visium chemistries used. inVAE, was used to integrate Visium SD data using the following parameters: n_epochs=200, lr_train=0.001, weight_decay=0.0001, latent_dim_spur = 10, latent_dim_inv = 10, spur_covar_keys_cat=[’sample_de-multiplexed2’,’kit_10x’] (representing individual tissue samples and the Visium spatial chemistry version), inv_covar_-keys_cat = [“new_structural_annotation_broad”] (representing the manual annotation of histological features) ^6^. Neighbours were computed using scanpy.pp.neighbors() from inVAE invariant latent representation followed by Leiden clustering.

The abundance of cell states predicted from cell2location, the structural annotation of histological features and the expression of maker genes was used to annotate Leiden clusters with biologically relevant names. To identify nerves and neural ganglia which are small, but histologically distinct features, adipose tissue cluster was separated from the object, reintegrated and sub-clustered.

Two Leiden clusters were described as low quality control clusters arising from tissue processing and technical artifacts. One enriching for haemoglobin genes and overlapping with areas of tissue haemorrhage and the second characterised by low levels of cardiomyocyte genes likely representing ambient RNA from neighbouring regions while the spatial localisation clearly indicated areas of sparsely cellular dense connective tissue.

An *a priori* goal of comparing myocardium and conduction system niches between different regions of the heart required 1. Merging of all cardiomyocyte-enriched Leiden clusters into a shared label ‘Myocardium’. 2. Dividing this niche based on the anatomical region of origin producing the following niche labels ‘AVN_Myocardium’, ‘AX_My-ocardium’, ‘IVS_Myocardium’, ‘LA_Myocardium’, ‘LV_Myocardium’, ‘Myocardial_Sleeve’, ‘RA_Myocardium’, ‘RV_My-ocardium’, ‘SAN_Myocardium’. 3. Dividing the the conduction system niche by region to identify: ‘AVN’, ‘IVS_Con-ductionSystem’ and ‘SAN’.

#### OrganAxis spatial gradient analysis

In cardiac valve samples, the flow exposed surfaces of the valves (the atrialis of the mitral valve and the ventricularis of the aortic valve) and pressure exposed fibrosa surfaces of the valves were annotated based on H&E images by 2 trained observers (LM & SYH) using TissueTag and the relative distances between the surfaces were computed using OrganAxis. ^102^ In vascular samples, the intima and adventitia of the vessel were annotated using TissueTag. The relative distances between the intima and adventitia were computed by OrganAxis creating a trans-medial gradient.

#### Coronary atherosclerosis Visium data

Visium spatial data from 12 atherosclerothic and healthy coronary arteries was retieved from Bleckwehl et al. and combined with our Visium spatial data from coronary arteries ^48^. To deconvolve cell state composition in Visium spatial transcriptomics sections from atherosclerotic and healthy coronary arteries, we applied cell2location (c2l) using cell state reference signatures derived from our own snRNA-seq reference atlas. Per-spot c2l abundance estimates were normalised by dividing each cell state’s predicted abundance by the sum of all cell state predictions within that spot, yielding a relative compositional measure. Structural regions (intima/plaque, media, adventitia) were annotated manually on each section, and a continuous radial axis coordinate spanning −1 (luminal/intima) to +1 (adventitia) was computed per spot using OrganAxis. This axis was discretised into five equidistant bins (inner, inner-mid, mid, mid-outer, outer). Spots annotated as intima/plaque or media were retained; OCT-embedded samples and sections from intermediate disease stages were excluded. Mean normalised cell state abundance per bin per donor was computed and visualised as line plots across the intima-to-adventitia axis, stratified by disease state (healthy vs. atherosclerosis).

### Visium HD analysis

#### Segmentation of Visium HD bins into cells

Visium HD 2 *µ*m-binned data were reconstructed into single cells with bin2cell ^103^. For each sample, the 2 *µ*m bin output of Space Ranger was read together with the high-resolution H&E image, bins with no counts were removed, and a scaled H&E image was generated at 0.3 *µ*m per pixel. Nuclei were segmented from the H&E image with StarDist ^104^ (2D_versatile_he model), with the probability threshold set to 0.03 after visual inspection across a range of thresholds. Because H&E segmentation can miss cells where nuclei are sparse, faint or atypically shaped, a complementary segmentation was performed on a gene-expression image: total counts per bin were rendered as an image, smoothed with a Gaussian filter (*σ* = 5 pixels) and segmented with StarDist (2D_versatile_fluo model, default probability threshold, non-maximum-suppression threshold 0.5). The H&E nuclear labels were expanded into neighbouring unlabelled bins and used as the primary segmentation, with the expression-derived labels used only to recover cells in regions not covered by an H&E object; the merged labels were then used to aggregate the constituent 2 *µ*m bins into per-cell count profiles.

#### Cell type and state annotation of Visium HD cells

Segmented cells were annotated using the single-nucleus RNA-seq atlas as a reference. A combined reference was assembled from the atlas (downsampled to at most 5,000 cells per cell type) together with, for cell types that could be confidently identified in the Visium HD data, the corresponding Visium HD cells, the latter labelled by marker-gene scoring of a deliberately over-clustered Leiden partition; cell types not expected in the profiled region were excluded. Cell-type composition was estimated for each segmented cell against this reference with TACCO^105^ (optimal-transport annotation with multi_center = 5), and each cell was assigned its single highest-probability cell type; cells with fewer than 100 detected genes were then discarded. The resulting profiles were integrated with scVI^91^ (sample as the batch covariate; total counts and mitochondrial and ribosomal fractions as continuous covariates; 20 latent dimensions), and the latent space was used for nearest-neighbour graph construction, Leiden clustering and UMAP in Scanpy ^87^. Clusters were annotated by canonical marker expression, with smooth muscle cells further resolved into arterial, coronary-artery and arteriolar states. Cell-state labels were transferred from the atlas with CellTypist ^10^. Cells that remained unannotated (for example transcriptionally mixed bins) were labelled by a weighted k-nearest-neighbour scheme combining spatial proximity and expression similarity (30 principal components of the 3,000 most highly variable genes; 15 neighbours in each space; Gaussian distance weighting; blending weight 0.2 in favour of spatial proximity), a label being assigned only when it received at least 40% of the weighted vote.

#### Smooth muscle cell transcriptional programmes

To resolve transcriptional zonation across the vessel wall, a transmural coordinate was defined for the bin2cell-segmented cells of a pulmonary vein sample. The luminal and adventitial surfaces of the wall were annotated on the H&E image at 8 *µ*m-bin resolution and assigned to each segmented cell by majority vote of its constituent 2 *µ*m bins. For every cell, Euclidean distances to the nearest luminal (*d*_lum_) and nearest adventitial (*d*_adv_) landmark cell were computed, and a transmural coordinate was defined as *d*_lum_*/*(*d*_lum_ + *d*_adv_), running from 0 at the lumen to 1 at the adventitia; cells were restricted to the vessel wall by retaining those whose summed landmark distance was below 1.5 times the median summed distance of the annotated landmark cells. Wall smooth muscle cells were ordered along this coordinate and divided into 20 equal-width bins, and pseudobulk profiles were formed as the mean log-normalised expression per bin (counts per 10,000, log(1 + *x*)) over the 2,000 most highly variable genes. Non-negative matrix factorisation was applied to the bin-by-gene pseudobulk matrix with scikit-learn ^92^ (n_components=2, random_state=42, max_iter=1000), yielding per-bin factor weights (normalised to proportions, with the transition point taken as the bin minimising the absolute difference between the two factor proportions) and per-gene factor loadings; per-cell factor scores were obtained by projecting individual cells onto the fitted factors and min–max scaling. The two factors corresponded to a synthetic programme enriched towards the luminal aspect (top-loading genes including *VCAN*, *TNC* and *ENG*) and a contractile programme enriched towards the adventitial aspect (*DES*, *PLA2G2A* and *ACTG2*), the layers being distinguished by the genes with the largest difference in loading between factors. The top-loading genes of each factor were visualised as *z*-scored mean expression across the transmural bins.

### Xenium spatial analysis

The segmentation of transcripts into cells was performed using Baysor. ^9^ Quality control metrics were calculated and cells with < 3 genes or < 25 counts were removed as low quality cells.

#### Xenium cell type annotation

Gene counts were normalised per cell to target sum of 1e4 and log1p transformed. ^106^ Principal component analysis was performed and neighbourhood graph computed followed by Leiden clustering. Clusters were annotated with broad cell type labels based on expression of previously described marker genes. ^1^

#### Cell state label transfer from snRNA-seq to Xenium

To recapitulate the granularity of cellular heterogeneity observed in snRNAseq data and preserve consistency of cell state labels CellTypist was used to transfer cell state labels from snRNAseq ^107^. Annotated snRNAseq object was subset to genes included in the Xenium Spatial Human Multi-Tissue and Cancer panel (377) and used for training the CellTypist models. The training dataset was divided into 8 objects based on the anatomical location corresponding to the anatomical origins of the Xenium samples in the prediction dataset (Ao, LAA, LAD, LV, PV, RAA, SAN, Valve). Subsequently, each regional training object was divided into subsets based on cell type (or group thereof) identified in the Xenium data in the previous step (Adipocytes, EndothelialCells, Pericytes, NeuralCells, MyeloidCells + LymphoidCellsNonB + LymphoidCellsB, Fibroblasts + ValveFibroblasts + EpicardialCells + MastCells, SmoothMuscleCells, LymphaticEndothe-lialCells, AtrialCardiomyocytes + VentricularCardiomyocytes) that were found in the relevant region. Separate CellTypist models were trained from these 62 snRNAseq objects and applied to predict cell state labels in the relevant portion of the Xenium object.

Predicted labels and majority voting approaches from the CellTypist package were applied for cell state label predictions. The predicted labels approach was selected for further analyses as it captured the cell state granularity in rarer cell states. Confidence scores were obtained and cells with conf_score < 0.5 were excluded from further analyses as low quality predictions.

#### Clustering-based Xenium cell state annotation

To explore if new cell states not recovered in the snRNAseq data could be found with Xenium, an alternative cell state annotation was produced using a clustering approach. The cell type annotated object was divided by the cell type cluster annotation. The cell type cluster of Fibroblasts + ValveFibroblasts + EpicardialCells + MastCells was a priori divided by the region of origin to valves and all other regions. Each resultant object was re-integrated using inVAE in a manner consistent with the integration and clustering of snRNAseq data. The parameters used for the inVAE models were n_epochs=200, lr_train=0.001, weight_decay=0.0001, inv_covar_keys_cat = [“tissue”], spur_covar_keys_cat=[’tissue_-block_id’]. To aid annotation in alignment with snRNAseq data, top marker genes of cell states in snRNAseq data were obtained within the restricted space of genes included in the Xenium probe set. Gene scores for these combinations of genes were computed for cells in the Xenium data set. Leiden clustering based on neighbours from inVAE latent representation was undertaken on each object and clusters were annotated based on maker gene expression and marker gene scores derived from snRNAseq cell state labels. This annotation was used to describe Neurons that were not recovered in snRNA-seq data (Supplementary Fig. S2h, i).

#### Niche identification in Xenium spatial data (NicheCompass)

Niches of co-localising cells sharing common gene programs were identified using NicheCompass (v0.2.3), a graph based deep-learning method. ^11^ The default prior Knowledge Gene Program Mask based on OmniPath, MEBOCOST and NicheNet was used as described previously. ^11^ Active gene program threshold ratio 0.2 was used.

Leiden clustering based on NicheCompass latent representation was used to identify niches. We selected the lowest Leiden resolution that reproduced all tissue features a priori known from our Visium Spatial analysis. Niche names were selected on the basis of cell type composition and spatial organisation. Clusters restricted to a single sample with < 30 cells were excluded from further analysis. At the chosen resolution, multiple cardiomyocyte-dominated clusters without meaningful differences in cell type composition were identified. A dendrogram was computed based on correlations in normalised cell state composition of niches. Niches with correlation distance <0.1 were aggregated according to the dendrogram producing the final annotation. The niches were named according to their cell type composition and spatial distribution.

#### Niche composition comparison across modalities

To correlate spatial niches identified across modalities, niche cell-state composition vectors were derived independently for each platform. For Xenium data, niches were defined using NicheCompass and cell-state labels were transferred from matched snRNA-seq data with CellTypist; the proportion of each cell state within each niche was computed by counting labelled cells and row-normalising to sum to one. For Visium data, niches were identified at 55 µm spot resolution and cell-state abundances were estimated using cell2location; raw predicted cell counts were first normalised per spot to compositional fractions (sum to one), and these per-spot vectors were then averaged across all spots belonging to each niche. Given the shared reference for CellTypist and cell2location, both modalities shared a common cell-state vocabulary of 65 cardiac cell states, enabling direct compositional comparison. Pairwise cosine similarity was computed between all Xenium and Visium niche vectors, providing a scale-invariant measure of compositional correspondence that is robust to differences in cell density and quantification method between platforms. Niches were ordered by average-linkage hierarchical clustering on cosine distance for visualisation. Results were displayed as a cosine similarity heatmap.

#### Gene expression imputation (scENVI)

To address the limitation of 377 gene panel of the Xenium Spatial method, environmental variational inference (scENVI v0.4.5) was used to impute the expression of all genes in Xenium cells as described previously. ^108^ Xenium data was divided by region of origin for imputation and snRNAseq data from the corresponding region was used for training.

#### Cell-cell communication in Xenium niches (CellChat)

Xenium data with imputed gene expression was divided into separate objects based on niche and region of origin. CellChat (v2.2.0.9001) was used to predict cell-to-cell signalling between cell states in each region-niche. Distance thresholds of 200 *µ* and 30 *µ* were used for secreted and contact signalling respectively. ^109^ Niches in regions of a priory clinical interest and relevance for cardiovascular disease (e.g. myocardial sleeve of the pulmonary veins) were selected for further interrogation.

#### Xenium 5K cell type annotation

Xenium *in situ* Multimodal Cell Segmentation was used to assign transcripts to cells in the Xenium 5K data. Cell state labels were transferred from snRNAseq data using CellTypist. For each Xenium 5K sample, CellTypist models were trained on snRNAseq data from the corresponding anatomical region using the 5001 genes included in the Xenium 5K panel. ^107^.

Xenium cells were filtered to retain those with a minimum of 25 total counts and at least 3 detected genes before CellTypist prediction was run. A confidence score threshold of >0.2 was applied to retain high-confidence annotations. This procedure was applied separately to two coronary artery Xenium sections and one aortic Xenium section.

### Quantification of macrophages from post-Xenium immunofluorescence

Following Xenium imaging, the same valve sections were stained and imaged by immunofluorescence on a RareCyte platform, producing full-resolution pyramidal OME-TIFF images at 0.325 *µ*m per pixel across a 19-channel panel that included a Hoechst nuclear stain, CD163 (ArgoFluor580L) as a macrophage marker and E-cadherin (ArgoFluor730). For each valve leaflet section, two freehand regions of interest were drawn in Fiji ^82^ on a downsampled pyramid level (level 3, 8× downsampled), tracing the fibrosa and non-fibrosa borders of the leaflet as a labelled pair; five sections were annotated in this way. The paired ROI coordinates were scaled to full resolution and used to extract a multi-channel image crop with 30 *µ*m of padding around each section.

Nuclei were segmented on the full-resolution Hoechst channel of each crop with Cellpose ^110^ using the pre-trained nuclei model (flow_threshold 0.4, cellprob_threshold 0.0). Because the default percentile normalisation (1st–99th) performed poorly on sparsely cellular crops with highly skewed intensity distributions, the nuclear diameter (20, 30 or 40 pixels) and the upper normalisation percentile (99.0, 99.9 or 99.99, with the lower bound fixed at the 1st percentile) were swept, and a diameter of 20 pixels (∼6.5 *µ*m at 0.325 *µ*m per pixel) with an upper normalisation percentile of 99.9 were selected after visual inspection of segmentation quality. For each segmented nucleus, the median intensity of every channel was quantified from the full-resolution pyramid, and the nuclear centroid was recorded as its spatial coordinate. Each nucleus was assigned to its enclosing ROI polygon by point-in-polygon testing of its centroid, with nuclei falling outside both ROIs labelled unassigned.

To place each nucleus along the leaflet, a local fibrosa-to-non-fibrosa coordinate was computed separately for each section from the relative proximity of the nucleus to the two border ROIs. Euclidean distances from each nucleus to the nearest nucleus of one border (*d_A_*) and of the other (*d_B_*) were obtained with KD-trees, and an edge score (*d_A_*−*d_B_*)*/*(*d_A_*+*d_B_*) was calculated, ranging from −1 at one border to +1 at the other; this local distance-ratio formulation was used in preference to a single global axis because the leaflet borders are thin, curved structures that do not align with one linear axis. The fibrosa or non-fibrosa identity of each border was assigned by inspection of the tissue morphology, and the edge scores were oriented accordingly into a common coordinate running from −1 (fibrosa) to +1 (non-fibrosa). One section, corresponding to a valve chorda rather than a leaflet, was excluded, leaving four sections for analysis.

Within each section, nuclei were classified as CD163-positive (macrophages) if their CD163 intensity exceeded the 90th percentile of CD163 intensity for that section; a per-section threshold was used to accommodate differences in staining intensity between sections. Macrophage enrichment towards the non-fibrosa surface was assessed across the four sections by comparing the CD163-positive fraction in the most fibrosa decile (lowest 10% of the coordinate) with that in the most non-fibrosa decile (highest 10%), using a one-sided paired Wilcoxon signed-rank test (*p* = 0.062). In each section, the CD163-positive proportion was additionally compared between the fibrosa and non-fibrosa halves of the coordinate using a one-sided Fisher’s exact test, and these per-section tables were combined with a Cochran–Mantel–Haenszel test stratified by section.

### Gene regulatory network analysis (SCENIC+)

The SCENIC+ pipeline v1.0a2^111^ was used to predict transcription factors, putative target genes and regulatory genomic regions harbouring transcription factor binding sites for each of the 18 cell types in the Multiome atlas. For each analysis, gene expression and chromatin-accessibility peak counts from the paired Multiome data were used as input. Differentially accessible regions were identified using ArchR ^90^ for each annotated cell type. Marker regions for each cell type were identified against the remaining cell types using FDR < 0.1 and absolute log2 fold-change > 0.5, and exported as BED files for downstream SCENIC+ analysis. For each cell type, metacells were created using SEACells v0.3.3^112^ based on the MultiVI latent space ^113^, followed by aggregation of counts and fragments. Metacells supported by fewer than 10 nuclei were removed before downstream analysis. The cisTopic framework ^114^ was applied to identify region topics from chromatin-accessibility fragment counts. Topic-derived regions and ArchR-derived differentially accessible regions were used as candidate regions for transcription factor binding. Motif enrichment was then performed using pycisTarget with cisTarget and Differential Enrichment of Motifs against ENCODE-v4 motif databases. Enriched motifs were assigned to transcription factors using the v10nr_clust HGNC annotation table, retaining both direct and orthology-extended transcription factor assignments. Gene regulatory networks were assembled using the SCENIC+ eGRN inference workflow. Gene expression and chromatin-accessibility matrices were first aligned into a unified MuData object. A transcription start site-centred search space spanning 1–150 kb upstream and downstream of each gene was used to link candidate regulatory regions to genes. Region-to-gene and transcription factor-to-gene links were inferred using gradient-boosted regression, with Spearman correlation used for region-to-gene associations. Direct and extended eRegulons were constructed by linking transcription factors, regulatory regions and target genes across the inferred cistromes, region-to-gene links and transcription factor-to-gene links. Regulon activity scores were computed with AUCell based on target-gene expression and target-region accessibility. The final SCENIC+ outputs included direct and extended eRegulons, AUCell activity matrices, and region-to-gene and transcription factor-to-gene adjacency tables. These outputs were assembled into a unified SCENIC+ MuData (using Muon ^115^) object for downstream analysis and visualisation.

### eRegulon activity analysis in cardiomyocyte metacells

Cell-state specific eRegulon activity was analysed in ventricular and atrial cardiomyocyte metacells. For each metacell, direct gene-based and direct region-based AUCell scores were used to quantify the transcriptional and chromatin-accessibility components of each eRegulon, respectively. Metacells were assigned to cardiomyocyte cell states based on their dominant cell-state label. To reduce ambiguity from mixed-state metacells, only metacells with cell-state assignment purity ≥ 0.70 were retained. Cell states represented by fewer than three purified metacells were excluded from downstream testing. LeftStressed was excluded because only one purified metacell was available.

Cell-state specific eRegulons were required to pass all of the following criteria: (1) the corresponding transcription factor was expressed in at least 10% of metacells in at least one cell state; (2) gene-based and region-based AUCell scores showed a positive correlation across metacells, indicating consistency between the transcriptional and enhancer-accessibility components of the regulon; (3) the eRegulon ranked within the top 30 by regulon-specificity score in at least one state; and (4) high-AUCell metacells were contributed by at least two distinct donors, used as a proxy for donor robustness. Among quality-passing eRegulons, state-associated activity was assessed using a one-vs-rest Mann–Whitney U test. Multiple testing was controlled using the Benjamini–Hochberg false discovery rate. For visualisation, the top quality-passing eRegulons per state were selected based on statistical ranking and biological relevance.

### GWAS summary statistics and fine-mapping

Fine-mapped genome-wide association study (GWAS) variants were obtained from the Open Targets Platform ^116^. We selected studies mapped to the trait ontology identifiers for atrial fibrillation (AF; EFO_0000275), coronary artery disease (CAD; EFO_0001645), and aortic valve stenosis (AVS; EFO_0000266). For each trait, statistically fine-mapped 95% credible sets were retrieved together with variant posterior inclusion probabilities (PIPs). Variants were represented on GRCh38 coordinates and harmonised by genomic position and reference/alternative allele.

For downstream regulatory analyses, we retained credible-set variants with PIP *>* 0.4. This yielded 435 AF variants, 212 CAD variants, and 8 AVS variants. For analyses requiring allelic read assignment, variants were further restricted to biallelic single-nucleotide variants overlapping the consensus snATAC-seq peak set of the atlas, thereby selecting disease-associated variants located within accessible chromatin. The resulting trait-specific variant sets were merged into a single non-redundant set of 655 candidate SNPs for allele-specific accessibility analysis. Population allele frequencies were annotated from gnomAD ^117^ through the myvariant.info service.

For SNP2CELL analyses, the same PIP-filtered fine-mapped variants were intersected with peaks in the healthy adult heart enhancer gene regulatory network (eGRN) derived from SCENIC+ analyses. SNPs overlapping eGRN peaks were assigned to the corresponding regulatory regions, and peak-level genetic scores were calculated as the sum of overlapping SNP PIPs for each trait. These trait-specific peak scores were subsequently propagated through the eGRN as described below.

### SNP2CELL analysis

We constructed a healthy adult heart eGRN from SCENIC+ analyses performed independently on 18 cell type. Fine-mapped GWAS variants were obtained from the Open targets, and credible-set SNPs with posterior inclusion probabilities over 0.4 were retained for atrial fibrillation (435 SNPs), coronary artery disease (212 SNPs) and aortic valve stenosis (8 SNPs). SNPs were mapped to overlapping eGRN peaks, and peak-level genetic scores were calculated as the sum of overlapping SNP PIPs.

Trait-specific GWAS scores were propagated across the eGRN using SNP2CELL v0.3.0 with personalised PageRank and 1,000 score-shuffled permutations. ^19^ Group-wise differential-expression scores were computed on the matched healthy heart Multiome object at fine cell-state levels using SNP2CELL. GWAS-propagated and differential-expression scores were then integrated by taking the inner minimum of their robust z-scores for each gene-group pair, prioritising genes jointly supported by trait genetics and cell-state specific expression.

### Allele-specific accessibility analysis

#### Variant selection and donor genotyping

Candidate variants for allele-specific accessibility testing were drawn from statistically fine-mapped GWAS credible sets in the Open Targets Platform ^116^. Studies were selected by matching their mapped trait ontology identifiers to atrial fibrillation (EFO_0000275), coronary artery disease (EFO_0001645) and calcific aortic valve stenosis (EFO_0000266), and the variants in each 95% credible set were retained together with their posterior inclusion probabilities. Across the three traits, variants were filtered to biallelic single-nucleotide variants with a posterior inclusion probability above 0.4 that overlapped the consensus snATAC-seq peak set of the atlas (i.e. lay within accessible chromatin), and were merged into a single non-redundant set of 655 candidate SNPs. Population allele frequencies were annotated from gnomAD ^117^ through the myvariant.info service.

Donor genotypes at these SNPs were determined across 26 donors by two complementary approaches. For 15 donors with array data, genotypes were taken from a genome-wide SNP-genotyping array (Affymetrix Axiom) in GRCh38 coordinates and exported as a single-sample VCF per donor. For the remaining 11 donors, and to benchmark the array calls, genotypes were inferred directly from the sequencing reads: bases at each SNP were piled up from the pooled snRNA-seq and snATAC reads (minimum base and mapping quality 20) and a genotype was called where at least 30 reads were available — homozygous reference at an alternate-allele fraction ≤ 0.20, homozygous alternate at ≥ 0.80, and heterozygous otherwise. Array genotypes took precedence where available. Read-inferred genotypes were concordant with the array calls (approximately 96% across the 15 array-genotyped donors, pooling both modalities), and the resulting genotype matrix was used to identify, for each SNP, the donors heterozygous at that site.

#### Allelic remapping and per-barcode allele counting

To prepare single-nucleus reads for allelic analysis, per-sample BAM files were harmonised to a common, globally unique cell identifier: each read’s cell-barcode (CB) tag was rewritten to match the corresponding nucleus name in the integrated count and peak matrices, prepending the sample or Multiome-library identifier to avoid barcode collisions across samples; reads lacking a CB tag were discarded; and contig names were standardised to the chr-prefixed GRCh38 convention. Harmonised sample-level BAMs were merged into one alignment file per donor and modality with SAMtools ^118^ (snRNA-seq for 26 donors, snATAC for 19), retaining only reads from quality-control-passing, annotated nuclei via a per-donor barcode whitelist.

Reference-mapping bias was removed with WASP^54^. For each donor, reads overlapping one of that donor’s heterozygous SNPs were identified, re-mapped after swapping the allele at the overlapping site, using STAR^119^ for snRNA-seq and BWA-MEM ^120^ for snATAC reads, with snATAC mate pairs processed as independent single-end reads, and discarded unless they re-mapped to their original position. Reads passing this filter were merged with reads that did not overlap any heterozygous site. WASP removed a small fraction of reads (on average 2.8% of snATAC and 9.3% of snRNA-seq reads per donor).

Because WASP re-mapping does not preserve cell-barcode tags, alleles were counted from the original CB-tagged BAMs, restricted to the reads that had passed WASP filtering (matched by read name). At each SNP, bases were piled up (minimum base and mapping quality 20) and assigned to the reference or alternate allele per cell barcode. snRNA-seq counts were UMI-deduplicated, reads sharing a cell barcode and UMI were collapsed to a single allele call by majority vote, discarding ties, while snATAC counts were taken at the read level, yielding per-barcode reference and alternate counts at each SNP.

#### Beta-binomial test for allele-specific accessibility

Per-barcode allele counts were aggregated to donor-level pseudobulk at three resolutions using the atlas annotation: across all nuclei of a donor (bulk), and within each of the 18 cell types and 65 cell states. At every SNP only donors heterozygous at that site contributed (487 testable heterozygous SNPs, on average 4.8 heterozygous donors per SNP). For each SNP × group × modality, allelic imbalance was tested only where at least four heterozygous donors had at least 10 reads each and the imbalance was directionally consistent across donors — at least 70% of those donors showing the same direction of allelic bias as the pooled estimate; combinations failing these criteria were not tested. Imbalance was assessed with a beta-binomial likelihood-ratio test comparing a model with a free mean allelic ratio and overdispersion against a null in which the mean ratio was fixed at 0.5 (a binomial test against 0.5 was also recorded). P values were corrected within each group × modality (Benjamini–Hochberg), and associations with a false-discovery rate below 0.15 were considered significant. Tests on chromatin accessibility (snATAC) define allele-specific accessibility and the equivalent tests on expression (snRNA-seq) allele-specific expression; running the test at all three resolutions allowed allelic effects seen in bulk to be resolved to individual cell types and states.

### Variant effect prediction with deep-learning sequence models

#### Fine-tuning ChromBPNet

To model cell-type-resolved chromatin accessibility at base resolution, we trained ChromBPNet models ^56^ on pseudobulk snATAC-seq profiles for each cell type, using a pipeline adapted from the Human Development Multiomic Atlas (HDMA) ^121^ (https://github.com/GreenleafLab/HDMA). For each cell type, fragments from all quality-control-passing, annotated nuclei were extracted from the ArchR project ^90^, pooled across samples and coordinate-sorted to yield a single pseudobulk fragment file. All steps used the GRCh38 reference genome (10x Genomics Cell Ranger ARC reference, 2020-A).

Accessible regions were called from the pseudobulk fragments with MACS2^122^ (callpeak -f BED -g hs -p 0.01 –nomodel –shift −75 –extsize 150 –keep-dup all –call-summits -B –SPMR). Peaks were ranked by significance, the top 300,000 retained, and ENCODE blacklist (v2) regions removed with BEDTools ^123^; each retained summit was then extended to a fixed 1,000 bp window centred on the summit to define the final peak set. GC-matched background (non-peak) regions were generated per fold with chrombpnet prep nonpeaks (input length 2,114 bp, stride 1,000 bp), excluding blacklist regions. Models were trained and evaluated by five-fold cross-validation across chromosomes, each fold holding out a distinct set of test chromosomes (fold 0: chr1, chr3, chr6; fold 1: chr2, chr8, chr9, chr16; fold 2: chr4, chr11, chr12, chr15, chrY; fold 3: chr5, chr10, chr14, chr18, chr20, chr22; fold 4: chr7, chr13, chr17, chr19, chr21, chrX) and a separate pair of validation chromosomes, with the remaining chromosomes used for training.

Tn5 enzymatic bias was modelled once and shared across all cell types: a bias model was trained with chrombpnet bias pipeline (–data-type ATAC, –bias-threshold-factor 0.4) on the cell type with the greatest number of fragments (Ventricular Cardiomyocytes), using its peak set, GC-matched negatives and fold-0 chromosome split, with fragments subsampled to 50 million where this cap was exceeded. For each target cell type, ChromBPNet accessibility models were then trained per fold with chrombpnet pipeline (–data-type ATAC) from that cell type’s own fragments and peak set together with the shared, frozen bias model, again subsampling fragments to a maximum of 50 million, yielding five bias-factorised accessibility models (input length 2,114 bp; output length 1,000 bp) per cell type.

To interpret the trained models, per-base contribution scores for the total-count (accessibility) output head were computed for up to 150,000 top-ranked peaks per fold with DeepSHAP (chrombpnet contribs_bw) and averaged across the five folds. *De novo* motifs were discovered from the averaged contribution scores with TF-MoDISco ^124^ (sliding_-window_size 20, flank_size 5, target_seqlet_fdr 0.05, max_seqlets_per_metacluster 10^5^, n_leiden_runs 2) and annotated by matching to the JASPAR 2024 CORE vertebrates (non-redundant) database ^101^ with TOMTOM ^125^.

Cell-type-matched ChromBPNet models outperformed both a Tn5-bias-only baseline and mismatched (cross-cell-type) models in variant-effect prediction; the variant-scoring procedure and benchmark are described below.

#### Variant scoring with ChromBPNet

Variant effects on chromatin accessibility were predicted with the cell-type-matched, bias-factorised ChromBPNet models (the bias-corrected chrombpnet_nobias model) ^56^. For each SNP, a 2,114 bp reference sequence centred on the variant was extracted from GRCh38 and the reference and alternate alleles were substituted at the central position, after confirming that the reference base matched the genome. Each allele was scored with all five cross-validation fold models, predicting both the per-base accessibility profile and the total log-count (coverage), with predictions computed on the forward and reverse-complement strands and averaged. The predicted allelic effect was summarised as the log_2_ fold-change in predicted total counts between the alternate and reference alleles, and the predicted change in profile shape as the Jensen–Shannon divergence between the allelic profiles, both averaged across folds.

To benchmark predictions against observed allele-specific accessibility, the predicted alternate-allele fraction (predicted alternate counts divided by the sum of predicted reference and alternate counts) was compared with the observed pooled alternate-allele fraction for each significant ASA SNP–cell-type pair (**Fig. 5d**), and agreement was quantified by Pearson and Spearman correlation and by directional concordance (the fraction of pairs in which the predicted and observed alternate-allele fractions fell on the same side of 0.5, i.e. agreed on the more accessible allele). To assess the contribution of cell-type-matched training, the same scoring and comparison were repeated using the cell-type-matched model, models trained on other (mismatched) cell types, and the Tn5-bias model alone (**Supplementary Fig. S5g, h**).

To place each variant within its local *cis*-regulatory context, all common variants (gnomAD v4.1^117^, minor allele frequency ≥ 1%) within a 50 kb window of the index SNP were scored identically, and linkage disequilibrium (*r*^2^) with the index SNP was computed from 1000 Genomes Phase 3 European samples ^85^; these supported the locus visualisations.

#### Nucleotide contribution and motif attribution

Per-base contribution scores were computed for the reference and alternate allele sequences by integrated gradients ^126^ applied to the total-count (accessibility) output head of the cell-type-matched ChromBPNet models. For each sequence, gradients were integrated over 50 steps along the linear path from each baseline to the observed sequence using 20 dinucleotide-shuffled baselines, computed on the forward and reverse-complement strands and then averaged across both orientations and the five fold models. Both the raw attribution and the projected attribution (attribution multiplied by the one-hot sequence) were retained for each allele.

To identify the transcription-factor motif affected by each variant, a query motif was constructed from the projected attribution in a 20 bp window on either side of the variant: positive attribution values were normalised per position to a probability matrix and trimmed to the region of highest information content around the variant, weighted by the magnitude of the reference-versus-alternate attribution difference, to give a compact (8–25 bp) position-weight matrix. This query was matched against a MEME-format motif database with TOMTOM ^125^ (-no-ssc, minimum overlap 5, E-value scoring), retaining only matches whose aligned region overlapped the variant position and excluding yeast (YeTFaSCo) motifs; significant matches (E-value *<* 0.05) were ranked by TOMTOM *P* value, and the expression of the matched transcription factors was retrieved per cell type from the single-nucleus RNA-seq atlas. This identified creation of a MEF2-family motif by the alternate allele of rs2979489 (*P* = 2.7 × 10^−7^) and disruption of a CREB1-like motif by the alternate allele of rs1706003 (*P* = 2.8 × 10^−6^).

#### Fine-tuning AlphaGenome

We used a pytorch version of AlphaGenome for the fine tuning. Single-nucleus ATAC-seq and RNA-seq profiles from the heart atlas were partitioned into 65 cell states. For each cell state and modality, per-base coverage tracks were generated and converted to bigWig format using snapatac-scooby ^78^. RNA-seq tracks were generated separately for the positive and negative strands (gex_plus and gex_minus). Each track was CPM-normalised with interval-size weighting. For each normalised track, we then computed the non-zero mean, defined as the length-weighted mean coverage across non-zero intervals. This value was used as the precomputed track-normalisation statistic during fine-tuning.

To improve the efficiency of our fine-tuning and encourage the model to discriminate between cell states in our dataset, we identified cell-state-specific marker peaks using ArchR differential accessibility analysis and aggregated across the 65 heart cell states. For each AlphaGenome fold-0 interval, the number of overlapping marker peaks was counted, and intervals containing ≥ 10 marker peaks were retained as informative regions. This identified 18,462 informative intervals in the TRAIN partition and 2,716 in the VALID partition.

Training, validation and test sets were drawn from the disjoint TRAIN, VALID and TEST subsets of the AlphaGenome fold-0 genome partition using GRCh38 reference coordinates. The training set included all 41,694 TRAIN intervals plus three additional copies of the 18,462 informative TRAIN intervals, resulting in 97,080 intervals and 4× representation of informative regions in the shuffled training stream to facilitate learning from sparse cell-state-specific marker-peak signals. The validation set comprised the 2,716 VALID intervals overlapping ≥ 10 marker peaks.

The held-out test set comprised independent ATAC and RNA components. The ATAC evaluation set consisted of a 10% random subsample of heart ATAC peaks located within fold-0 TEST intervals. Peaks were excluded if they lay within ±524 kb of any training interval or if a full model-input window could not be generated because of chromosome-boundary constraints, yielding 9,477 test peaks. The RNA evaluation set comprised protein-coding genes located within fold-0 TEST intervals, selected using a maximum per-cell CPM ≥ 10 and an across-cell coefficient of variation ≥ 0.5 as a proxy for cell-state-specific expression. The peak and gene sets were used for ATAC and RNA-seq evaluation, respectively.

Starting from the publicly released AlphaGenome weights, we trained two parameter-efficient adaptation strategies in parallel. In the LoRA16 model, rank-16 low-rank adaptation modules with *α* = 16 were injected into the transformer query and value projections, with an input sequence length of 524,288 bp. In the linear model, the per-modality output heads were trained, while all backbone parameters were frozen; this model used a 1,048,576-bp input sequence.

Both models produced 1-bp-resolution predictions for ATAC and stranded RNA-seq tracks. The ATAC and RNA-seq modalities were optimised with equal weights using the AlphaGenome composite multinomial loss, with positional and count loss weights of 5.0 and 1.0, respectively. Models were optimised using AdamW with a learning rate of 5 × 10^−5^, weight decay of 0.1, cosine learning-rate scheduling with 500 warm-up steps, maximum gradient norm of 1.0, batch size of 1 and gradient accumulation of 1. Checkpoints and held-out test-set losses were saved every 10,000 training steps.

#### Variant scoring with AlphaGenome

Variant effects were predicted using the fine-tuned LinProb AlphaGenome model. Chromatin-accessibility effects were quantified with CenterMaskScorer(output=atac, width=501, agg=diff_log2_sum, res=1bp), which sums the predicted alternative-versus-reference log2 fold-change over a 501-bp window centred on each variant. Gene-expression effects were quantified with GeneMaskLFCScorer(output=rna_seq, mode=body, res=1bp), which integrates the predicted alternative-versus-reference log-fold change across protein-coding gene bodies within the scored interval. Predictions were averaged over forward and reverse-complement input orientations, and the reported raw score was defined as the mean of the orientation-specific scores.

Raw scores were aggregated into variant-by-cell-state effect matrices separately for ATAC and RNA. For ATAC, the absolute raw score was averaged across accessibility tracks assigned to the same cell state. For RNA, strand-aware filtering was applied before aggregation: gex_plus tracks were retained for positive-strand genes and gex_minus tracks for negative-strand genes. The RNA effect for each variant–cell-state pair was then defined as the maximum absolute score across strand-matched protein-coding genes within the scored interval. For each trait–cell-state pair, ATAC and RNA effects were summarised by averaging the absolute raw scores across variants assigned to that trait.

### CardioSleuth: interactive viewer for gene regulatory networks

An interactive web-based viewer, CardioSleuth, was developed to support exploration of cell-state-resolved trait–network results. The viewer displays SNP, transcription factor and gene nodes within cell-state-specific enhancer–gene regulatory networks and integrates precomputed quantitative priors, including per-trait SNP2CELL propagated scores, differential-expression statistics, SCENIC+ network attributes and AlphaGenome-based variant-effect predictions.

A large language model-assisted module was integrated to support user research questions using the heart atlas-derived gene regulatory networks and associated quantitative priors as the primary knowledge base. Natural-language queries were mapped to the active disease, cell-state and gene context, enabling context-aware network visualisation and interpretation of regulatory and genetic relationships. For literature-supported statements, PubMed was queried in real time, and candidate references were verified by internal retrieval functions before citation to reduce unsupported claims. Retrieved references were stored with disease, cell-state and gene metadata, and user feedback was used to support prospective construction of context-specific credible citation sets.

### RUNX1 manipulation in cultured human VSMCs

Human vascular smooth muscle cells (hVSMCs) were isolated from aortic tissue obtained from patients undergoing aortic valve replacement after informed consent (using protocols approved by Cambridge or Huntingdon Research Ethical Committees, conforming to the principles outlined in Declaration of Helsinki). Cells were isolated as described ^127^ and cultured in smooth muscle cell growth medium (SMC-GM2, C22062; PromoCell) supplemented with penicillin–streptomycin under standard conditions (37C, 5% CO2).

For knock-down experiments, cells were reverse transfected with 25nM RUNX1-targeting siRNA (RUNX1 HSS189454 and RUNX1 HSS189453; Thermo Fisher Scientific) or non-targeting control (NTC; Dharmacon, D-001810-10-20) using 1% (v/v) Lipofectamine RNAiMAX (13778030; Invitrogen). A RUNX1 lentiviral vector was generated by cloning RUNX1 cDNA (OHu26363C, GenScript) into a pGenLenti mCherry backbone using Gibson assembly. For overexpression of RUNX1, cells were transduced with lentivirus including RUNX1, or the empty vector, in culture medium supplemented with 10 µg/mL protamine sulfate (P3369; Sigma-Aldrich).

Primary hVSMCs from four donors (2 male, 2 female, passages 1–2) were used for single cell RNA sequencing experiments. Cells were harvested 72 hours after transfection/transduction and flow cytometry-assisted cell sorting (Influx) used to isolate 7-AAD negative, live cells for si-RNA transfected cells, and mCherry-positive for lentivirus-transduced cells. Cells were immediately loaded for cDNA library preparation (On Chip Multiplexing, 10x Genomics) that were sequenced on a NovaSeqX (Illumina).

#### Custom transgene reference and read mapping

To quantify transgene expression directly from the single-cell data, a custom reference was built by adding a single contig representing the transcribed lentiviral cassette to the GRCh38 reference (refdata-gex-GRCh38-2024-A, 10x Genomics). The overexpression construct (RUNX1c–IRES–mCherry) and the empty-vector construct (mCherry only) were supplied as circular SnapGene plasmid maps, and the transcribed cassette was extracted programmat-ically with snapgene_reader. Because the cassette spans the plasmid origin, the linear contig was reconstructed by concatenating the sequence downstream of the CMV enhancer/promoter with the segment preceding it, giving a 4,179 bp contig (RUNX1_OE_transgene) with the structure CMV promoter–RUNX1c CDS–FLAG–IRES–mCherry–WPRE–polyadenylation signal. A single contig was used to represent both constructs because the IRES–mCherry–WPRE segment is identical between the overexpression and empty-vector plasmids, which avoids spurious multimapping of reporter-derived reads. Two features were annotated on the contig: RUNX1-transgene, spanning the 1,442 bp RUNX1c coding sequence, and mCherry, spanning the mCherry coding sequence extended through the downstream WPRE element to capture 3*^′^*-biased reads. Coordinates were validated by confirming the expected start and stop codons of each coding sequence and the identity of the shared mCherry–WPRE region between constructs. The transgene sequence and annotation were concatenated with the GRCh38 FASTA and GTF and the combined reference was built with cellranger mkref (Cell Ranger v9.0.1, 10x Genomics).

Sequencing reads were aligned and quantified with cellranger multi (v9.0.1) against the transgene-augmented reference. Each run corresponded to one GEM well containing cells pooled from two donors (one male and one female), and the four experimental conditions within a well, non-targeting siRNA control, RUNX1 siRNA knockdown, empty-vector control and RUNX1 overexpression, were distinguished by their 10x On-Chip Multiplexing (OCM) barcodes. Intronic reads were excluded (include-introns=false), restricting quantification to spliced transcripts, and position-sorted BAM files were retained for inspection of transgene pile-ups.

Within each OCM-demultiplexed sample, the two pooled donors were separated by genotype using souporcell ^84^ run with two clusters; singlets were retained while cross-donor doublets and unassigned barcodes were discarded. Genotype clusters were matched to individual donors using sex-specific expression, scoring each cell for XIST and a panel of Y-chromosome genes (RPS4Y1, EIF1AY, DDX3Y, KDM5D, UTY, USP9Y and ZFY) and labelling each cluster as male or female accordingly; these assignments were concordant with the souporcell genotypes. Ambient and background RNA were then removed from each library with CellBender remove-background ^86^, and the denoised, donor-resolved single-cell matrices were taken forward for analysis.

#### Single-cell RNA-seq analysis of RUNX1 perturbation

The CellBender-denoised, souporcell-singlet matrices from the eight OCM libraries were concatenated. Because souporcell removes only cross-donor doublets, within-donor doublets were flagged per library with Scrublet ^88^ (expected doublet rate 0.06) and cells with a doublet score above 0.3 were removed. Quality-control metrics were computed in Scanpy ^87^, and cells were retained if they had between 1,000 and 10,000 detected genes, at least 2,000 UMIs and fewer than 5% mitochondrial counts. Counts were normalised to 10,000 per cell and log-transformed, and the two empty-vector labels used across the GEM wells were merged into a single empty-vector control condition.

A scVI^91^ latent representation was used as the basis for a nearest-neighbour graph, Leiden clustering (resolution 0.8) and UMAP visualisation. Endogenous RUNX1, the RUNX1 transgene and mCherry were quantified per cell from the custom reference, with endogenous RUNX1 distinguished from the transgene by its Ensembl gene identifier. For the overexpression and empty-vector conditions, successfully transduced cells were defined as those with detectable mCherry (UMI *>* 0). Four groups were used for comparison: non-targeting control, RUNX1 siRNA knockdown, mCherry^+^ empty-vector control and mCherry^+^ RUNX1 overexpression. Knockdown was assessed against the non-targeting control across all cells, whereas overexpression was assessed between mCherry^+^ overexpression and mCherry^+^ empty-vector cells so that only transduced cells were compared.

Cell-cycle phase was assigned with score_genes_cell_cycle in Scanpy using canonical S-phase and G2/M marker gene sets ^53^, and cells in S or G2/M phase were classed as cycling. The cycling fraction was summarised per donor for each group and compared between paired conditions (knockdown versus non-targeting control, and overexpression versus empty vector) using paired t-tests across donors.

Differential expression was performed on donor-level pseudobulk profiles. For each contrast, counts were summed within each donor–condition combination (pseudobulk samples with fewer than 10 cells were excluded) and tested with PyDESeq2^128^ via pertpy ^96^, using condition as the design variable. Genes with a Benjamini–Hochberg-adjusted P value below 0.05 and an absolute log_2_ fold-change above 0.5 were considered differentially expressed. The knockdown and overexpression contrasts were then compared, and genes significant in both with opposite directions of change were defined as concordant RUNX1 targets. Functional enrichment of the up-regulated, down-regulated and concordant gene sets was assessed with g:Profiler ^129^ against GO, KEGG, Reactome and WikiPathways, with significance at an adjusted P value below 0.05. To confirm that the transcriptional changes were not driven solely by proliferation, the differential-expression and enrichment analyses were repeated after removing the cell-cycle genes used for phase scoring.

To distinguish switch-like from dose-dependent responses to RUNX1 overexpression, transgene dosage was approximated by mCherry level within mCherry^+^ overexpression cells. The 3,000 most highly variable genes were selected on raw counts (seurat_v3), and as a pragmatic dose-trend screen their log-normalised expression was modelled by ordinary least squares as a function of mCherry level with donor as a covariate, with P values for the mCherry coefficient corrected across genes (Benjamini–Hochberg). Differentially expressed genes with a significant mCherry coefficient (adjusted P *<* 0.05) were classed as dose-responsive and the remainder as switch-like (differentially expressed but lacking a significant dose relationship); this category is therefore defined by the absence of a detectable dose effect rather than by a positive test for thresholding. In parallel, the relationship between transgene dosage and proliferation was examined by binning mCherry^+^ overexpression cells into mCherry quartiles and quantifying the association between per-donor cycling fraction and quartile rank by Spearman correlation.

Finally, the smooth-muscle-cell gene modules defined in vivo by Hotspot ^93^ (module membership at FDR *<* 10^−8^) were scored per cell in the perturbation data with score_genes, and per-donor mean module scores were compared between paired conditions (overexpression versus empty vector, and knockdown versus non-targeting control) using paired t-tests; the coronary-artery smooth-muscle-cell module containing RUNX1 was examined in particular.

## Competing interests

S.A.T. is a scientific advisory board member of Bioptimus, ForeSite Labs, and Xaira Therapeutics; a co-founder, Board observer and equity holder of TransitionBio; a co-founder, consultant and Board Director of Ensocell Therapeutics; a non-executive director of 10x Genomics; and a part-time employee of GlaxoSmithKline. The remaining authors declare no competing interests.

## Data availability

All raw sequencing data will be publicly available at the time of publication. Processed data (sc/snRNA-seq/ATAC-seq, Visium, Xenium) and fine-tuned sequence-to-function models (ChromBPNet and AlphaGenome) will be available from the Heart Cell Atlas (https://www.heartcellatlas.org/v3) at the time of publication. The external atherosclerosis Visium data from Bleckwehl *et al.* ^48^ is available from https://zenodo.org/records/14007461.

## Code availability

CardioSleuth will be publicly available at https://www.heartcellatlas.org/cardiosleuth/ at the time of publication. The code for the other analyses performed in this study is available at GitHub (https://github.com/Teichlab/hypersampling).

## Author contributions

Project design: J.C., L.M., K.K., M.N. and S.A.T. Data generation (tissue procurement): J.C., L.M., L.W., J.D., R.C. and M.N. Data generation (tissue dissection): J.C. and L.M. Data generation (tissue sampling): J.C., L.M., K.K., A.M.A.M., S.N.B. and M.N. Data generation (multiome): J.C., K.K., K.R., L.R., C.S. and R.K. Data generation (Visium): J.C., A.W.- C. and S.Pe. Data generation (Xenium): J.C., A.W.-C. and S.Pe. Data generation (immunofluorescence): L.M., B.T., A.W.-C. and M.N. Data generation (RUNX1 perturbation): J.Z. and H.J. Data generation (3D heart imaging): J.B. Data mapping: K.P. Data analysis (sc/snRNA-seq and multiome): J.C., L.M., T.J., K.K. and S.K. Data analysis (Visium): J.C., L.M. and K.K. Data analysis (Xenium): J.C., L.M., K.K. and Y.X.Z. Data analysis (genetics): I.S. Data analysis (vessels): A.-M.C. Structural annotation: J.C., L.M., S.Y.H., J.L.R. and M.M. Cell type annotation: J.P. Data analysis (RUNX1 perturbation): J.C. and H.J. Data analysis (SCENIC+): W.L. Data analysis (sequence-to-function models): J.C. and T.J. CardioSleuth: T.J. Writing: J.C., L.M., T.J., M.N. and S.A.T. Review and editing: Y.X.Z. and S.Pe. Supervision: J.D., S.Pr., M.N. and S.A.T.

## Acknowledgements

We thank the donors and their families for providing cardiac tissues for research. We are grateful to the staff of NHS Blood and Transplant and the Cambridge Biorepository for Translational Medicine (CBTM) for their support. We thank Ursula Herbort-Brand (National Heart and Lung Institute, Imperial College London, UK) for her immunofluorescence microscopy work contributing to this manuscript, and Professor Peter Lee (ESRF) for providing the HiP-CT heart image. We also acknowledge the staff of the Facility for Imaging by Light Microscopy (FILM) and Connor Preston from the Research Histology Facility at the National Heart and Lung Institute, Imperial College London, for their technical support. We thank the Wellcome Sanger Institute Cytometry and Cellular Generation and Phenotyping (CGaP) teams for their contributions to data generation. We are grateful to the Wellcome Sanger Institute Cellular Genetics Informatics team—particularly Batuhan Cakir, Stanislaw Makarchuk, and Martin Prete—for their support in establishing the Heart-CellAtlas.org webportal. We also thank the Core DNA Pipelines team, the Cellular Generation and Phenotyping (CGaP) team, and the Imperial College Healthcare NHS Trust Tissue Bank for their contributions to this work. NHS Blood and Transplant provided Relevant Material in support of this research. The views expressed in this publication are those of the authors and not necessarily those of NHS Blood and Transplant. The illustration in Fig. 4j was created using BioRender (https://biorender.com).

This project was made possible in part by the Wellcome Trust (WT206194, S.A.T); the Chan Zuckerberg Foundation (2019-002431 and 2019-202666, 2021-237882 to M.N. and S.A.T.); the British Heart Foundation and Deutsches Zentrum für Herz-Kreislauf-Forschung (BHF/DZHK: SP/19/1/34461 to M.N. and S.A.T.); the NIHR Imperial Biomedical Research Centre (BRC) funding to R.A.C. and M.N. This project has received funding from the Wellcome Trust Clinical PhD Fellowship to J.C.; National Institute of health Research (NIHR) Clinical Lectureship to J.C.; a British Society for Heart Failure Research Fellowship to L.M.; a British Heart Foundation Clinical Research Training Fellowship (FS/CRTF/23/24444) to L.M.; a Rosetrees Trust Intermediate Project Grant (PGS23/100028) to L.M, S.K.P. and M.N., a British Medical Association Foundation Josephine Lansdell Grant to L.M. a Heart Research UK Translational Research Project Grant to L.M., S.K.P. and M.N.; the British Heart Foundation Imperial Centre of Research Excellence Award - RE/24/130023; Alexander Jansons Myocarditis UK grant to S.K.P.; Takeda Science Foundation Fellowship, Fukuda Foundation for Medical Technology Fellowship, The Cell Science Research Foundation Fellowship, and The Nakatomi Foundation Fellowship to T.J.; EMBO Postdoctoral Fellowship (ALTF 769-2022) and UK Research and Innovation through guaranteed funding for a Marie Skłodowska-Curie Postdoctoral Fellowship (EP/Z003326/1) to I.S.; the Overseas Research Fellowship of the Takeda Science Foundation to K.K.; a National Heart and Lung Institute PhD studentship to S.N.B.; The Pathological Society of Great Britain & Ireland and The Jean Shanks Foundation Fellowship to M.M.

## Supplementary Figures

**Figure S1: Supplementary Figure S1:**
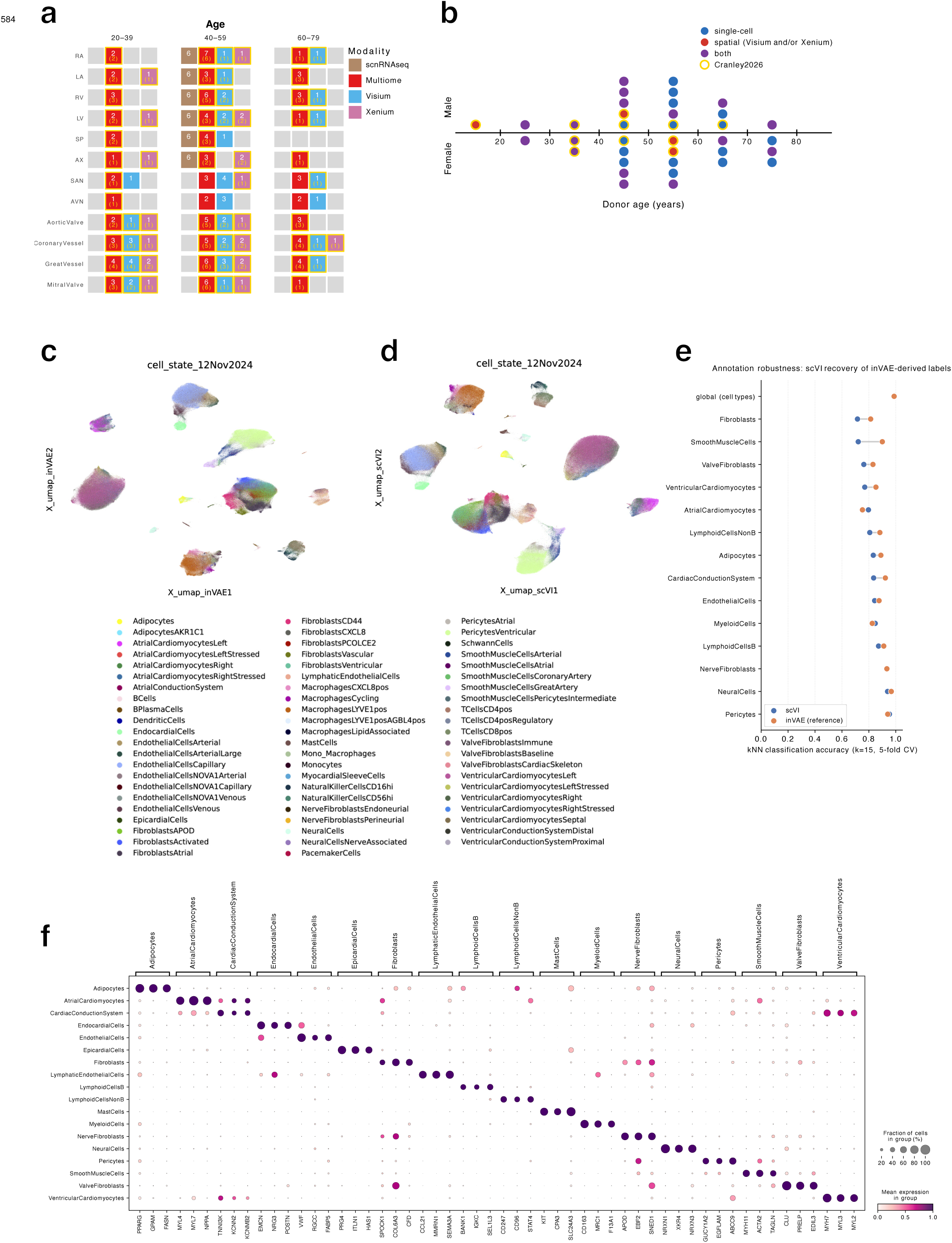
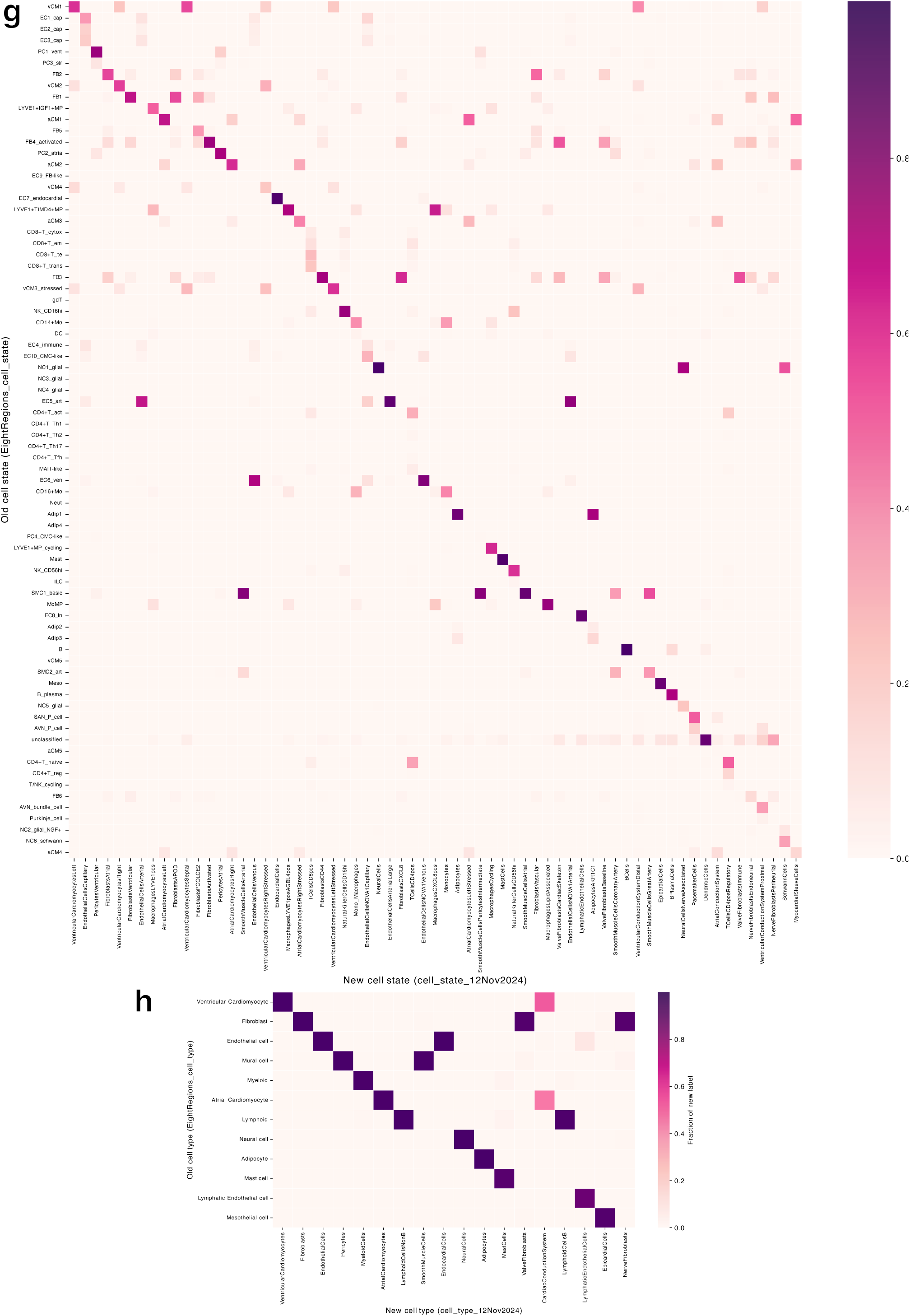
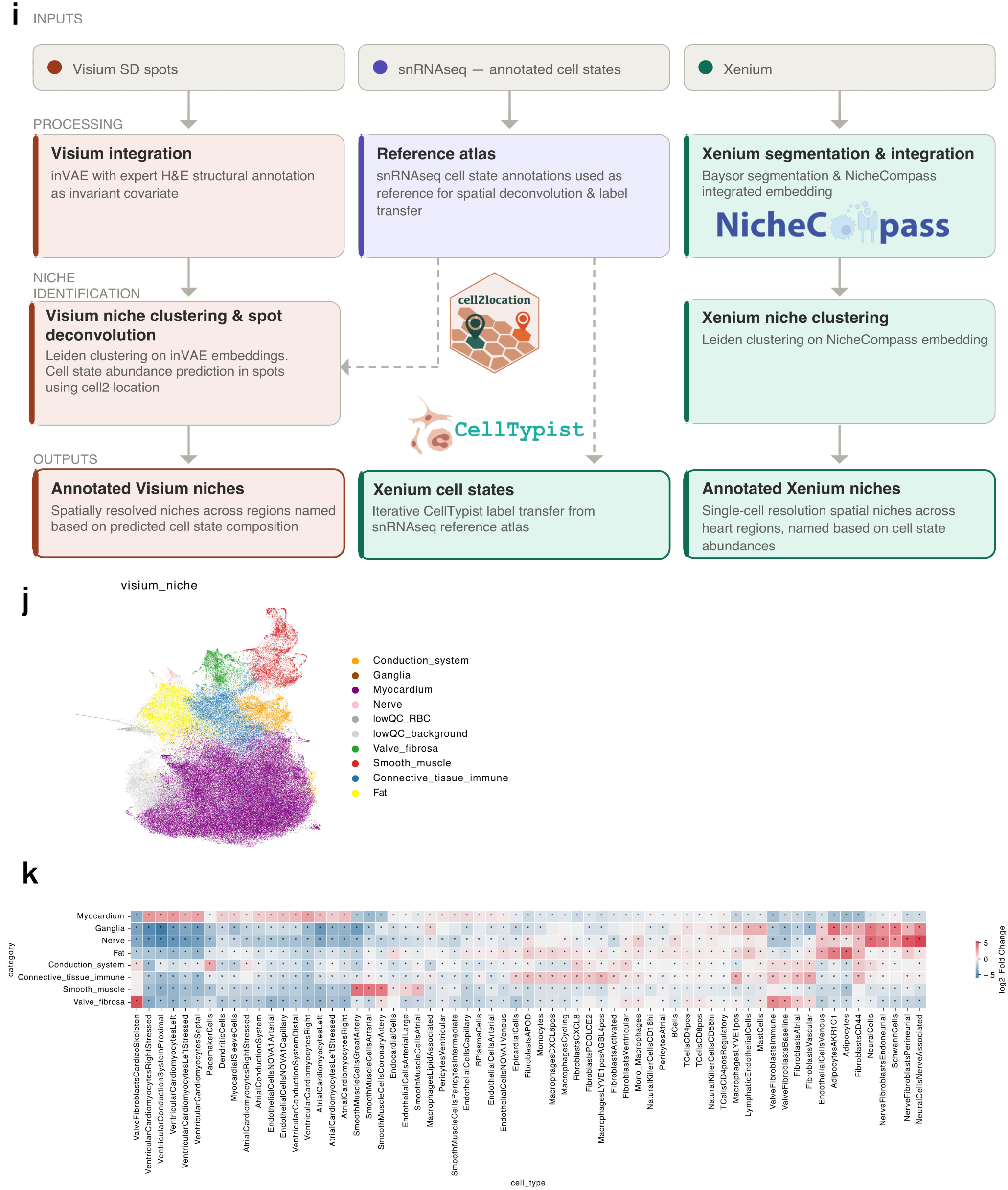
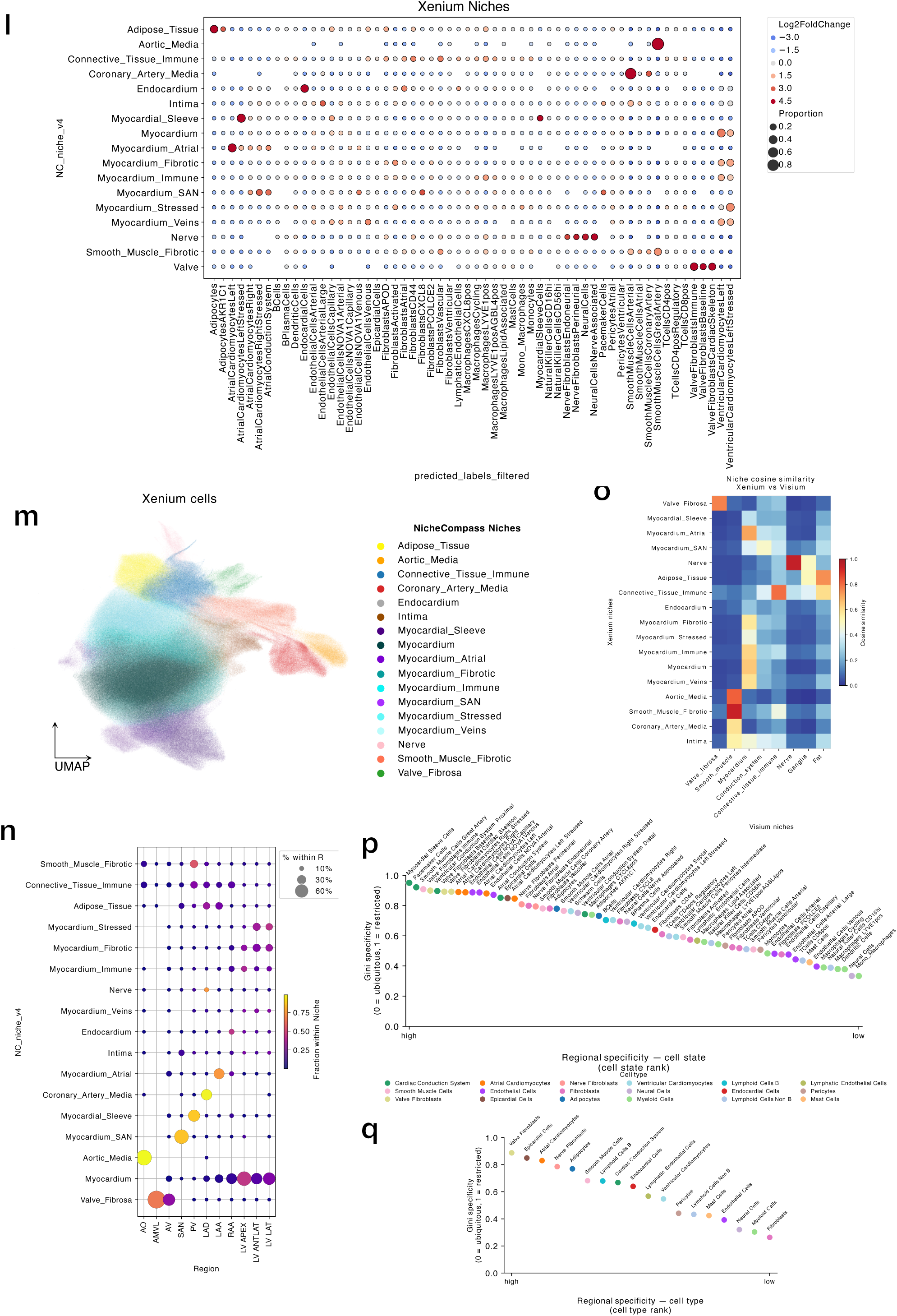
Cell-state annotation, taxonomy crosswalk and spatial niche enrichment. **a,** Number of donor-modality pairs per ‘region coarse’ (12 category system), stratified by donor age group. Number in white represents total, number in gold represents newly added in this study. **b,** Timeline showing age of donors contributing to this study. Each donor is a circle. Colour encodes modality: single-cell (RNA-seq and/or Multiome), spatial (Xenium and/or Visium), both. Outer gold rim reflects a new donor for this study (11 new in total). **c,** UMAP embedding of integrated single-cell and single-nucleus RNA-seq profiles in the inVAE latent space, coloured by annotated cell state. **d,** UMAP embedding of the same profiles in an scVI latent space, coloured by annotated cell state. **e,** Annotation robustness assessment showing 5-fold cross-validated *k*-nearest-neighbour classification accuracy (*k* = 15) for recovery of inVAE-derived cell-type labels in the scVI latent space, compared with the inVAE reference latent space. Points are global (using cell type categories) and cell-type-specific (using cell state categories) classification accuracies. **f,** Dot plot of marker gene expression across the 18 annotated cell types. Dot size indicates the fraction of cells in each cell type expressing the marker and colour denotes mean expression within that cell type. **g,** Cell-state taxonomy confusion matrix between Kanemaru *et al.* ^3^ and the updated annotation in this study. Colour denotes the fraction of cells assigned to each updated cell-state label that were assigned to each previous cell-state label. **h,** Cell-type taxonomy confusion matrix between Kanemaru *et al.* and the updated annotation. Colour denotes the fraction of cells assigned to each updated cell-type label that were assigned to each previous cell-type label. **i,** Schematic of the spatial annotation workflow. Visium standard-definition spots were integrated using inVAE with expert H&E structural annotation as an invariant covariate, deconvolved using cell2location with the single-nucleus RNA-seq reference, and clustered to define Visium niches. Xenium data were segmented with Baysor, cell states were transferred from the single-nucleus RNA-seq reference using CellTypist, and NicheCompass embeddings were clustered to define single-cell-resolution Xenium niches. **j,** UMAP embedding of Visium standard-definition spots from all profiled regions, coloured by assigned Visium niche. **k,** cell2location-predicted cell-state abundance enrichment across Visium niches. Colour indicates log_2_ fold change relative to all other Visium niches; asterisks indicate FDR-adjusted *P <* 0.05. **l,** Enrichment dot plot of transferred cell states within NicheCompass-derived Xenium niches. Dot size indicates the proportion of cells in each niche assigned to each cell state, and colour indicates log_2_ fold-change enrichment relative to all other Xenium niches. **m,** UMAP embedding of Xenium-derived cells coloured by NicheCompass niche identity, revealing 17 tissue niches. **n,** Regional distribution of NicheCompass-derived Xenium niches across anatomical sampling sites. Dot size denotes the proportion of segmented cells from each region assigned to each niche, and dot colour denotes the fraction of each niche located within each region. Region abbreviations: AO, aorta; AMVL, anterior mitral valve leaflet; AV, aortic valve; SAN, sinoatrial node; PV, pulmonary vein; LAD, left anterior descending coronary artery; LAA, left atrial appendage; RAA, right atrial appendage; LV APEX, left ventricular apex; LV ANTLAT, left ventricular anterolateral wall; LV LAT, left ventricular lateral wall. **o,** Comparison of Xenium (y axis) and Visium (x axis) niches using cosine similarity between cell state composition vectors derived from CellTypist (Xenium) and cell2location (Visium) using an identical snRNAseq training dataset. **p,** Regional specificity of cell states, quantified as a Gini index of mean cell-type proportion across anatomical regions (0, ubiquitous distribution across regions; 1, restriction to a single region). Cell states are ranked from most to least regionally restricted and coloured by their parent cell type. **q,** Regional specificity of cell types. Cell types are ranked from most to least regionally restricted and coloured by cell type.

**Figure S2: Supplementary Figure S2:**
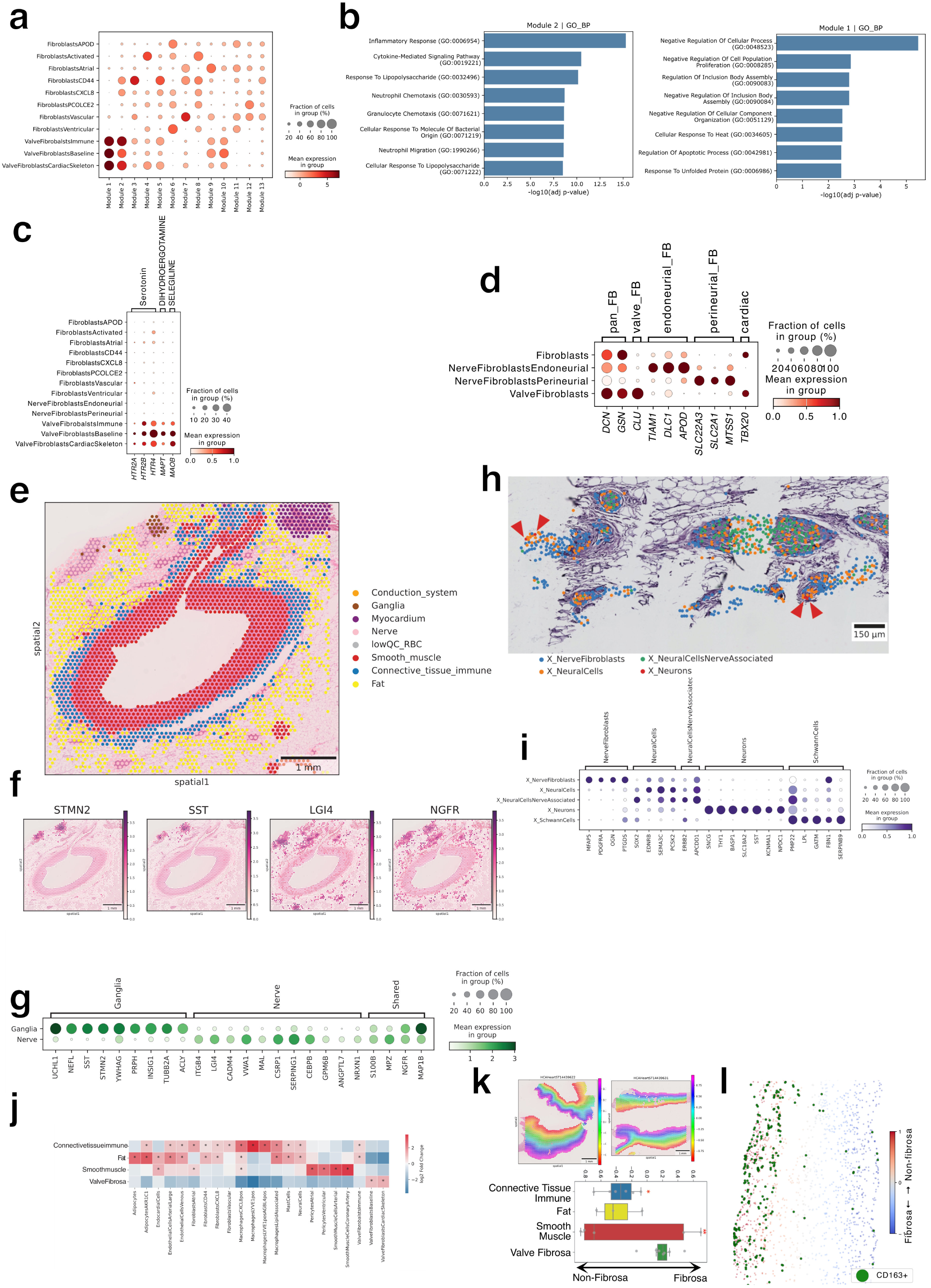
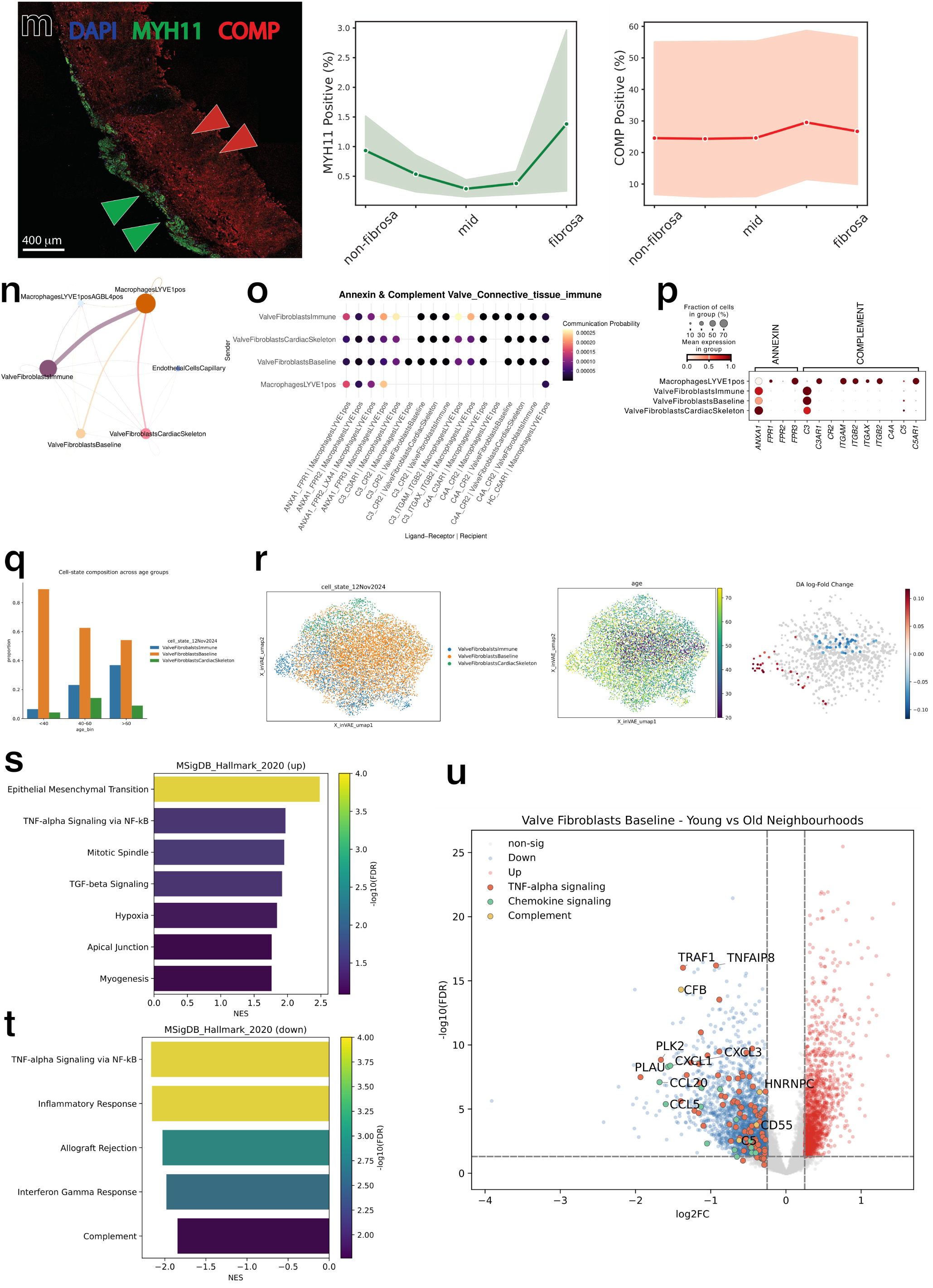
Fibroblast heterogeneity across cardiac niches. **a,** Dot plot showing the expression of Hotspot gene modules across fibroblast cell states. **b,** Gene Ontology enrichment analysis of genes assigned to modules 1 and 2 (enriched in Valve Fibroblasts). **c**, Drug2cell (d2c) scores dotplot of selected drugs implicated in serotonergic valvulopathy (dexfenfluramine, prucalopride, dihydroergotamine) and the MAO-B inhibitor selegiline, grouped by mechanism: serotonergic agonists (5-HT2/5-HT4), MAPT-binding ergot, and MAO-B. **d,** Dotplot of marker genes comparing Nerve Fibroblasts cell states (Endoneurial and Perineurial) with other fibroblast cell types. Cardiac development transcription factor *TBX20* expression is prominently absent from Nerve Fibroblasts. **e**, Spatial projection of Visium niches in a coronary artery section highlighting Ganglia and Nerves as distinct niches. **f**, Spatial plot of niche marker genes of Ganglia (*STMN2* and *SST*), Nerves (*LGI4*) and a shared marker of these niches (*NGFR*). **g**, Dotplot of niche marker genes of Nerves and Ganglia in Visium data. **h,** Spatial projection of Xenium neural tissue associated cell states including Neurons (red arrows) overlaid on post-Xenium H&E stain. **i,** Marker genes dotplot for the neural tissue associated cell states in Xenium. **j,** Predicted cell state abundance (cell2location) enrichment in Visium niches of the cardiac valves. Colour denotes log2 Fold Change compared to all other niches, * denotes FDR adjusted p < 0.05 **k,** (Top) Visium sections with OrganAxis-derived coordinates representing relative distance from the flow-exposed surface (negative values; mitral valve atrialis / aortic valve ventricularis) to the pressure-exposed surface (positive values; fibrosa), enabling quantitative comparison of niche composition across the valve depth. (Bottom) Boxplots summarising the mean OrganAxis position of each niche in each sample, with * indicating statistically significant positional bias (observed vs expected). **l**, Spatial distribution of CD163+ macrophages (green dots) across a valve section in post-Xenium Rarecyte staining, with cells coloured by position along the fibrosa-to-non-fibrosa axis. **m**, Immunofluorescence imaging of aortic (n=3) and mitral (n=3) valve leaflets localising the smooth muscle marker (MYH11) to the surface of the valve and the valve fibroblast marker (COMP) throughout the leaflets. **n,** Circle plot depicting CellChat-inferred signalling probabilities between cell states within the connective tissue immune niche in valves for all pathways combined. **o**, Dot plot of communication probabilities for individual ligand-receptor pairs within the Annexin and Complement pathways in connective tissue immune Xenium niche of cardiac valves, showing sender cell states (rows) and ligand-receptor-recipient combinations (columns). **p**, Dot plot confirming expression of Annexin and Complement pathway ligands and receptors in snRNAseq data. Dot size indicates the fraction of cells expressing each gene and colour reflects scaled mean expression level. **q,** Proportional changes in valvular fibroblast states across ageing periods. **r,** UMAP of aortic-valve Valve Fibroblasts showing the distribution of cell states, age and ageing-associated differential-abundance log fold change. **s,** Gene set enrichment results for genes upregulated when old age-enriched aortic Valve Fibroblasts Immune neighbourhoods were compared with other Valve Fibroblasts Immune neighbourhoods. The x axis represents the normalised enrichment score (NES), and colour indicates − log_10_(FDR). **t,** Gene set enrichment results for genes downregulated when old age-enriched aortic Valve Fibroblasts Baseline neighbourhoods were compared with other Valve Fibroblasts Baseline neighbourhoods. Corresponding to pathways enriched in old age-enriched genes. The x axis represents the normalised enrichment score (NES), and colour indicates − log_10_(FDR). **u,** Volcano plot of differentially expressed genes in younger-enriched versus remaining Baseline Valve Fibroblast neighbourhoods. Highlighted genes are coloured by pathway: TNF*α* signalling (yellow), Chemokine signalling (green).

**Figure S3: Supplementary Figure S3:**
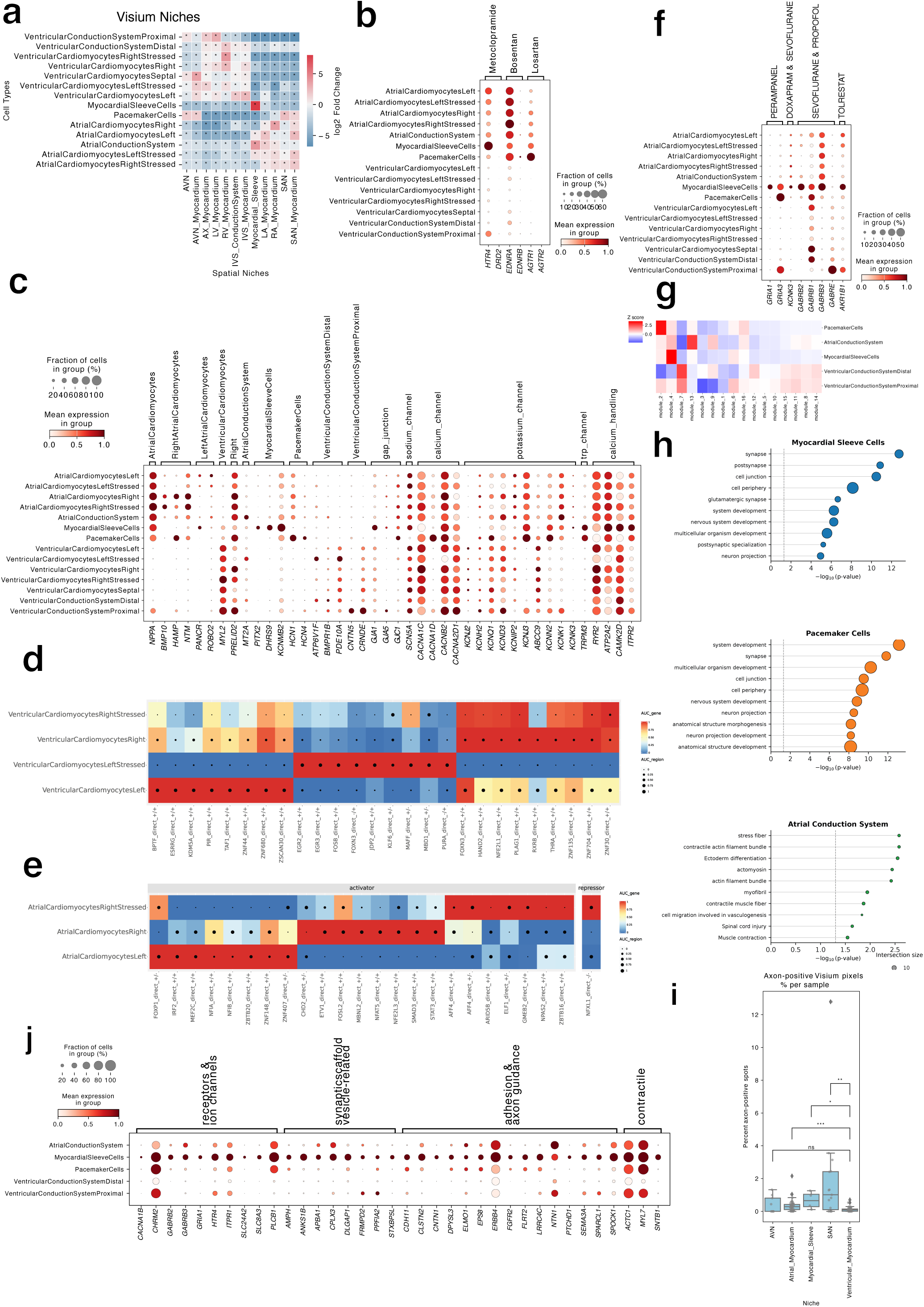
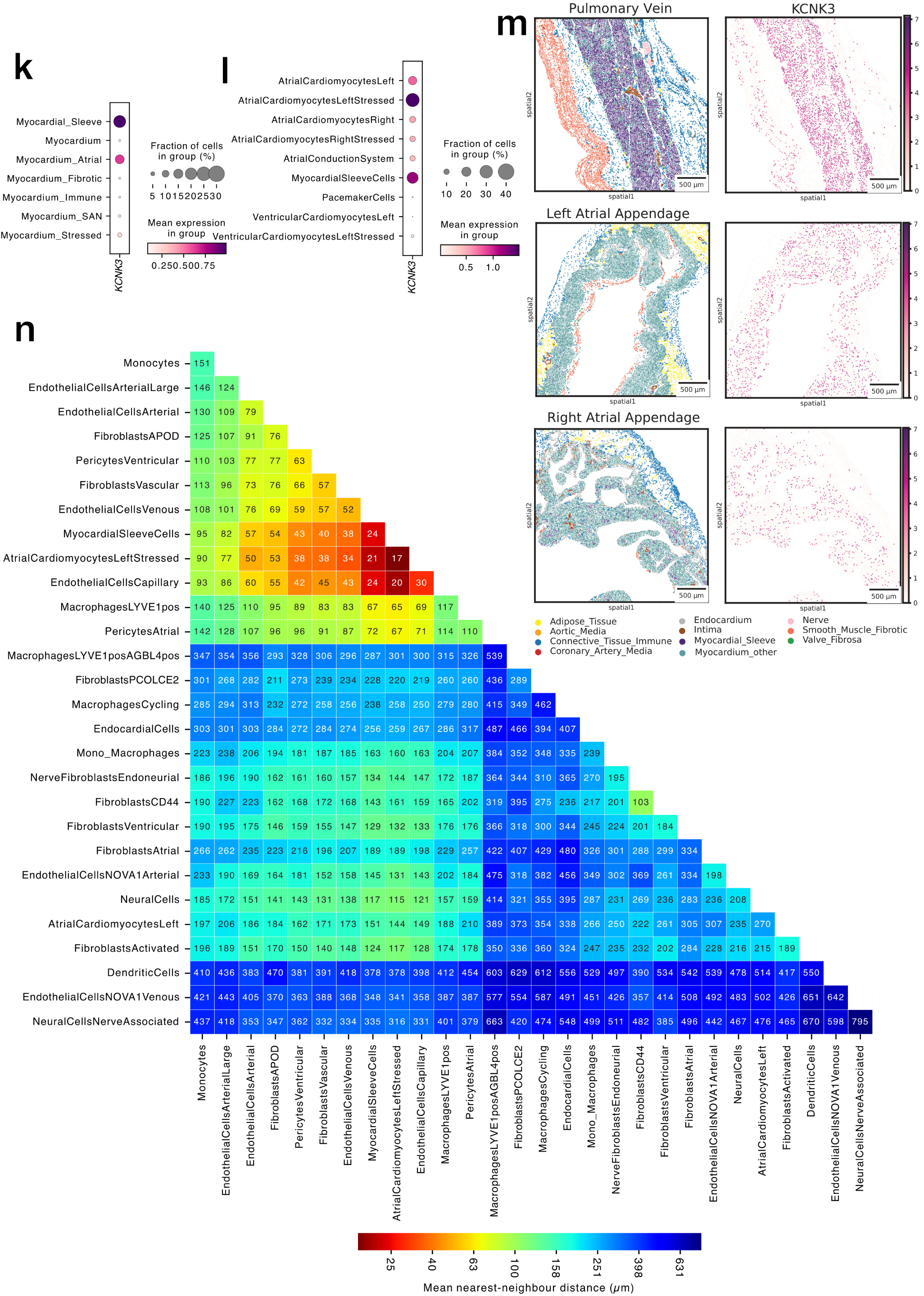
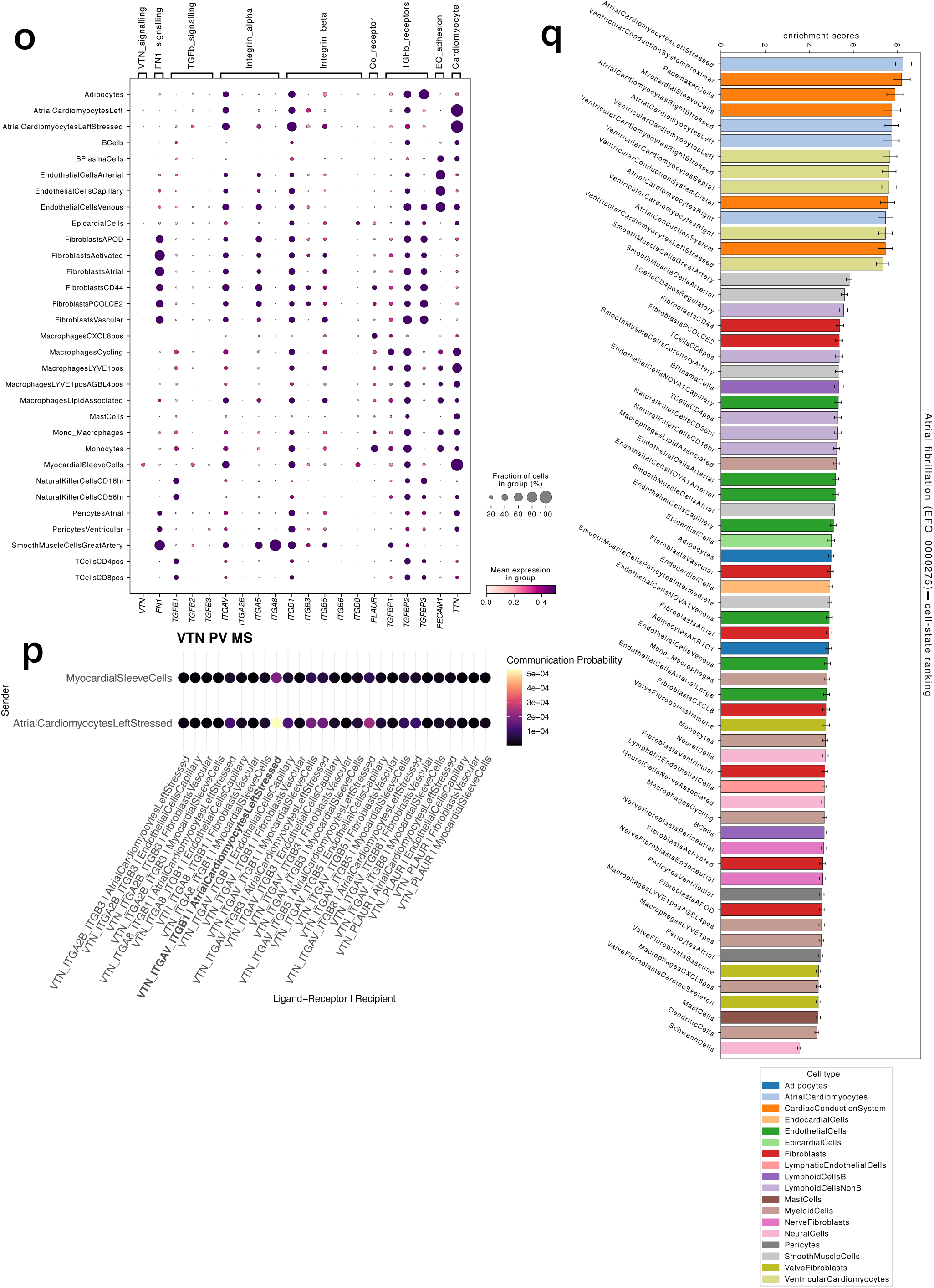
Cardiomyocytes and their niches. **a,** cell2location-predicted cell-state abundance enrichment across conduction system and other myocardial Visium niches divided by region. Colour indicates log_2_ fold change relative to all other niches; asterisks indicate FDR-adjusted *P <* 0.05. **b,** drug2cell analysis showing drug targets with predicted enrichment in atrial, ventricular and conduction system cardiomyocyte cell states. **c,** Dot plot depicting expression of key marker genes, ion channels and cardiac conduction implicated genes in atrial, ventricular and conduction system cardiomyocyte cell states. **d** and **e,** Heatmap showing SCENIC+ enhancer-driven regulon activity in ventricular (d) and atrial (e) cardiomyocyte metacells. Box colour indicates mean direct gene-based AUCell and dot size indicates mean direct region-based AUCell, summarised by cardiomyocyte cell state after excluding states with fewer than three pure-cell state metacells. Regulons were filtered to retain only eRegulons passing expression, gene–region AUCell consistency, regulon-specificity and donor-robustness criteria. The top quality-passing regulons per state are shown, ranked by Mann–Whitney U testing and biological relevance. **f,** drug2cell analysis showing drug targets with predicted enrichment in atrial, ventricular and conduction system cardiomyocyte cell states, including perampanel, doxapram, sevoflurane, propofol and tolrestat. **g,** Hotspot analysis of cardiac conduction system cells highlighting gene expression modules specific to Myocardial Sleeve Cells (module 4), Pacemaker Cells (module 2) and Atrial Conduction System (module 13). **h,** Gene-ontology terms enriched in cell state-specific Hotspot modules. **i,** Percentage of Visium standard-definition spots expected to contain axons (i.e. expressing *PRPH*, *NEFL*, *NEFM* or *NEFH*), stratified by Visium myocardial niches. (∗ *P <* 0.05, ∗ ∗ *P <* 0.01, ∗ ∗ ∗ *P <* 0.001; Mann–Whitney, FDR-adjusted). **j,** Dotplot of selected genes from the Myocardial Sleeve Cells-enriched Hotspot module (module 4), grouped by biological pathways. **k,** Expression of *KCNK3* in myocardial Xenium niches. This encodes the two-pore-domain potassium channel TASK-1, identified as a drug target in Myocardial Sleeve Cells by drug2cell analysis. **l,** *KCNK3* expression in cardiomyocyte and conduction system Xenium cells. **m,** Spatial projections showing Xenium niche annotations (left) and *KCNK3* expression (right) in the pulmonary vein, left atrial appendage (LAA) and right atrial appendage (RAA). Myocardial niches other than Myocardial Sleeve were merged under one label for visual clarity. **n,** Mean nearest-neighbour distance between cell state pairs in the Myocardial Sleeve Xenium niche of the pulmonary veins, combining data from three pulmonary vein sections. **o,** Expression of ligands and receptors in single-nucleus RNA-seq cell states of pulmonary veins. **p,** Dot plot of communication probabilities for individual ligand–receptor pairs within the vitronectin pathway in the Myocardial Sleeve Xenium niche of the pulmonary veins, showing sender cell states (rows) and ligand–receptor–recipient combinations (columns). **q,** SNP2CELL network-propagation scores summarised across all 65 heart cell states for AF (EFO_0000275). For each node in the regulatory network, SNP2CELL integrates a GWAS-propagation z-score with a cell-state DE-specificity z-score using the element-wise minimum of the two robust median/MAD-based z-scores. Bars show the mean ± 95% CI of the integrated score across GWAS-prioritised nodes with z-MAD > 2. Bar and tick colours indicate parent cell-type.

**Figure S4: Supplementary Figure S4:**
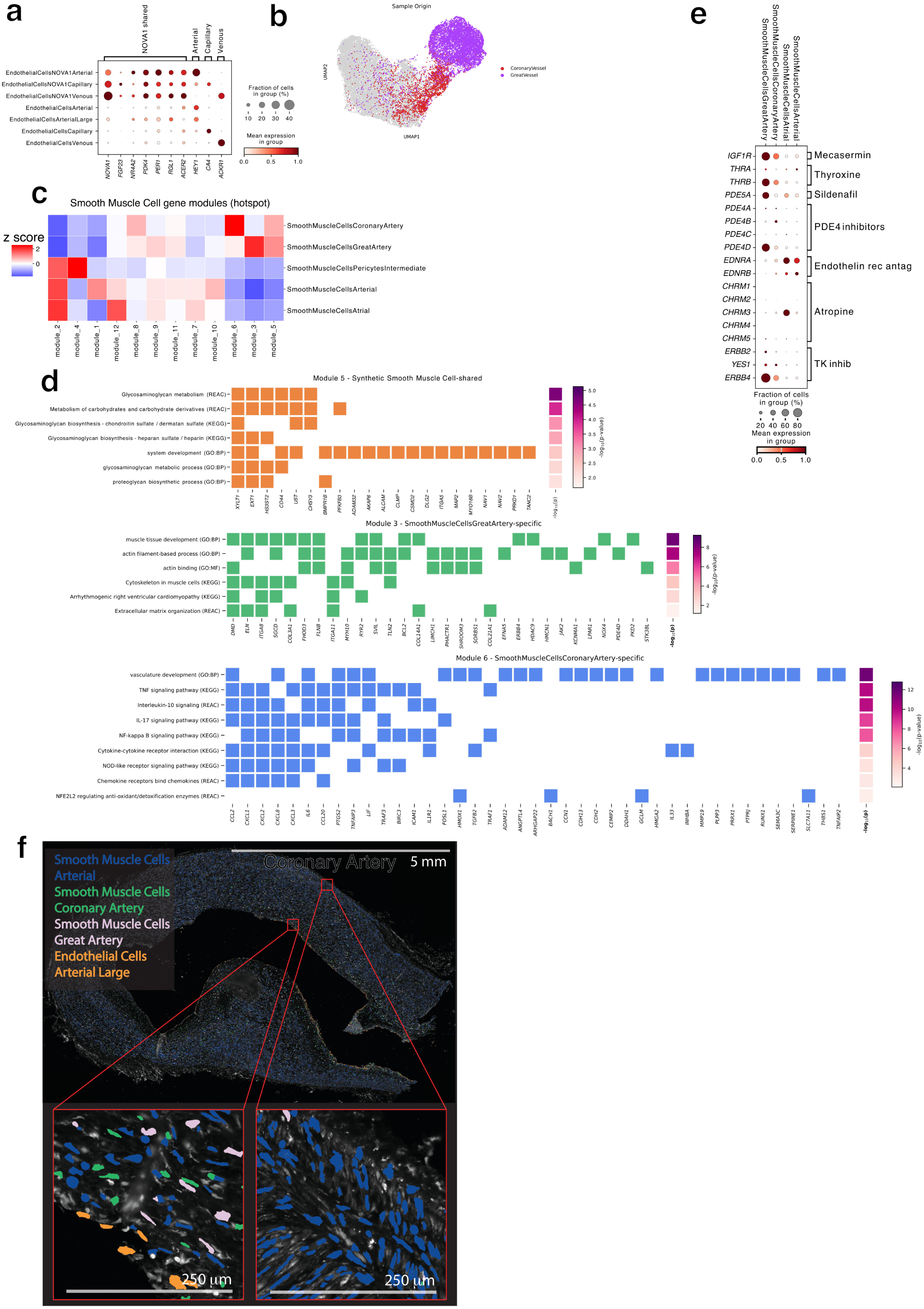
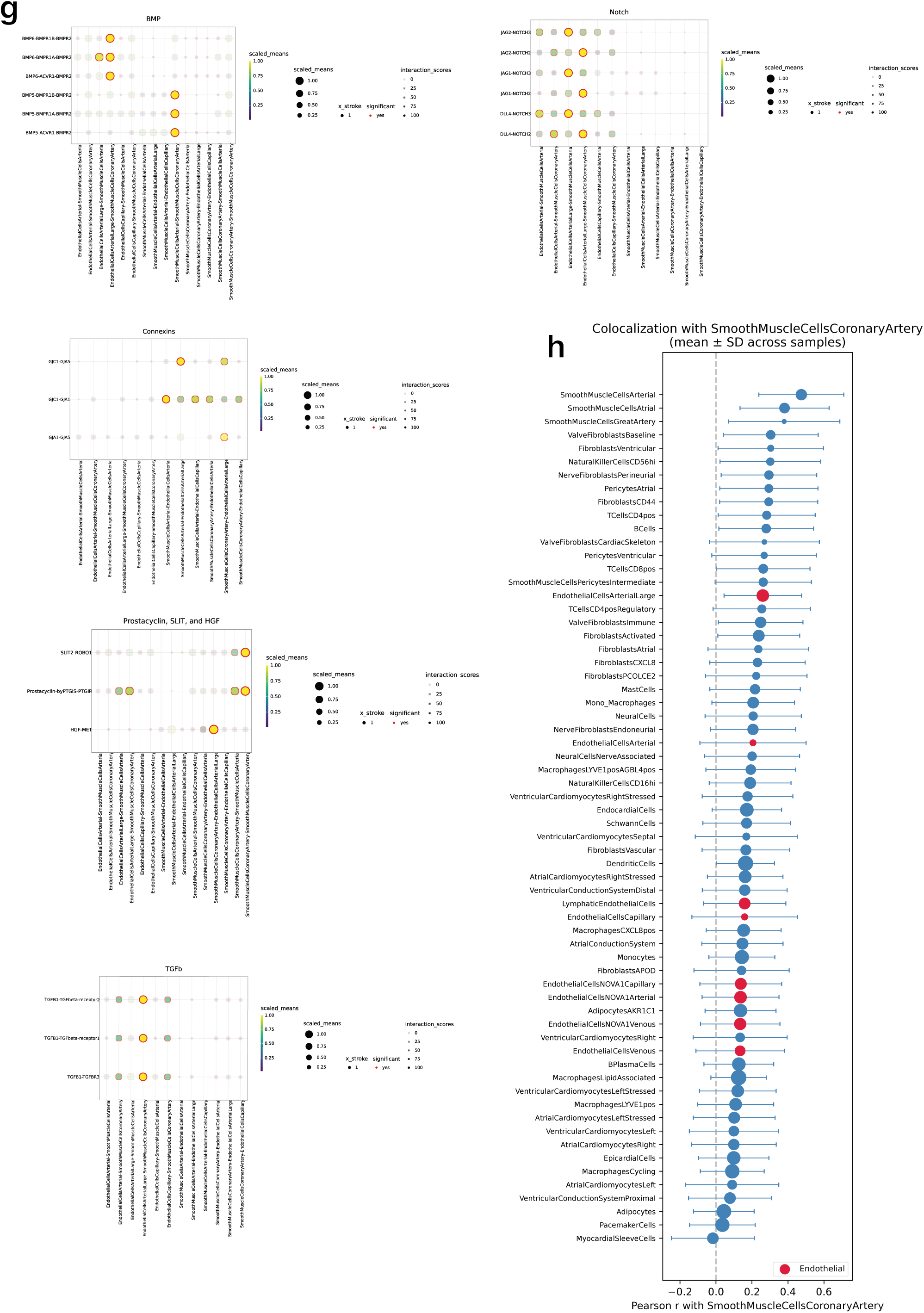
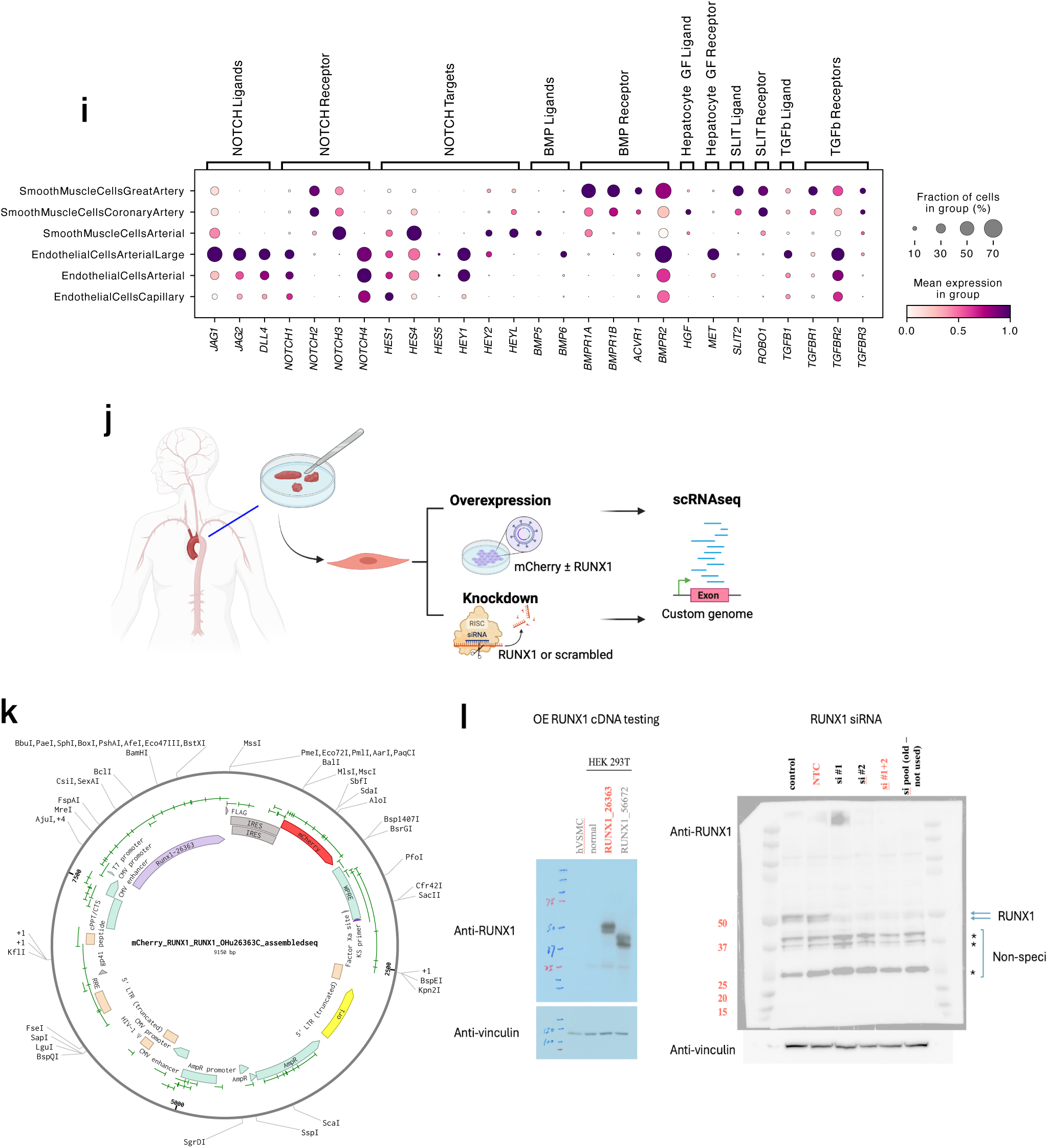
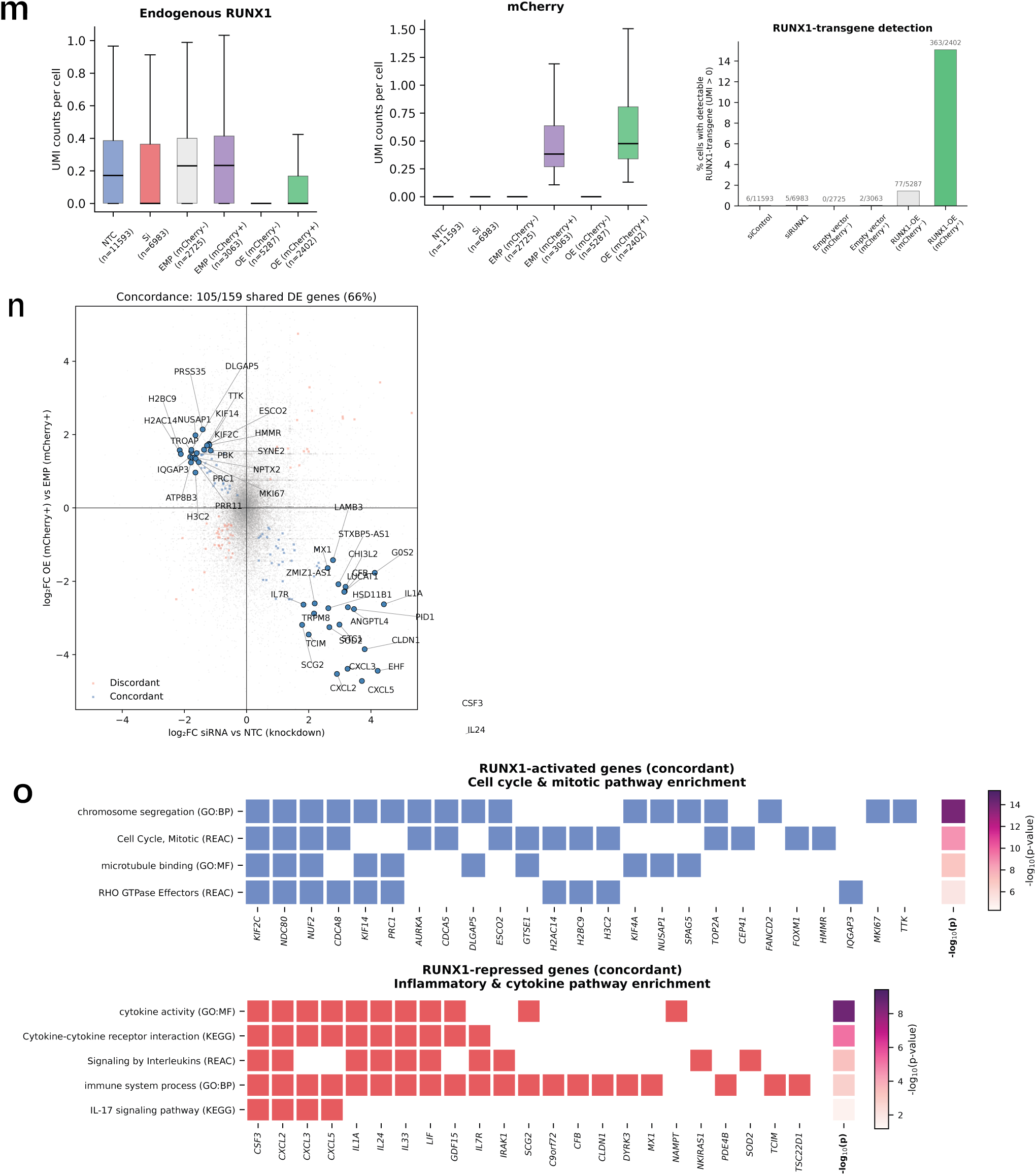
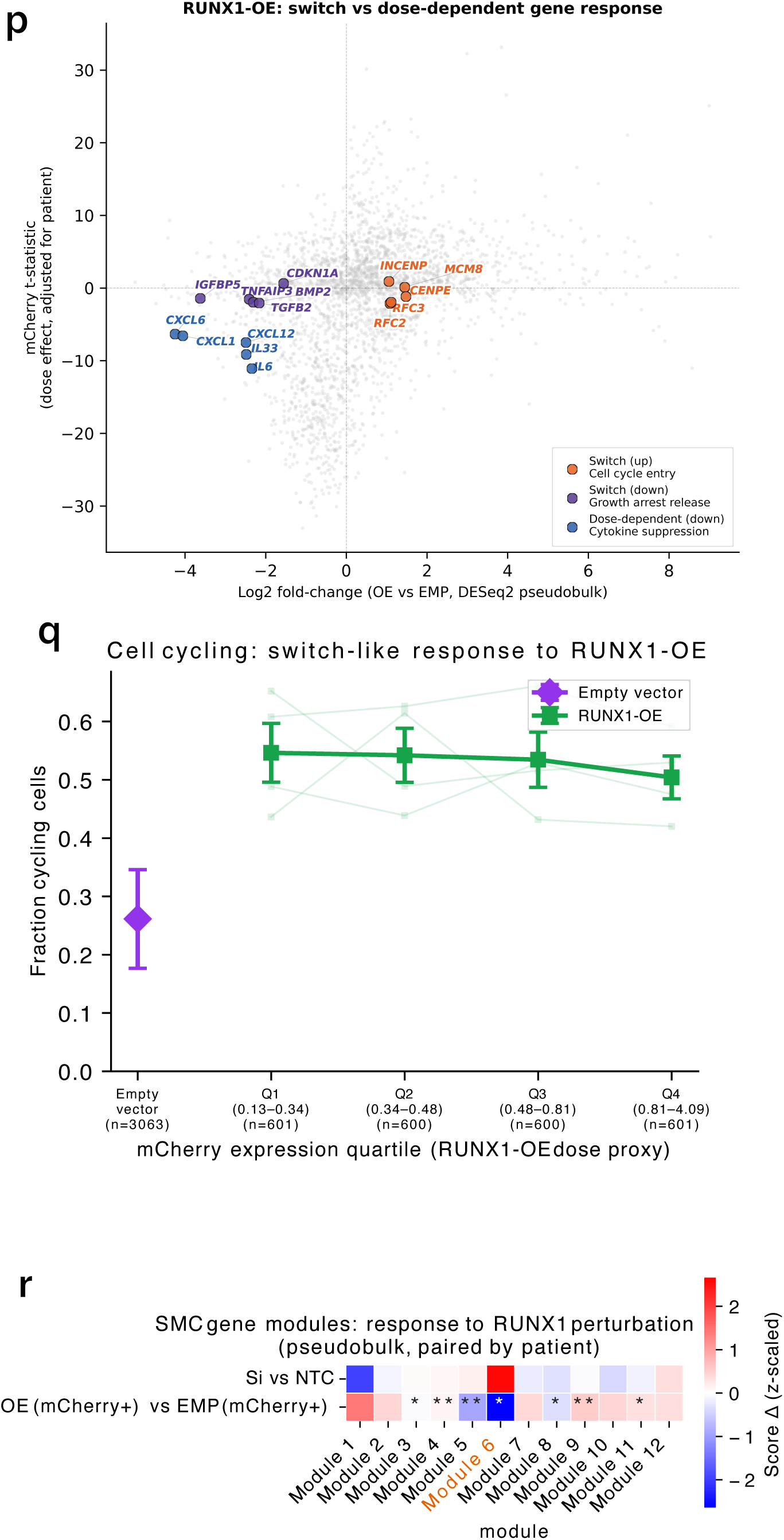
Vascular smooth muscle cell specialisation, coronary artery niche signalling and RUNX1 perturbation. **a,** Dot plot of the shared *NOVA1* endothelial programme (*NOVA1*, *FGF23*, *NR4A2*, *PDK4*, *PER1*, *RGL1*, *ACER2*) together with vascular-bed identity markers (arterial, *HEY1*; capillary, *CA4*; venous, *ACKR1*) across endothelial cell states. Dot size denotes the fraction of cells expressing each gene and colour denotes mean expression, scaled per gene. **b,** UMAP embedding of smooth muscle cells (all cell states) coloured by anatomical sampling source, highlighting coronary-vessel-derived and great-vessel-derived cells. **c,** Heatmap of Hotspot-derived gene module expression across annotated smooth muscle cell states. Colour denotes the *z*-scored module activity in each state (red, high; blue, low; white, zero-centred). **d,** Marker genes and pathway enrichment terms for selected smooth muscle cell gene modules: Module 5, shared by synthetic smooth muscle cells (Smooth Muscle Cells Coronary Artery and Great Artery); Module 3, enriched in Smooth Muscle Cells Great Artery; and Module 6, enriched in Smooth Muscle Cells Coronary Artery. Coloured squares indicate genes contributing to each enriched pathway and side bars denote enrichment significance as − log_10_(*P*). **e,** drug2cell dot plot of drug-target gene expression across smooth muscle cell states. Dot size denotes the fraction of cells expressing each target gene and colour denotes mean expression within the cell state. **f,** Xenium spatial transcriptomics (5,001-gene panel) of a coronary artery cross-section, with segmented cells coloured by cell state; insets show the subintima and tunica media. **g,** CellPhoneDB ligand–receptor analysis between smooth muscle and endothelial cell states for BMP, NOTCH, connexin, prostacyclin, SLIT–ROBO, HGF–MET and TGF*β* signalling interactions. Plot aesthetics encode scaled mean expression, interaction score and CellPhoneDB significance. **h,** Spatial colocalisation of cell states with Smooth Muscle Cells Coronary Artery in Visium standard-definition data from all anatomical regions. Points show the mean Pearson correlation coefficient across samples and horizontal bars show standard deviation; endothelial cell states are highlighted. **i,** Dot plot showing expression of ligand, receptor and downstream target genes for selected signalling pathways (NOTCH, BMP, HGF–MET, SLIT–ROBO and TGF*β*) across smooth muscle and endothelial cell states. Dot size denotes the fraction of cells expressing each gene and colour denotes mean expression within the cell state. **j,** Experimental design for bidirectional *RUNX1* perturbation in primary human smooth muscle cells. Primary human aortic smooth muscle cells were subjected to lentiviral *RUNX1* overexpression or siRNA-mediated *RUNX1* knockdown, followed by single-cell RNA-seq using a custom reference including the transgene sequence. **k,** Lentiviral plasmid map for the mCherry–*RUNX1* overexpression construct. **l,** Western blot validation of the perturbation reagents. Left, testing of *RUNX1* cDNA overexpression constructs in HEK293T; right, validation of siRNA-mediated *RUNX1* knockdown. Vinculin is shown as a loading control. **m,** Single-cell UMI count distributions for endogenous *RUNX1* and mCherry across perturbation groups, together with the percentage of cells with detectable *RUNX1* transgene counts. Cell numbers are shown below group labels and positive-cell counts above bars. Abbreviations: NTC, non-targeting control (the siRNA control arm); siRNA, *RUNX1* siRNA; EMP (mCherry^+^), empty-vector transduced cells; OE (mCherry^+^), *RUNX1*-overexpression transduced cells. **n,** Concordance of pseudobulk differential expression responses to reciprocal *RUNX1* perturbation. Each point is a gene. The x-axis shows log_2_ fold-change for *RUNX1* siRNA versus non-targeting control and the y-axis log_2_ fold-change for *RUNX1* overexpression versus empty vector in mCherry^+^ cells. Genes regulated in opposite directions by knockdown and overexpression are classified as concordant (105/159 shared differentially expressed genes, 66%). Highlighted genes include cell cycle and inflammatory cytokine genes. **o,** Pathway enrichment of concordantly *RUNX1*-activated and *RUNX1*-repressed genes. Activated genes are enriched for cell-cycle and mitotic pathways; repressed genes for inflammatory and cytokine-signalling pathways. Coloured squares indicate genes contributing to each pathway and side bars denote enrichment significance as — log_10_(*P*). **p,** Classification of *RUNX1*-overexpression responses into switch-like and dose-dependent components. Each point is a gene. The x-axis shows pseudobulk log_2_ fold-change for *RUNX1* overexpression versus empty vector (DESeq2); the y-axis shows the mCherry dose effect, the *t*-statistic of the mCherry coefficient from a per-gene ordinary least-squares regression of single-cell expression on mCherry level with donor as a covariate, fitted in overexpression mCherry^+^ cells. Genes that are differentially expressed but dose-independent are classified as switch-like, and those with a significant mCherry coefficient as dose-dependent. **q,** Fraction of cycling cells across mCherry expression quartiles in *RUNX1*-overexpressing cells, using mCherry as a proxy for overexpression dose; empty-vector mCherry^+^ cells shown for comparison. The proliferative response is approximately constant across dose quartiles. **r,** Response of the *in vivo* Hotspot-derived smooth muscle cell gene modules to *RUNX1* perturbation by donor-paired pseudobulk analysis. Colour denotes the *z*-scaled change in module score for *RUNX1* siRNA versus non-targeting control and for *RUNX1* overexpression versus empty vector in mCherry^+^ cells; asterisks indicate significant changes. Module 6, the Smooth Muscle Cells Coronary Artery-specific module, is highlighted. Abbreviations: NTC, non-targeting control; Si, *RUNX1* siRNA; EMP, empty vector; OE, *RUNX1* overexpression.

**Figure S5: Supplementary Figure S5:**
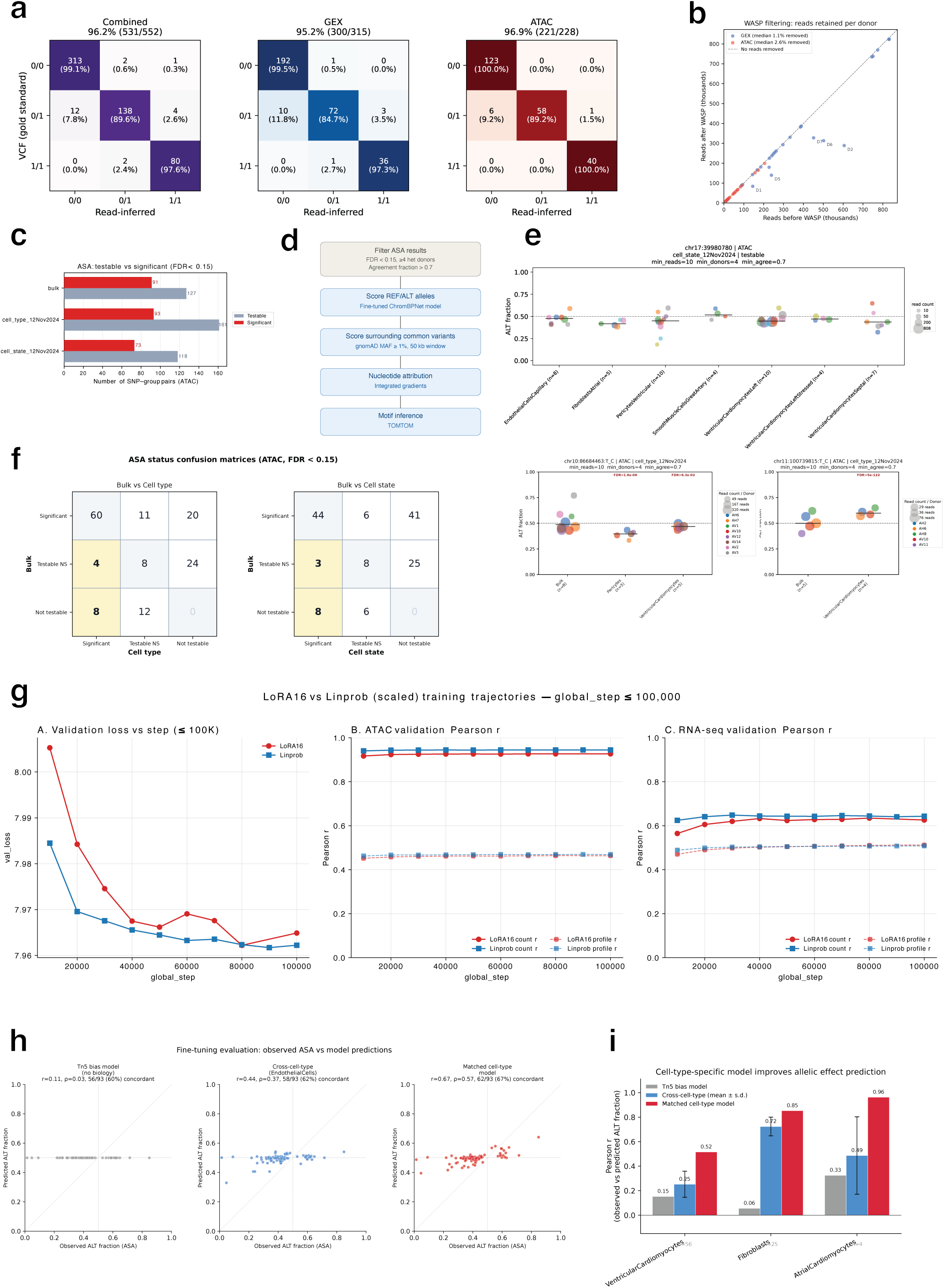
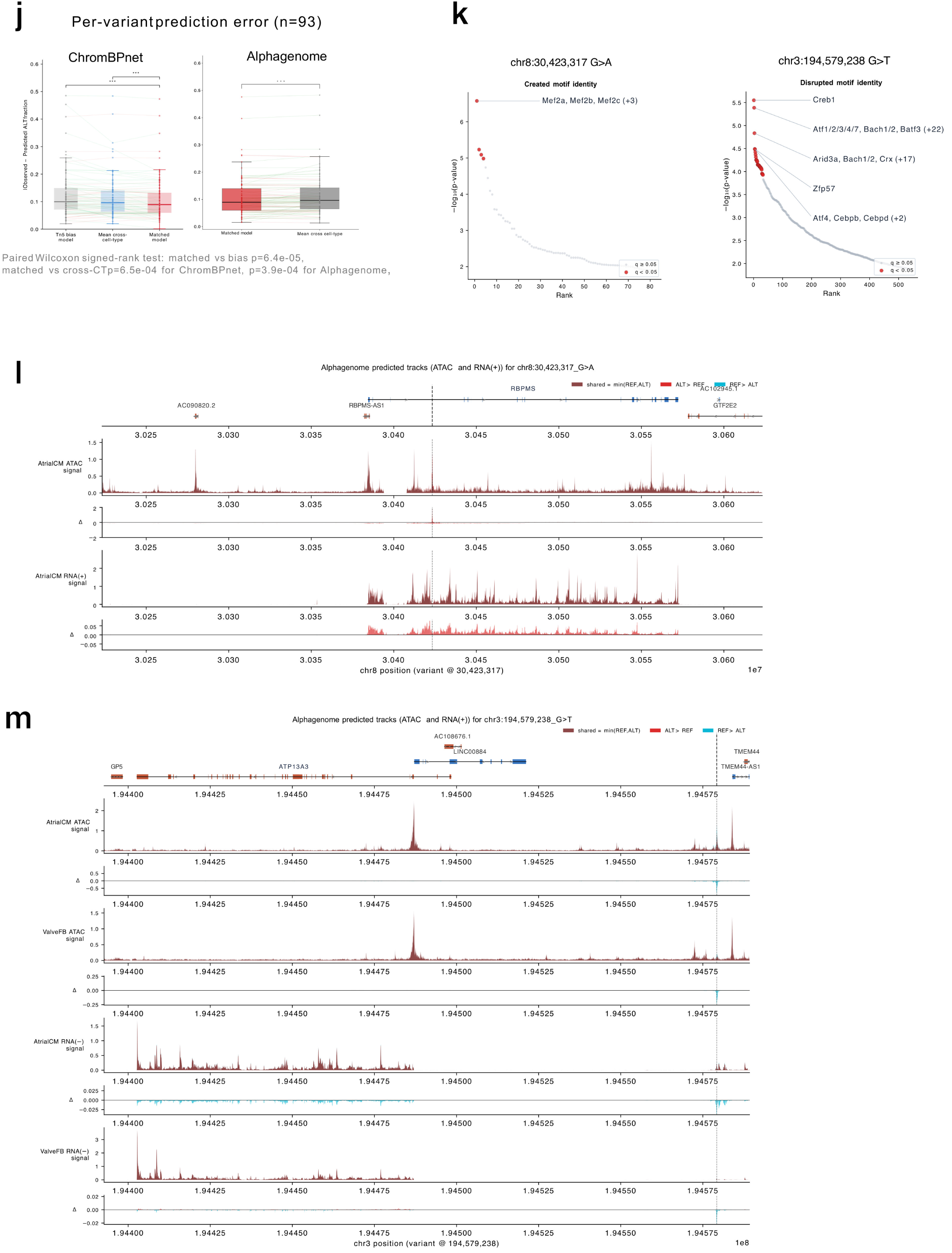
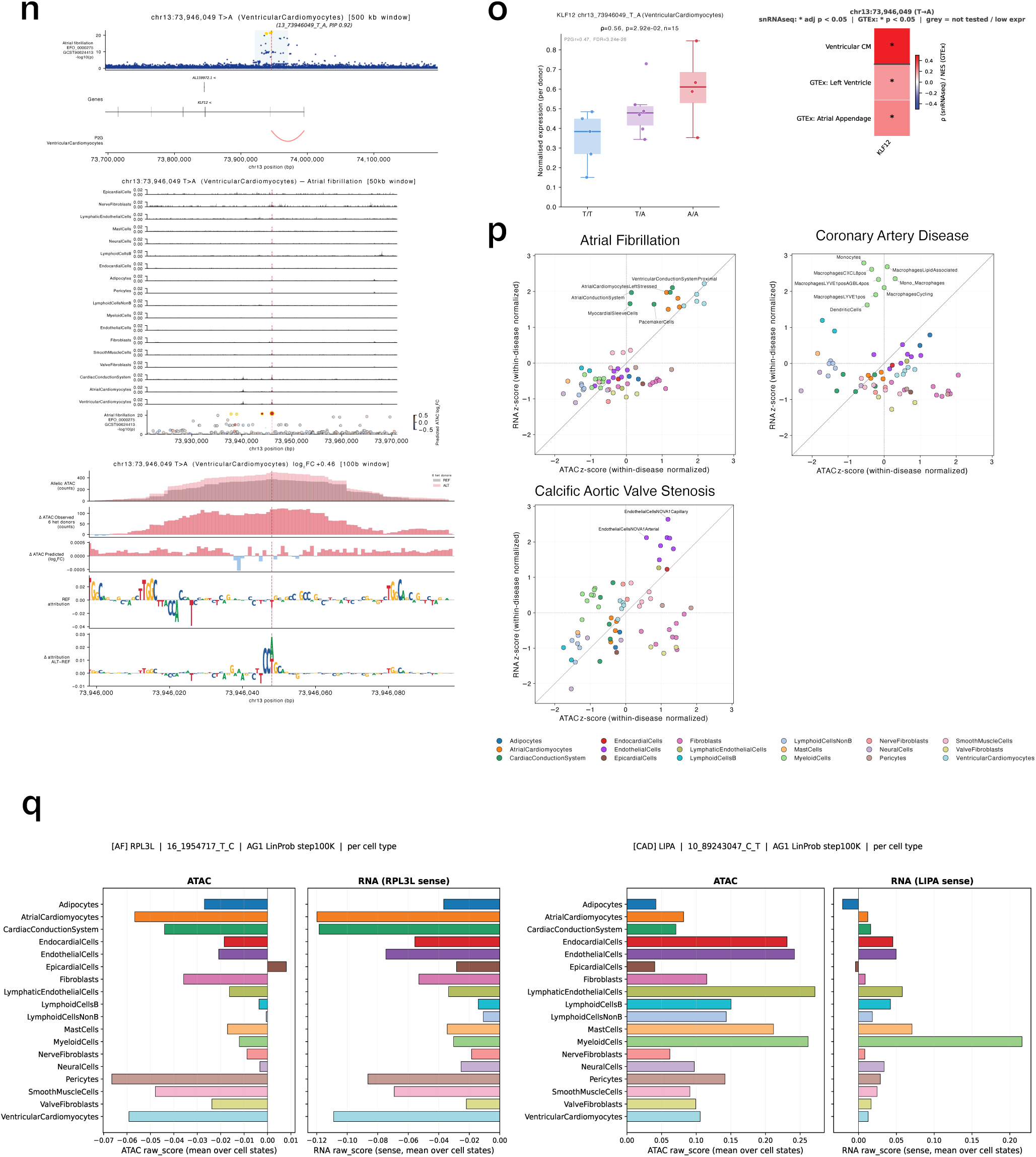
Allele-specific accessibility and sequence-model prediction of cell type-specific GWAS variant effects. **a,** Concordance of read-inferred donor genotypes with array-derived VCF genotypes. Confusion matrices are shown for combined GEX and ATAC reads, GEX reads only and ATAC reads only for each donor-variant pair. **b,** Effect of WASP allelic remapping on retained read counts per donor. Points show GEX and ATAC read counts before and after WASP filtering; the diagonal indicates no reads removed. Off-diagonal samples (D1, D2, D5, D6, D7) were mapped using a different hg38 reference, however these donors (first published in Litvinukova *et al.* ^1^) did not undergo Multiome and therefore did not contribute to ASA analysis. **c,** Number of SNP–group pairs (where a group is a donor at bulk, donor-cell type or donor-cell state level) passing ASA testing filters and reaching significance at the bulk, cell-type and cell-state levels. Significant pairs were defined at FDR *<* 0.15. **d,** Workflow for interpreting ASA-positive variants with fine-tuned ChromBPNet. ASA-significant variants were filtered by FDR, heterozygous donor count and allelic agreement, REF and ALT alleles were scored with the matched cell-type model, neighbouring common variants were scored within a 50-kb window, nucleotide attributions were calculated using integrated gradients and motif identities were inferred with TOMTOM. **e,** Representative allele-specific accessibility examples. Points denote heterozygous donors, point size denotes UMI-deduplicated read count and colours denote donors. The dashed horizontal line indicates balanced allelic accessibility at ALT fraction 0.5; FDR values are shown for significant group-level tests. **f,** Comparison of ASA status between bulk and cell-type-level testing, and between bulk and cell-state-level testing. Entries show the number of SNP–group pairs classified as significant, testable but non-significant, or not testable at FDR *<* 0.15. At the cell-type level, 12 SNP–group pairs were significant that were not significant at the bulk level (8 not testable, 4 testable but not significant). **g,** AlphaGenome fine-tuning benchmark comparing LoRA16 and linear-probe models over training steps. Panels show validation loss, ATAC validation Pearson correlation and RNA-seq validation Pearson correlation for count and profile prediction tasks. **h,** Comparison of observed ASA with ChromBPNet-predicted ALT allele fraction for a Tn5-bias-only baseline, a cross-cell-type model and the matched cell-type model. Pearson correlation, Spearman correlation and directional concordance are indicated above each panel. **i,** Cell-type-specific performance of ChromBPNet allelic effect prediction. Bars show Pearson correlation between observed and predicted ALT fraction for Ventricular Cardiomyocytes, Fibroblasts and Atrial Cardiomyocytes using the Tn5-bias baseline, cross-cell-type models and matched cell-type models. **j,** Per-variant prediction error for ChromBPNet and AlphaGenome. Left, absolute error between observed and predicted ALT fraction across ASA-significant SNP–cell-type pairs for the Tn5-bias baseline, mean cross-cell-type prediction and matched cell-type model for ChromBPNet. Right, paired comparison of prediction error for matched versus mean cross-cell-type models for AlphaGenome predictions. Significance was assessed by paired Wilcoxon signed-rank test (∗ ∗ ∗ *P <* 0.001). **k,** TOMTOM-ranked motif identities inferred from ChromBPNet attribution differences. Left, attribution differences for the atrial fibrillation-associated variant rs2979489 (chr8:30,423,317 G>A) are consistent with creation of a MEF2-family motif. Right, analysis of the aortic valve stenosis-associated variant rs1706003 (chr3:194,579,238 G>T) suggests disruption of a CREB1-like motif. **l,** AlphaGenome-predicted ATAC and positive-strand RNA tracks are shown for Atrial Cardiomyocytes at the atrial fibrillation-associated variant rs2979489 (chr8:30,423,317 G>A). The genomic window is centred on the variant, indicated by the vertical dashed line. For each modality, the upper panel shows the predicted signal, whereas the lower panel shows the change in predicted signal from the reference to the alternative allele (REF→ALT Δ signal). Shared signal is shown as the minimum of REF and ALT predictions, with ALT > REF and REF > ALT components highlighted separately. **m,** As in **l**, AlphaGenome predictions for the aortic valve stenosis-associated variant rs1706003 (chr3:194,579,238 G>T) in Atrial Cardiomyocytes and Valve Fibroblasts; the vertical dashed line indicates the variant position. **n,** Multi-scale locus view for the atrial fibrillation-associated variant rs1886512 (chr13:73,946,049 T*>*A) in Ventricular Cardiomyocytes, as in Fig. 5f. **Top** (500 kb window, SNP marked by the red dashed line): GWAS association (− log_10_ *p*) for atrial fibrillation, with credible-set variants in gold and all others in blue; a gene track; and cell type-specific ArchR peak-to-gene (P2G) arcs coloured by peak–gene correlation (red positive, blue negative). **Middle** (50 kb window): cell type-specific ATAC accessibility tracks, with a lower scatter of common variants coloured by their AlphaGenome-predicted accessibility effect (red increase, blue decrease; gold edge marks credible-set membership; grey points lack a prediction). **Bottom** (100 bp window): allelic ATAC pileup over donors heterozygous for rs1886512, with the observed and predicted ALT−REF differences and base-resolution contribution scores for the REF allele followed by the ALT−REF difference. **o,** Genotype-expression evidence for *KLF12* at rs1886512. Left, donor-level normalised *KLF12* expression in Ventricular Cardiomyocytes stratified by genotype. Right, comparison of *KLF12* genotype-expression effects in snRNA-seq and GTEx heart tissues. Colour denotes Spearman *ρ* for snRNA-seq or GTEx normalised effect size; asterisks indicate significant associations and grey denotes genes not tested or with low expression. **p,** Disease-level summary of AlphaGenome-predicted variant effects across credible-set SNPs for atrial fibrillation, coronary artery disease and aortic valve stenosis. Each point denotes a cell state coloured by parent cell type. The x-axis shows within-disease-normalised ATAC effect score and the y-axis shows within-disease-normalised RNA effect score. **q,** AlphaGenome-predicted per-cell-type variant effects at exemplar disease loci. Predicted ATAC and positive-strand RNA raw scores are shown for *RPL3L* at an atrial fibrillation locus and *LIPA* at a coronary artery disease locus, averaged over cell states within each parent cell type. Abbreviations: ASA, allele-specific accessibility; REF, reference allele; ALT, alternative allele; GEX, gene expression; ATAC, assay for transposase-accessible chromatin; VCF, variant call format; P2G, peak-to-gene.

## References

[1] Litvinukova, M. et al. Cells of the adult human heart. Nature (2020).

[2] Tucker, N. R. et al. Transcriptional and cellular diversity of the human heart. Circulation (2020).

[3] Kanemaru, K. et al. Spatially resolved multiomics of human cardiac niches. Nature (2023).

[4] Aragam, K. G. et al. Discovery and systematic characterization of risk variants and genes for coronary artery disease in over a million participants. Nature Genetics (2022).

[5] Roselli, C. et al. Multi-ethnic genome-wide association study for atrial fibrillation. Nature Genetics (2018).

[6] Aliee, H., et al. invae: conditionally invariant representation learning for generating multivariate single-cell reference maps. bioRxiv (2024).

[7] Gayoso, A., Lopez, R., Xing, G. et al. A Python library for probabilistic analysis of single-cell omics data. Nature Biotechnology 40, 163–166 (2022).

[8] Kleshchevnikov, V. et al. Cell2location maps fine-grained cell types in spatial transcriptomics. Nature Biotechnology (2022).

[9] Petukhov, V. et al. Cell segmentation in imaging-based spatial transcriptomics. Nature Biotechnology (2022).

[10] Domínguez Conde, C., et al. Cross-tissue immune cell analysis reveals tissue-specific features in humans. Science (2022).

[11] Birk, S. et al. Quantitative characterization of cell niches in spatially resolved omics data. Nature Genetics (2025).

[12] Humphreys, D. T. et al. Clusterin has chaperone-like activity similar to that of small heat shock proteins. Journal of Biological Chemistry (1999).

[13] Hochgrebe, T. T. et al. A reexamination of the role of clusterin as a complement regulator. Experimental Cell Research (1999).

[14] Rothman, R. B. et al. Evidence for possible involvement of 5-HT2B receptors in the cardiac valvulopathy associated with fenfluramine and other serotonergic medications. Circulation (2000).

[15] Sosdean, R. et al. Monoamine oxidase contributes to valvular oxidative stress: A prospective observational pilot study in patients with severe mitral regurgitation. International Journal of Molecular Sciences 25, 10307 (2024).

[16] Madissoon, E. et al. A spatially resolved atlas of the human lung characterizes a gland-associated immune niche. Nature Genetics 55, 66–77 (2023).

[17] Latif, N., Sarathchandra, P., Chester, A. H. & Yacoub, M. H. Expression of smooth muscle cell markers and co-activators in calcified aortic valves. European Heart Journal 36, 1335–1345 (2015).

[18] Zheng, Y. et al. Glioma-derived ANXA1 suppresses the immune response to TLR3 ligands by promoting an anti-inflammatory tumor microenvironment. Cellular Molecular Immunology 21, 47–59 (2024).

[19] To, K. et al. A multi-omic atlas of human embryonic skeletal development. Nature (2024).

[20] Wang, S. et al. PALMD regulates aortic valve calcification via altered glycolysis and NF-*κ*B-mediated inflammation. Journal of Biological Chemistry 298, 101887 (2022).

[21] Taniguchi, K. & Karin, M. NF-*κ*B, inflammation, immunity and cancer: coming of age. Nature Reviews Immunology 18, 309–324 (2018).

[22] Wang, J. et al. Pitx2 prevents susceptibility to atrial arrhythmias by inhibiting left-sided pacemaker specification. Proceedings of the National Academy of Sciences (2010).

[23] Reyat, J. S., et al. Reduced left atrial cardiomyocyte PITX2 and elevated circulating BMP10 predict atrial fibrillation after ablation. JCI Insight 5, e139179 (2020).

[24] Hennings, E. et al. Association of bone morphogenetic protein 10 and recurrent atrial fibrillation after catheter ablation. Europace 25, euad149 (2023).

[25] Gore-Panter, S. R. et al. PANCR, the PITX2 adjacent noncoding RNA, is expressed in human left atria and regulates PITX2c expression. Circulation: Arrhythmia and Electrophysiology 9, e003197 (2016).

[26] Sheikh, F. et al. An fhl1-containing complex within the cardiomyocyte sarcomere mediates hypertrophic biomechanical stress responses in mice. Journal of Clinical Investigation 118, 3870–3880 (2008).

[27] Liang, Y.-J. et al. Mechanical stress enhances serotonin 2b receptor modulating brain natriuretic peptide through nuclear factor-*κ*b in cardiomyocytes. Cardiovascular Research 72, 303–312 (2006).

[28] Yoshida, Y. et al. Ccn1 protects cardiac myocytes from oxidative stress via *β*1 integrin–akt pathway. Biochemical and Biophysical Research Communications 355, 611–618 (2007).

[29] McCalmon, S. A. et al. Modulation of angiotensin II–mediated cardiac remodeling by the MEF2A target gene Xirp2. Circulation Research 106, 952–960 (2010).

[30] Haissaguerre, M. et al. Spontaneous initiation of atrial fibrillation by ectopic beats originating in the pulmonary veins. New England Journal of Medicine (1998).

[31] Gudbjartsson, D. F. et al. Variants conferring risk of atrial fibrillation on chromosome 4q25. Nature (2007).

[32] Napoli, J. L. Physiological insights into all-trans-retinoic acid biosynthesis. Biochimica et Biophysica Acta (BBA)-Molecular and Cell Biology of Lipids 1821, 152–167 (2012).

[33] Deshmukh, A. et al. Left atrial transcriptional changes associated with atrial fibrillation susceptibility and persistence. Circulation: Arrhythmia and Electrophysiology 8, 32–41 (2015).

[34] Moldenhauer, H. J., Dinsdale, R. L., Alvarez, S., Fernández-Jaén, A. & Meredith, A. L. Effect of an autism-associated kcnmb2 variant, g124r, on bk channel properties. Current Research in Physiology 5, 404–413 (2022).

[35] Li, F. & Yuan, Y. Meta-analysis of the cardioprotective effect of sevoflurane versus propofol during cardiac surgery. BMC Anesthesiology 15, 128 (2015).

[36] Wiedmann, F. et al. Treatment of atrial fibrillation with doxapram: TASK-1 potassium channel inhibition as a novel pharmacological strategy. Cardiovascular Research 118, 1728–1741 (2022).

[37] Schmidt, C. et al. Upregulation of k2p3.1 k+ current causes action potential shortening in patients with chronic atrial fibrillation. Circulation 132, 82–92 (2015).

[38] Schmidt, C. et al. Inverse remodelling of k2p3.1 k+ channel expression and action potential duration in left ventricular dysfunction and atrial fibrillation: Implications for patient-specific antiarrhythmic drug therapy. European Heart Journal 38, 1764–1774 (2017).

[39] Perez-Pomares, J. M. et al. Retinoic acid signalling during cardiac development. Development (2004).

[40] Barnett, S. N. et al. An organotypic atlas of human vascular cells. Nature Medicine (2024).

[41] Gong, L., Wang, S., Shen, L., et al. SLIT3 deficiency attenuates pressure overload–induced cardiac fibrosis and remodeling. JCI Insight 5, e136852 (2020).

[42] Zandstra, T. E. et al. Asymmetry and heterogeneity: part and parcel in cardiac autonomic innervation and function. Frontiers in Physiology (2021).

43. Allahverdian, S., et al. Transcriptomic, specific marker, and pathway analysis of smooth muscle cell foam cells compared to macrophage foam cells in human atherosclerosis. bioRxiv (2026).

[44] Okuda, T. et al. AML1, the target of multiple chromosomal translocations in human leukemia, is essential for normal fetal liver hematopoiesis. Cell (1996).

[45] Lambert, J. et al. Network-based prioritization and validation of regulators of vascular smooth muscle cell proliferation. Nature Cardiovascular Research (2024).

[46] Stary, H. C. et al. A definition of the intima of human arteries and of its atherosclerosis-prone regions: a report from the committee on vascular lesions of the council on arteriosclerosis, american heart association. Circulation (1992).

[47] Nakashima, Y. et al. Early atherosclerosis in humans: role of diffuse intimal thickening and extracellular matrix proteoglycans. Cardiovascular Research (2008).

[48] Bleckwehl, T. et al. Encompassing view of spatial and single-cell rna sequencing renews the role of the microvasculature in human atherosclerosis. Nature Cardiovascular Research 4, 26–44 (2025).

[49] Efremova, M. et al. CellPhoneDB: inferring cell-cell communication from combined expression of multi-subunit ligand-receptor complexes. Nature Protocols (2020).

[50] Chen, Y., et al. Circulating hepatocyte growth factor reflects activation of vascular repair in response to stress. JACC: Basic to Translational Science (2022).

[51] Ogunmoroti, O. et al. Hepatocyte growth factor is associated with greater risk of extracoronary calcification: results from the multi-ethnic study of atherosclerosis. Open Heart (2022).

[52] Lane, K. B. et al. Heterozygous germline mutations in BMPR2, encoding a TGF-*β* receptor, cause familial primary pulmonary hypertension. Nature Genetics (2000).

[53] Tirosh, I. et al. Dissecting the multicellular ecosystem of metastatic melanoma by single-cell RNA-seq. Science 352, 189–196 (2016).

[54] van de Geijn, B. et al. WASP: allele-specific software for robust molecular quantitative trait locus discovery. Nature Methods (2015).

[55] Kim-Hellmuth, S. et al. Cell type-specific genetic regulation of gene expression across human tissues. Science (2020).

[56] Trevino, A. E. et al. Chromatin and gene-regulatory dynamics of the developing human cerebral cortex at single-cell resolution. Cell (2021).

[57] Avsec, Z. et al. Alphagenome: advancing regulatory variant effect prediction with a unified dna sequence model. Nature (2026).

[58] Gan, P. et al. Rbpms regulates cardiomyocyte contraction and cardiac function through rna alternative splicing. Cardiovascular Research (2024).

[59] Yuan, S. et al. Cross-population GWAS and proteomics improve risk prediction and reveal mechanisms in atrial fibrillation. Nature Communications (2025).

[60] Mayr, B. & Montminy, M. Transcriptional regulation by the phosphorylation-dependent factor CREB. Nature Reviews Molecular Cell Biology 2, 599–609 (2001).

[61] Southgate, L. et al. Atp13a3 in pulmonary arterial hypertension and polyamine transport. Cardiovascular Research (2024).

[62] Hu, M. et al. KLF12 aggravates angiotensin II-induced cardiac remodeling in male mice by transcriptionally inhibiting SMAD7. Journal of the American Heart Association (2025).

[63] Li, F. et al. LIPA, a risk locus for coronary artery disease: decoding the variant-to-function relationship. European Heart Journal 46, 5273–5288 (2025).

[64] Shiraishi, C. et al. RPL3L-containing ribosomes determine translation elongation dynamics required for cardiac function. Nature Communications 14, 2131 (2023).

[65] Minvielle Moncla, L.-H., Briend, M., Bossé, Y. & Mathieu, P. Calcific aortic valve disease: mechanisms, prevention and treatment. Nature Reviews Cardiology 20, 546–559 (2023).

[66] Passos, L. S. A., Lupieri, A., Becker-Greene, D. & Aikawa, E. Innate and adaptive immunity in cardiovascular calcification. Atherosclerosis 306, 59–67 (2020).

[67] Qin, C. X. et al. Cardioprotective actions of the annexin-A1 N-terminal peptide, Ac2-26, against myocardial infarction. Frontiers in Pharmacology 10, 269 (2019).

[68] Prevete, N., Poto, R., Marone, G. & Varricchi, G. Unleashing the power of formyl peptide receptor 2 in cardiovascular disease. Cytokine 169, 156298 (2023).

[69] Van Gelder, I. C. et al. 2024 ESC guidelines for the management of atrial fibrillation developed in collaboration with the european association for cardio-thoracic surgery (EACTS). European Heart Journal (2024).

[70] Mommersteeg, M. T. M. et al. Pitx2c and nkx2-5 are required for the formation and identity of the pulmonary myocardium. Circulation Research (2007).

[71] Kirchhof, P. et al. Pitx2c is expressed in the adult left atrium and reducing pitx2c expression promotes atrial fibrillation inducibility. Circulation: Cardiovascular Genetics (2011).

[72] Wirka, R. C. et al. Atheroprotective roles of smooth muscle cell phenotypic modulation and the tcf21 disease gene. Nature Medicine (2019).

[73] Li, D. Y., Monteiro, J. P., Zhao, Q., Stitziel, N. O. & Quertermous, T. Single-cell studies advance understanding of the genetic and molecular basis of atherosclerosis. Circulation Research 139, e327472 (2026).

[74] Amrute, J. M. et al. Targeting modulated vascular smooth muscle cells in atherosclerosis via FAP-directed im-munotherapy. Science (2026).

[75] Stary, H. C. et al. Natural history and histological classification of atherosclerotic lesions. Arteriosclerosis, Thrombosis, and Vascular Biology (2000).

[76] Nakazawa, G. et al. Pathological intimal thickening in human coronary arteries. *Arteriosclerosis*, Thrombosis, and Vascular Biology (2007).

[77] Martin, T. P. et al. Ribonucleic acid interference or small molecule inhibition of runx1 in the border zone prevents cardiac contractile dysfunction following myocardial infarction. Cardiovascular Research (2023).

[78] Hingerl, J. C. et al. scooby: modeling multimodal genomic profiles from DNA sequence at single-cell resolution. Nature Methods (2025).

[79] Hamouda, N. N. et al. ATP13A3 is a major component of the enigmatic mammalian polyamine transport system. Journal of Biological Chemistry (2020).

[80] Regev, A. et al. The human cell atlas. eLife (2017).

[81] Bogovic, J. A., Hanslovsky, P., Wong, A. & Saalfeld, S. Robust registration of calcium images by learned contrast synthesis. 2016 IEEE 13th International Symposium on Biomedical Imaging (ISBI) 1123–1126 (2016).

[82] Schindelin, J., Arganda-Carreras, I., Frise, E. et al. Fiji: an open-source platform for biological-image analysis. Nature Methods 9, 676–682 (2012).

[83] van der Walt, S., Schönberger, J. L., Nunez-Iglesias, J. et al. scikit-image: image processing in Python. PeerJ 2, e453 (2014).

[84] Heaton, H. et al. Souporcell: robust clustering of single-cell RNA-seq data by genotype without reference genotypes. Nature Methods 17, 615–620 (2020).

[85] Auton, A., Brooks, L. D., Durbin, R. M. et al. A global reference for human genetic variation. Nature 526, 68–74 (2015).

[86] Fleming, S. J., Chaffin, M. D., Arduini, A. et al. Unsupervised removal of systematic background noise from droplet-based single-cell experiments using CellBender. Nature Methods 20, 1323–1335 (2023).

[87] Wolf, F. A., Angerer, P. & Theis, F. J. SCANPY: large-scale single-cell gene expression data analysis. Genome Biology 19, 15 (2018).

[88] Wolock, S. L., Lopez, R. & Klein, A. M. Scrublet: Computational identification of cell doublets in single-cell transcriptomic data. Cell Systems 8, 281–291.e9 (2019).

[89] Oliver, A. J. et al. Single-cell integration reveals metaplasia in inflammatory gut diseases. Nature 635, 699–707 (2024).

[90] Granja, J. M., Corces, M. R., Pierce, S. E. et al. ArchR is a scalable software package for integrative single-cell chromatin accessibility analysis. Nature Genetics 53, 403–411 (2021).

[91] Lopez, R., Regier, J., Cole, M. B., Jordan, M. I. & Yosef, N. Deep generative modeling for single-cell transcriptomics. Nature Methods 15, 1053–1058 (2018).

[92] Pedregosa, F., Varoquaux, G., Gramfort, A. et al. Scikit-learn: Machine learning in Python. Journal of Machine Learning Research 12, 2825–2830 (2011).

[93] DeTomaso, D. & Yosef, N. Hotspot identifies informative gene modules across modalities of single-cell genomics. Cell Systems 12, 446–456.e9 (2021).

[94] Raudvere, U., Kolberg, L., Kuzmin, I. et al. g:Profiler: a web server for functional enrichment analysis and conversions of gene lists (2019 update). Nucleic Acids Research 47, W191–W198 (2019).

[95] Dann, E., Henderson, N. C., Teichmann, S. A., Morgan, M. D. & Marioni, J. C. Differential abundance testing on single-cell data using k-nearest neighbor graphs. Nature Biotechnology 40, 245–253 (2022).

[96] Heumos, L. et al. Pertpy: an end-to-end framework for perturbation analysis. Nature Methods 23, 350–359 (2026).

[97] Robinson, M. D., McCarthy, D. J. & Smyth, G. K. edgeR: a Bioconductor package for differential expression analysis of digital gene expression data. Bioinformatics 26, 139–140 (2010).

[98] Fang, Z., Liu, X. & Peltz, G. GSEApy: a comprehensive package for performing gene set enrichment analysis in Python. Bioinformatics 39, btac757 (2023).

[99] Schep, A. N., Wu, B., Buenrostro, J. D. & Greenleaf, W. J. chromVAR: inferring transcription-factor-associated accessibility from single-cell epigenomic data. Nature Methods 14, 975–978 (2017).

[100] Weirauch, M. T., Yang, A., Albu, M. et al. Determination and inference of eukaryotic transcription factor sequence specificity. Cell 158, 1431–1443 (2014).

[101] Rauluseviciute, I., Riudavets-Puig, R., Blanc-Mathieu, R. et al. JASPAR 2024: 20th anniversary of the open-access database of transcription factor binding profiles. Nucleic Acids Research 52, D174–D182 (2024).

[102] Yayon, N. et al. A spatial human thymus cell atlas mapped to a continuous tissue axis. Nature (2024).

[103] Polanski, K., Bartolomé-Casado, R., Sarropoulos, I., et al. Bin2cell reconstructs cells from high-resolution Visium HD data. Bioinformatics 40, btae546 (2024).

[104] Schmidt, U., Weigert, M., Broaddus, C. & Myers, G. Cell detection with star-convex polygons. Medical Image Computing and Computer Assisted Intervention (MICCAI) 265–273 (2018).

[105] Mages, S., Moriel, N., Avraham-Davidi, I. et al. TACCO unifies annotation transfer and decomposition of cell identities for single-cell and spatial omics. Nature Biotechnology 41, 1465–1473 (2023).

[106] Bergen, V. et al. Generalizing rna velocity to transient cell states through dynamical modeling. Nature Biotechnology (2020).

[107] Xu, C. et al. Automatic cell-type harmonization and integration across human cell atlas datasets. Cell (2023).

[108] Miranda, A.-M. A. et al. Selective remodelling of the adipose niche in obesity and weight loss. Nature (2025).

[109] Jin, S. et al. Inference and analysis of cell-cell communication using cellchat. Nature Communications (2021).

[110] Stringer, C., Wang, T., Michaelos, M. & Pachitariu, M. Cellpose: a generalist algorithm for cellular segmentation. Nature Methods 18, 100–106 (2021).

[111] Bravo González-Blas, C., De Winter, S., et al. SCENIC+: single-cell multiomic inference of enhancers and gene regulatory networks. Nature Methods 20, 1355–1367 (2023).

[112] Persad, S., Choo, Z.-N., Dien, C. et al. SEACells infers transcriptional and epigenomic cellular states from single-cell genomics data. Nature Biotechnology 41, 1746–1757 (2023).

[113] Ashuach, T., Gabitto, M. I., Koodli, R. V. et al. MultiVI: deep generative model for the integration of multimodal data. Nature Methods 20, 1222–1231 (2023).

[114] Bravo González-Blas, C., et al. cisTopic: cis-regulatory topic modeling on single-cell ATAC-seq data. Nature Methods 16, 397–400 (2019).

[115] Bredikhin, D., Kats, I. & Stegle, O. MUON: multimodal omics analysis framework. Genome Biology 23, 42 (2022).

[116] Ochoa, D., Hercules, A., Carmona, M. et al. The next-generation Open Targets platform: reimagined, redesigned, rebuilt. Nucleic Acids Research 51, D1353–D1359 (2023).

[117] Karczewski, K. J., Francioli, L. C., Tiao, G. et al. The mutational constraint spectrum quantified from variation in 141,456 humans. Nature 581, 434–443 (2020).

[118] Danecek, P., Bonfield, J. K., Liddle, J. et al. Twelve years of SAMtools and BCFtools. GigaScience 10, giab008 (2021).

[119] Dobin, A., Davis, C. A., Schlesinger, F. et al. STAR: ultrafast universal RNA-seq aligner. Bioinformatics 29, 15–21 (2013).

[120] Li, H. Aligning sequence reads, clone sequences and assembly contigs with BWA-MEM. arXiv preprint arXiv:1303.3997 (2013).

[121] Liu, B. B., Jessa, S., Kim, S. H. et al. Multiomics and deep learning dissect regulatory syntax in human development. Nature (2026).

[122] Zhang, Y., Liu, T., Meyer, C. A. et al. Model-based analysis of ChIP-Seq (MACS). Genome Biology 9, R137 (2008).

[123] Quinlan, A. R. & Hall, I. M. BEDTools: a flexible suite of utilities for comparing genomic features. Bioinformatics 26, 841–842 (2010).

[124] Shrikumar, A., Tian, K., Avsec, et al. Technical note on transcription factor motif discovery from importance scores (TF-MoDISco). arXiv preprint arXiv:1811.00416 (2018).

[125] Gupta, S., Stamatoyannopoulos, J. A., Bailey, T. L. & Noble, W. S. Quantifying similarity between motifs. Genome Biology 8, R24 (2007).

[126] Sundararajan, M., Taly, A. & Yan, Q. Axiomatic attribution for deep networks. Proceedings of the 34th International Conference on Machine Learning 70, 3319–3328 (2017).

[127] Kaistha, A. et al. Premature cell senescence promotes vascular smooth muscle cell phenotypic modulation. Cardiovascular Research (2025).

[128] Muzellec, B., Telenczuk, M., Cabeli, V. & Andreux, M. PyDESeq2: a python package for bulk RNA-seq differential expression analysis. Bioinformatics 39, btad547 (2023).

[129] Kolberg, L. et al. g:Profiler—interoperable web service for functional enrichment analysis and gene identifier mapping (2023 update). Nucleic Acids Research 51, W207–W212 (2023).

